# An investigation into the patterns of occupancy and detection of two endemic montane bird species, in Southern Western Ghats, India

**DOI:** 10.1101/2024.04.17.589990

**Authors:** Paul Pop

## Abstract

*Montecincla fairbanki* (Palani Laughingthrush) and *Sholicola albiventris* (White-bellied Sholakili) are avian species found only in a few high-altitude sky islands of the Southern Western Ghats. A taxonomic revision has resulted in the need for better resolution of species biology, especially their distribution, and factors affecting their distribution. The current study was an attempt to address this, by repeated surveys at plantation sites in the northern landscape of Idukki district (Kerala), and occupancy modelling. Around 96% of the sites surveyed were occupied by *M. fairbanki* and ∼52% by *S. albiventris*. Model-averaged site-averaged occupancy probability (ψ) of *M. fairbanki* is 0.8618 (0.6659-0.9229), and of *S. albiventris*, it is 0.4543 (0.1358-0.9972). Model-, site-, and survey-averaged detection probability (p) of *M. fairbanki* is 0.6554 (0.5171-0.7341), and of *S. albiventris*, it is 0.5731 (0.4102-0.7939). This confirms the hypothesis that *S. albiventris* has lower occupancy and detection probability compared to *M. fairbanki*. There is likely no effect on ψ of one species by the presence of the other, but there is likely a positive effect on p of one species, by the occupancy or detection of the other. Most significant effects on ψ of *M. fairbanki* were from the exclusive presence of dirt roads, elevation, slope, and visibility, and for *S. albiventris*, it was from *Lantana camara*, elevation, and understorey height. Most significant effects on p of *M. fairbanki* were from windy weather, sky conditions, time of surveys, and survey number, whereas for *S. albiventris*, they were from neutral weather, and survey number. Of the interactive effects hypothesized, significant evidence was found for the increase of ψ as a function of elevation, and the reduction of ψ in the presence of the invasive *L. camara* (only for *S. albiventris*). The insights gained from this baseline study will be useful for further hypotheses testing and conservation efforts.

## 1. INTRODUCTION

Endemism of birds is common in mountainous regions and more pronounced in the tropics, where interplay between ecological isolation and climate of lowland from montane habitats may act as a barrier for dispersal of several high-altitude species to adjacent areas (Sekercioglu et al., 2008). The term ‘sky islands’ is used for such regions (Warshall, 1995), and Western Ghats is a host to several sky islands. Two avian genera endemic to the Western Ghats are *Sholicola* (sholakilis/blue robins) and *Montecincla* (laughingthrushes/chilappans), which are restricted to the *shola* sky island complex. Long-term research on these taxa has culminated in their taxonomic division into seven different species (Robin et al., 2017), six of which are currently accepted (Collar et al., 2020; Praveen & Jayapal, 2024). Of these, *Sholicola albiventris* (White-bellied Sholakili) and *Montecincla fairbanki* (Palani Laughingthrush) are found in the Anaimalai-Palani hills complex, located on the southern side of the Palghat gap - the widest biogeographical gap in Western Ghats (Robin et al., 2017). These are the species considered for this study (Figure 1).

**FIGURE 1.**
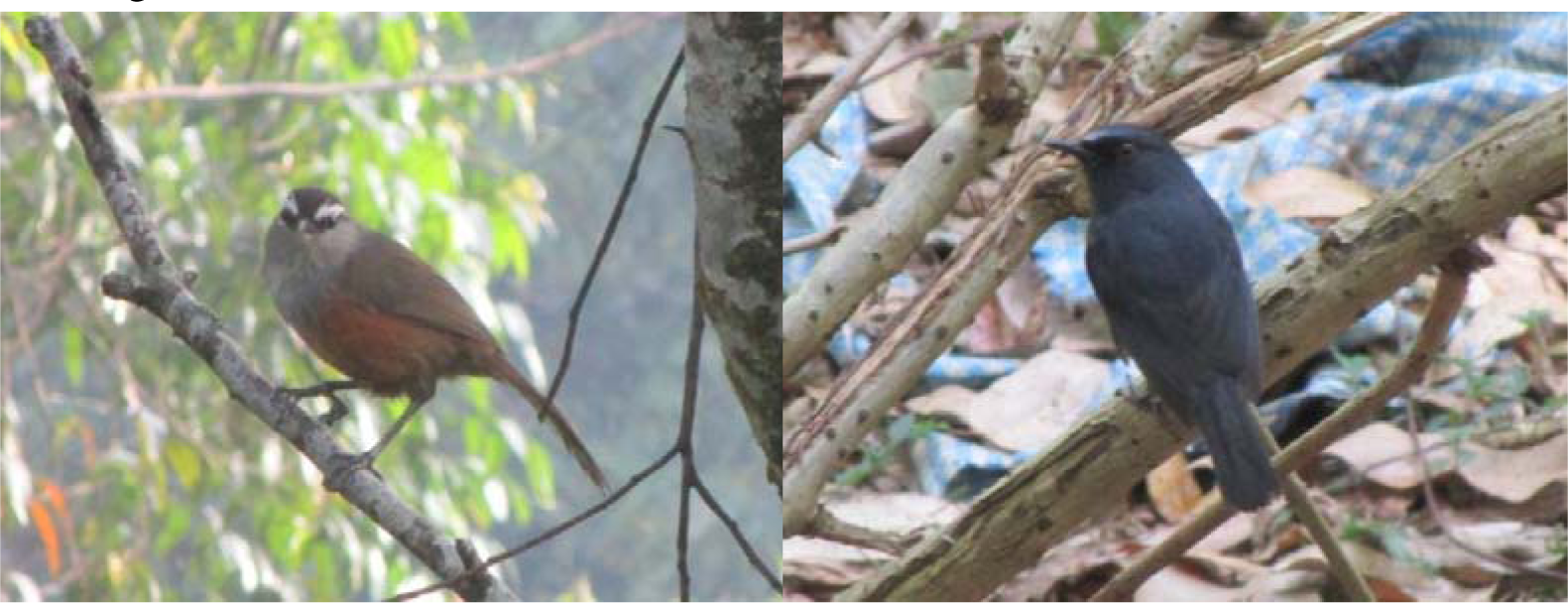
Study species: *Montecincla fairbanki* (left) and *Sholicola albiventris* (right).

Occupancy surveys are part of a sampling method involving multiple visits to selected sites during a suitable time-period when species detection is possible, for which a variety of sampling methods exist (Mackenzie et al., 2002). Non-detection is a crucial parameter which gives more meaning to the proportion of sites occupied by species and predicts the actual occupancy of the species in the study area. (Mackenzie et al., 2002). Sites in this detection–non-detection framework is analogous to individuals in a capture-recapture model and this model allows for multiple techniques, observers, seasons/time-periods, missing information to be incorporated into the design (Mackenzie et al., 2002). One of the main themes of occupancy studies is to look at how detection probability can affect the estimation of the distribution/occupancy of one or more species (eg: De Melo et al., 2022).

The aim of the study was to determine the occupancy status and detection probabilities of *M. fairbanki* and *S. albiventris* in the Idukki landscape, and determine alongside, the factors influencing these traits. The need for the study rests on the fact that knowledge gaps exists in literature about endemic avian species in Western Ghats, including occupancy status (Bawa, 2007; Robin & Nandini, 2012).

Exacerbating this situation is the somewhat recent taxonomic revision, which is a cause for data deficiency for individual species of these species complexes. Threat status, distribution, biology and other aspects of these different species are not or improperly known, and needs reassessment.

Based on the knowledge available prior to the study, I hypothesized that *S. albiventris* will have lesser occupancy and lower detection probability compared to *M. fairbanki* which occupy the same landscape. Since there was little knowledge about the factors affecting the occupancy probability (ψ) and detection probability (p) of these species, prior to the study, only a few hypotheses regarding them were made: i) The presence of the invasive species *Lantana camara* will reduce the ψ of both the species. ii) The presence of (running or standing) water will increase the ψ of both the species iii) The ψ will increase as a function of increasing elevation iv) p of both the species will be higher during cooler weather (given that the study was carried out in the summer).

## 2. MATERIAL AND METHODS

### 2.1 Study area and site selection

This study was carried out in the Devikulam taluk — northernmost portion of Idukki district in Kerala, India (Figure 2 & 3). Idukki is the second largest district in the state of Kerala, but has the lowest human population density in the state, with a population of 1,108,974 as of 2011 (ORGI, 2011).

**FIGURE 2.**
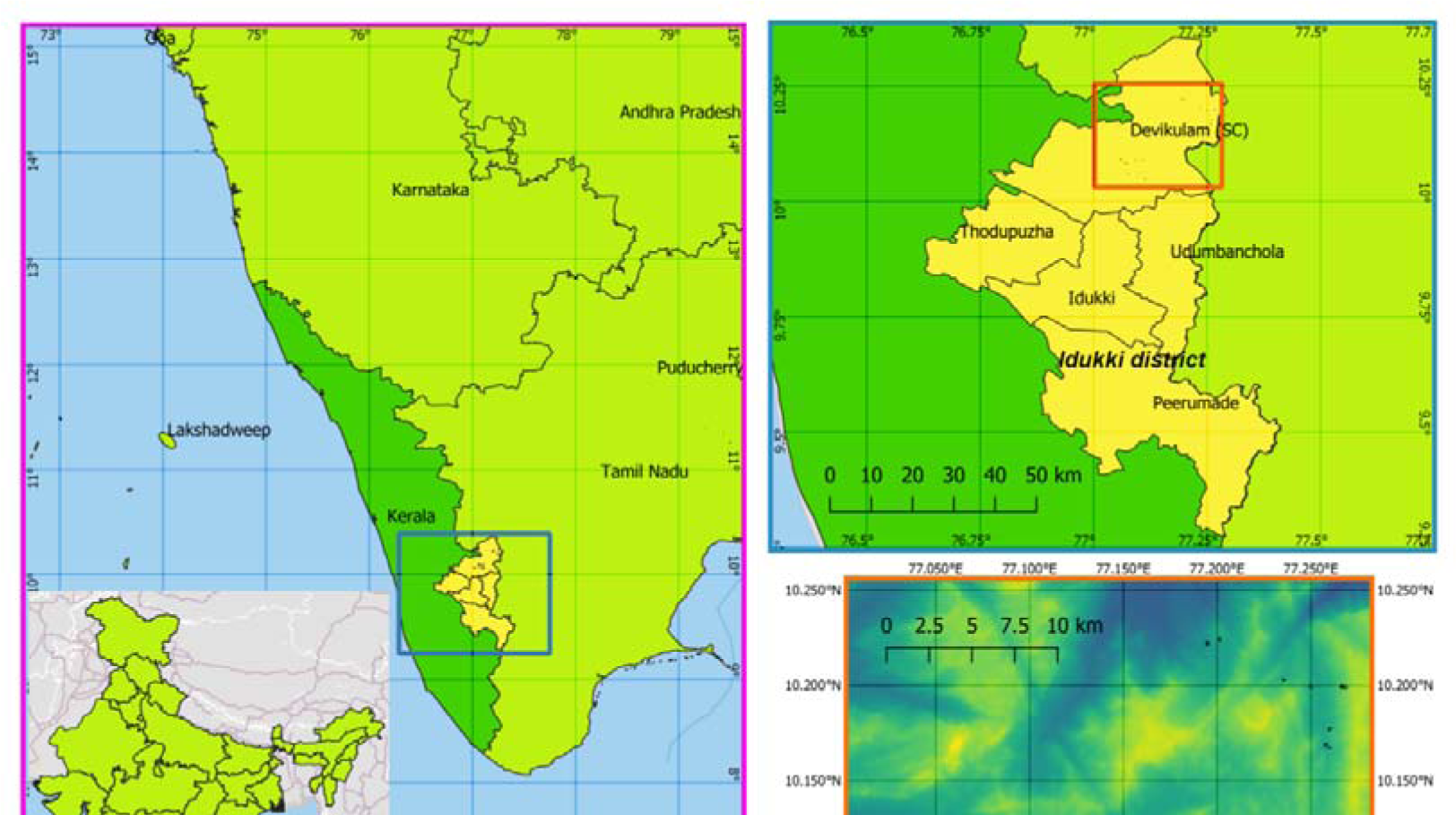
Study area map (note that the India outline is based on the boundaries claimed by the Govt. of India).

For site selection, Arasumani et al. (2018) was followed and GIS data was obtained from the authors of the paper based on the published methods. A random systematic sampling approach was followed for site selection. 10% of the plots generated were considered for surveys (112 plantation plots and 80 *shola* patches), a subset of which were the (mostly) *Eucalyptus* plantations which could be actually surveyed within the study period (n = 27; see Figure 3).

**FIGURE 3.**
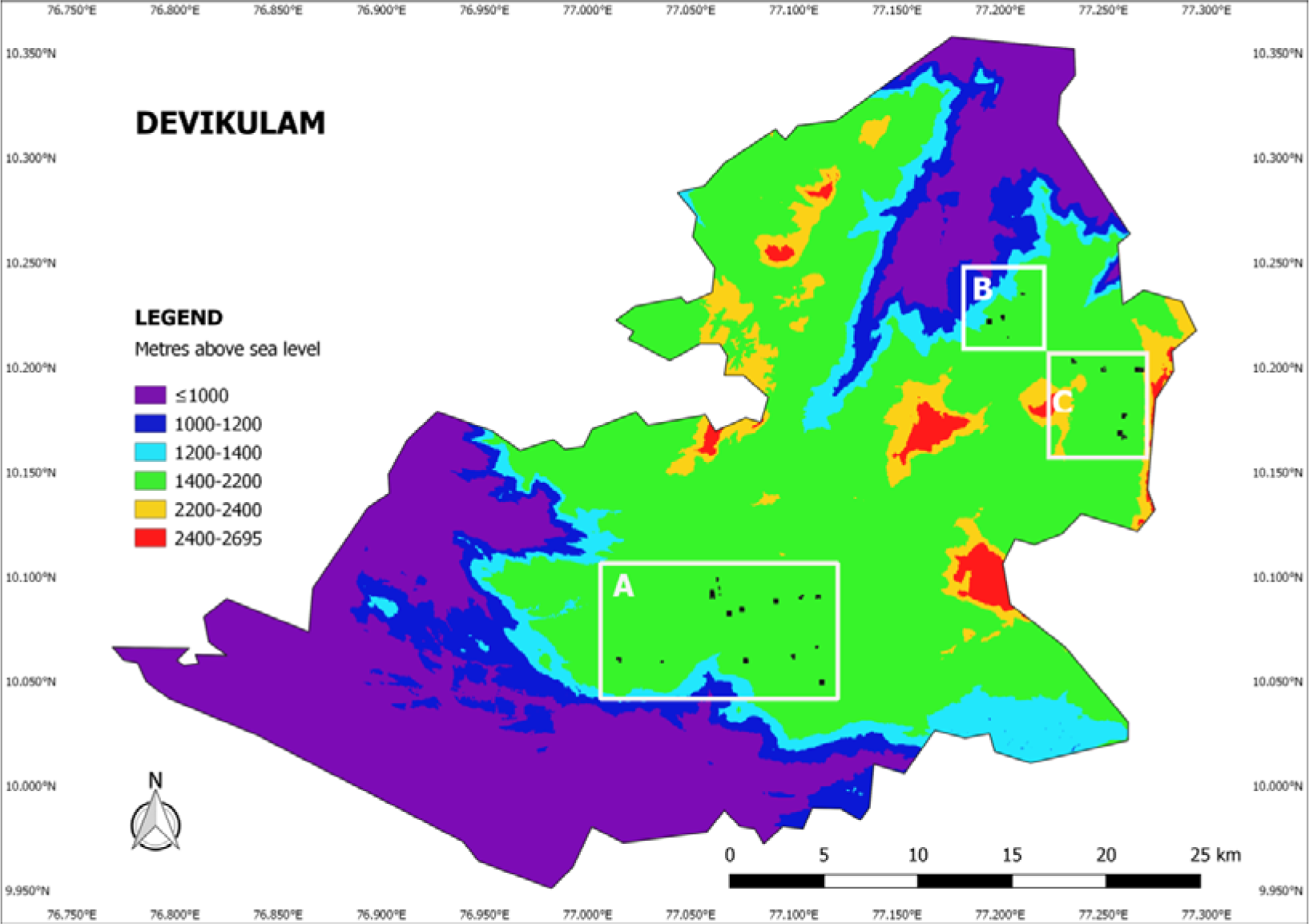
Study area map showing the survey sites and clusters.

The study was centered around three main clusters: *A*) Munnar *B*) Kanthaloor and *C*) Koviloor (Supporting information shows landscape images). *A* is mostly covered by tea estates, whereas *B* and *C* are occupied by *Eucalyptus* plantations and terrace-farms, with no tea gardens. All three places are tourist destinations with decreasing order of visits from *A* to *C*. Of the 27 plots surveyed, 16 were in *A*, four in *B*, and seven in *C*. Plot size varied from 0.09 to 5.0625 ha, with the majority of them larger than 2 ha in size. The highest plots sampled in *A*, *B* and *C* were at ∼ 1645 m a.s.l., 1571 m a.s.l. & 2132 m a.s.l.; whereas the lowest was at ∼1472 m a.s.l., 1457 m a.s.l., and 1652 m a.s.l.; respectively, showing the difference in elevational ranges.

All the three clusters’ climate is classified as warm and temperate. There’s more rain in summers compared to winters. The Köppen-Geiger climate classification is Cwb which means it has a mild temperature with dry winter and warm summer (Chen & Chen, 2013). It was summer when the surveys were carried out in all 3 clusters, with the average temperature being ∼20° C. There were summer rains during the study period in all three clusters, with little to moderate precipitation in March and high precipitation in April.

### 2.2 Field methods

The study duration in Idukki was from 5^th^ March to 17^th^ April, 2018. The author initiated surveys only after gaining sufficient confidence that the two species could be identified, either by call or sight without confusion (with training at Bombay shola, Kodaikanal, Dindigul district, Tamil Nadu). Mock surveys were carried out at several plots near Ooty (Nilgiris district, Tamil Nadu). Employment and standardization of various covariate estimation was done in this location, especially the modified staff-ball technique for horizontal cover estimation (Pop, 2019).

Four temporal replicates (surveys) were carried out in each plot: two surveys before and two after noon, with only two exceptions (assuming little to no effect on data collected since these two surveys fall quite close to the required time splits (AM/PM)). Each survey in the same plot was separated by a gap of at least (and almost always) one day so as to increase detection but by maintaining it consistent across the plots. Six minutes were spent per hectare and the differently sized plots were surveyed on the basis of this time:area ratio. Start and end time were noted for each survey. Detection of *S. albiventris* and *M. fairbanki* within this duration was noted. Surveys were avoided when there was precipitation, as commonly practiced in occupancy surveys (West et al., 2016; Shahan et al., 2017).

Distance sampling which “is an extension of plot sampling, in which birds are counted within a sample of defined areas (plots)” (Buckland, 2001) used for occupancy surveys of many species was not possible, as the terrain inside the randomly generated plots were previously unknown and unmarked; all parts inside a plot were not always accessible; time and energy for designing and maintaining transects would put a massive observer cost; and point counts for differently sized plots would bias the detection highly in favour of smaller plots. Therefore the surveys were conducted along animal trails/cleared paths across the plots, diagonally whenever possible, so as to cover the longest distance. Species observations by vocalisations were only recorded when absolutely confident positive identification was possible (to prevent false positives). All birds (besides the focal species) heard or seen within the survey were also noted down and checklists of all surveys were uploaded in eBird platform to aid further research.

Weather-associated parameters were written down for each survey — weather - “sunny” & “windy”; skies - “over-cast/misty”, “clear skies/calm-cloudy”; and felt temperature - “warm”, “neutral” and “cool”. Stage of plantation (“young” and “mature”) was recorded. Interesting vegetation found in the plots such as invasive species (for example, *Lantana camara*, *Parthenium hysterophorus*, *Chromolaena odorata* and *Ageratina adenophora*) and food plants of the species under study (e.g., *Rubus niveus*) were noted. Accessibility was divided into “no-road” (animal trail/cleared path), “dirt/foot road” and “metalled road”. Presence or absence of waterbody (lentic or lotic) and burn were also recorded. Waterbodies falling within 5 m from the boundary of plots were also considered, accounting for movement of the birds. Lotic (flowing water – rivers or streams) was noted as either dry or wet/moist. Lentic (standing water) included temporary pools, even found within a dry lotic system, ponds, and water tanks. Survey number was used as another covariate.

Vegetation parameters collected were of three types: understorey height, canopy height and horizontal cover/visibility. The first two were measured by visual estimation whereas horizontal cover was measured using a modified Staff-ball technique (Pop, 2019). Five measurements each of the former two were taken for every hectare, whereas three measurements of horizontal cover were taken per hectare. The higher number of spatial replicates for these quantities are to iron out outliers and provide a meaningful average for analysis.

### 2.3 Analysis

Data analysis was primarily carried out using the package RPresence v. 2.13.60 in R v. 4.4.0 (See Supplementary Code for full details) with help from documentation in occupancyTuts (Donovan et al., 2024). Both site-specific and survey-specific variables, either continuous or categorical were used, which when combined, constituted 26 covariates (see Supporting information). Correlations between continuous covariates were checked (see Supporting information). Single-season single-species model type was used for both species separately to see the factors affecting occupancy and detectability at individual resolution, and single-season species co-occurrence model type was employed for combined analysis of both species.

Most covariates were from field data, whereas some of them - aspect, elevation, and slope were all extracted from the ASTER GDEM layer obtained from USGS (ASTER GDEM is a product of METI and NASA) in QGIS. In QGIS v 1.8.0., the shapefiles of polygons encompassing the 27 study sites were over-layed over ASTER GDEM. DEM (Terrain Models) under “Analysis” were used to derive the slope and aspect layers, whereas contour extraction was carried out for deriving the Altitude. Zonal Statistics plugin was used for averaging of all three covariates within these polygons.

In the co-occurrence model of RPresence, default parameterisation and compressed inputs (coded) were used. Continuous variables were transformed into standardized form (Z values = (raw value – mean value)/standard deviation) with average value taken as zero and values generally lying in between +3 to -3. This standardizing helps in easier analysis of data, since it reduces the chances of optimisation algorithm failure in finding correct parameter estimates when met with large or small values of mean (Donovan et al., 2024). The number of covariates (26) was arrived at, after the elimination of some of the potential covariates such as presence of some invasive species - *Chromolaena odorata*, *Ageratina adenophora* and *Parthenium hysterophorus* since the first two were found in all sites except one and the last was found in only two sites, none of which allows for successful modelling. The covariates of maturity of plantations were not included in the analysis since all except one patch was mostly mature.

Theoretically, there are an infinite number of models that could be created from the data. So, focus was given on testing some hypotheses and narrowing down the number of covariates/variables, i.e., a multi-stage approach was used (Shahan et al., 2017). Models were hypothesized based on probable interactions of covariates on the occupancy and detection probabilities, and ones that did not make any such sense, were ignored. The number of covariates added for psi and p were based on constraints posed by the sample size. Observing this constraint, the number of covariates which were found to be significant for further testing were reduced drastically, by forward elimination. Significant models were found using corrected Akaike Information Criterion (AIC) values, under the Model Selection approach, which has gained popularity over other methods such as t-tests (coefficients), F-tests (ANOVA), adjusted *R^2^* and Likelihood Ratio Test (LRT) (Burnham & Anderson, 2002; Johnson & Omland, 2004).

QAICc of each model was used, instead of just AIC, to avoid the effect of low sample size on model selection as described for avian species in Mexico (Stockwell & Peterson, 2002) and Finland (Pakanen et al., 2018), as well as to adjust for the observed underestimation of variances in models. The ‘Q’ in QAICc denotes the quasiliklihood adjustment of AIC, and the ‘c’ denotes a small sample correction added to the QAIC, which gives a big penalty for small sample sizes. Models having ΔQAICc (Difference between the AICc value of the current model and model with the lowest QAICc) values between 0-2 are considered the best models and between 2-7, good models.

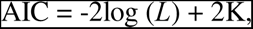

where *L* is the maximized log-likelihood/deviance and K is the correction for the asymptotic bias/number of estimable parameters in any given model (Burnham et al., 2011).

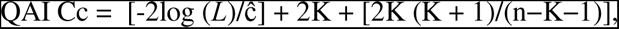

where n is sample size (Burnham et al., 2011) and ĉ is the quasilikelihood variance inflation factor (calculated by χ2GOF/df i.e. Chi-squared Goodness of fit test/degrees of freedom).

The most general set (containing the most number of parameters) in the single species model sets (for *M. fairbanki* and *S. albiventris*) were bootstrapped (50,000 iterations each) and tested for model fit (MacKenzie and Bailey goodness of fit test) so as to check the reliability of the models in the model set (Donovan et al., 2024). In these models, ‘ψ’ (psi) is the probability of occupancy and ‘p’ or ‘r’ is the detection probability. The strength and direction of effects of covariates were evaluated based on several factors: The strength of effects of covariates were evaluated based on these factors (see Supplementary Code section 4 for details):

i) Magnitude of standard error (SE) in relation to the estimate of betas (considerable significance for estimate > SE, and more for smaller SE)
ii) Range of confidence interval of betas (considerable significance when Confidence Interval (CI) doesn’t pass through 0, and more for smaller range of CI)
iii) The number of models they appear in the final mode set (a higher number indicate higher significance)
iv) The sums of weights of models in which they appear (a higher number indicate higher significance)
v) Plotting of the psi and p estimates against the covariates (expert evaluation of the pattern to assess magnitude of influence of covariate on psi/p).
vi) Spearman’s/Kendall’s rank correlation (interpretation can be found in Supplementary Code section 5.3).

Based on these, the strength of effects of covariates were classified into ‘very strong’, ‘strong’, ‘moderate’, ‘low’, ‘very low’, and ‘feeble’, in the decreasing order of significance.

The direction of effects of covariates were evaluated based on these factors:

i) Betas: negative betas indicate negative effect, and positive betas - positive effect.
ii) Sign of upper and lower limits of confidence intervals - if both are positive, then positive effect; if both are negative, then negative effect; and if they are opposite signs, it is likely inconclusive.
iii) Plotting of the psi and p estimates against the covariates (the direction of effect is sometimes apparent).
iv) Spearman’s/Kendall’s rank correlation - signs indicate direction.

Besides the occupancy and detection probability modelling, post-hoc simulations using the genpresEV function in RPresence was carried out to find the optimum survey design given the ψ and p values found from the study, so as to have higher precision of estimates, with the least expenditure of money and time (see Supplementary Code section 6 for more details).

## 3. RESULTS

From the field data collected from different sites (Supplementary Data 1 & 2), including observations taken outside of surveys (which has been submitted to eBird), it was found that ∼ 96.29% of the sites studied were occupied by *M. fairbanki* and ∼ 51.85% of sites studied were occupied by *S. albiventris*, indicating widespread distribution of the former in the study area and sparser distribution of the latter. Since the sites were surveyed four times each, it is unlikely that the species were present but undetected at all times. Cluster-wise, the occupancy status was as shown below (the numbers should be seen in the light of different sample sizes for each cluster): *A*) Munnar (and parts of Devikulam): 100% occupancy of *M. fairbanki* and only 37.50% occupancy of *S. albiventris B*) Kanthaloor: with only 75% occupancy of *M. fairbanki* and 25% occupancy of *S. albiventris*, the numbers were the lowest of the all the three clusters *C*) Koviloor (with Vattavada-Kottakamboor): 100% occupancy of both species.

While all types of models were tested, almost all of the models that were found to be significant were simple or additive linear models (including categorical models). In addition, only one polynomial linear model was present in the final model sets. Linear models with interactive effects were not present due to higher penalties from the higher number of parameters.

### 3.1 Significant models of single-season single-species analyses of *M. fairbanki*

Seventy-eight significant models (with a good possibility for more) are suitable for explaining the occupancy and detectability of *M. fairbanki*, out of which 14 are top models (see Supporting information). The top models for *M. fairbanki* include the covariates No road, Windy, Survey, Before noon, Dirt road, Slope, Elevation, and After noon.

Naïve occupancy estimate of *M. fairbanki* is 0.8519 (proportion of sites where the species were detected at least once). Here are the model-averaged (mod-avg) values for *M. fairbanki* (Figure 4, 5a; Supporting information): Mean of ψ = 0.8618, ii) Range of ψ = 0.6659-0.9229, iii) Range of SE of ψ = 0.0747-0.2889, iv) Range of 95% CI of ψ = 0.1359-0.9895, v) Mean of p = 0.6554, vi) Range of p = 0.5171-0.7341, vii) Range of SE of p = 0.0737-0.1774, viii) Range of 95% CI of p = 0.2101-0.8845.

**FIGURE 4.**
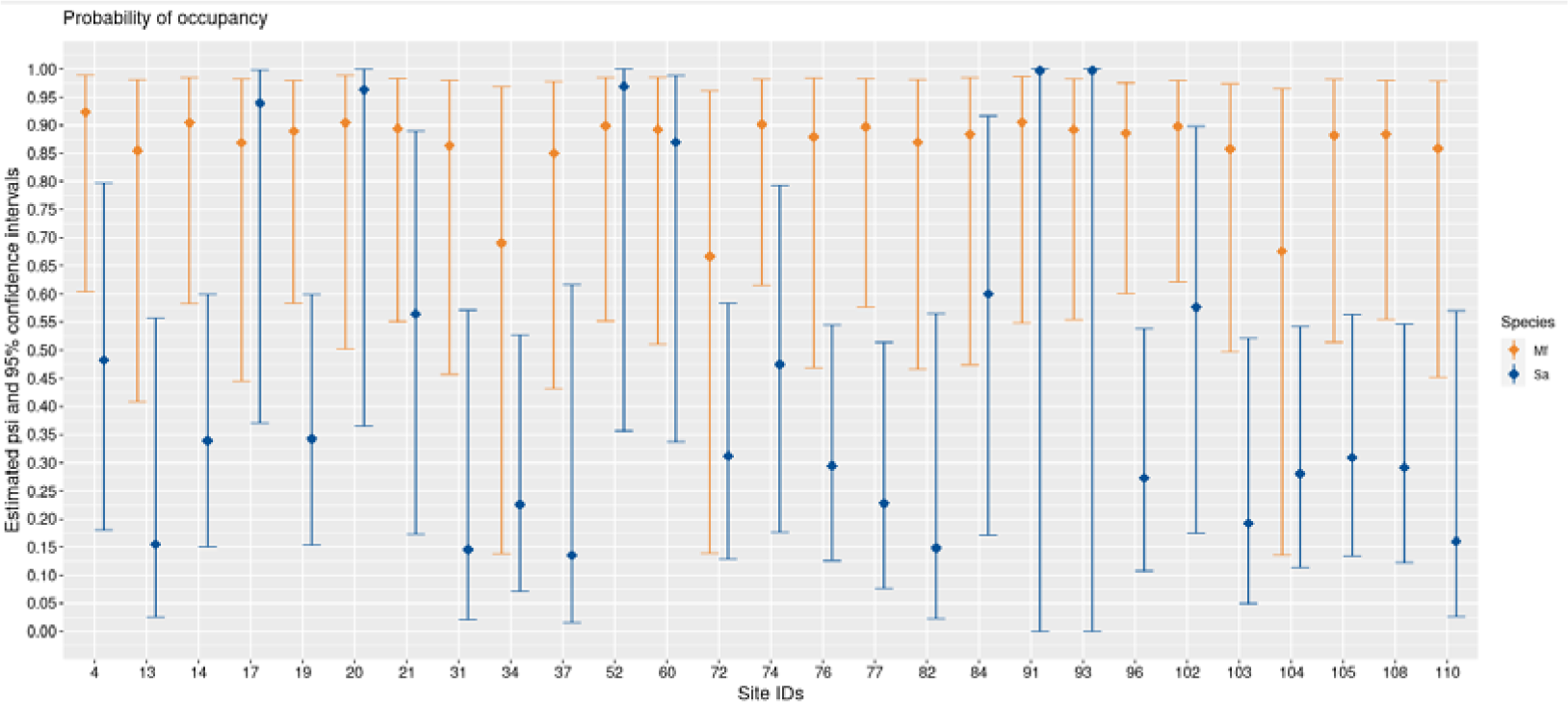
Sitewise probability of occupancy (ψ) of *M. fairbanki* (Mf) and *S. albiventris* (Sa).

**FIGURE 5.**
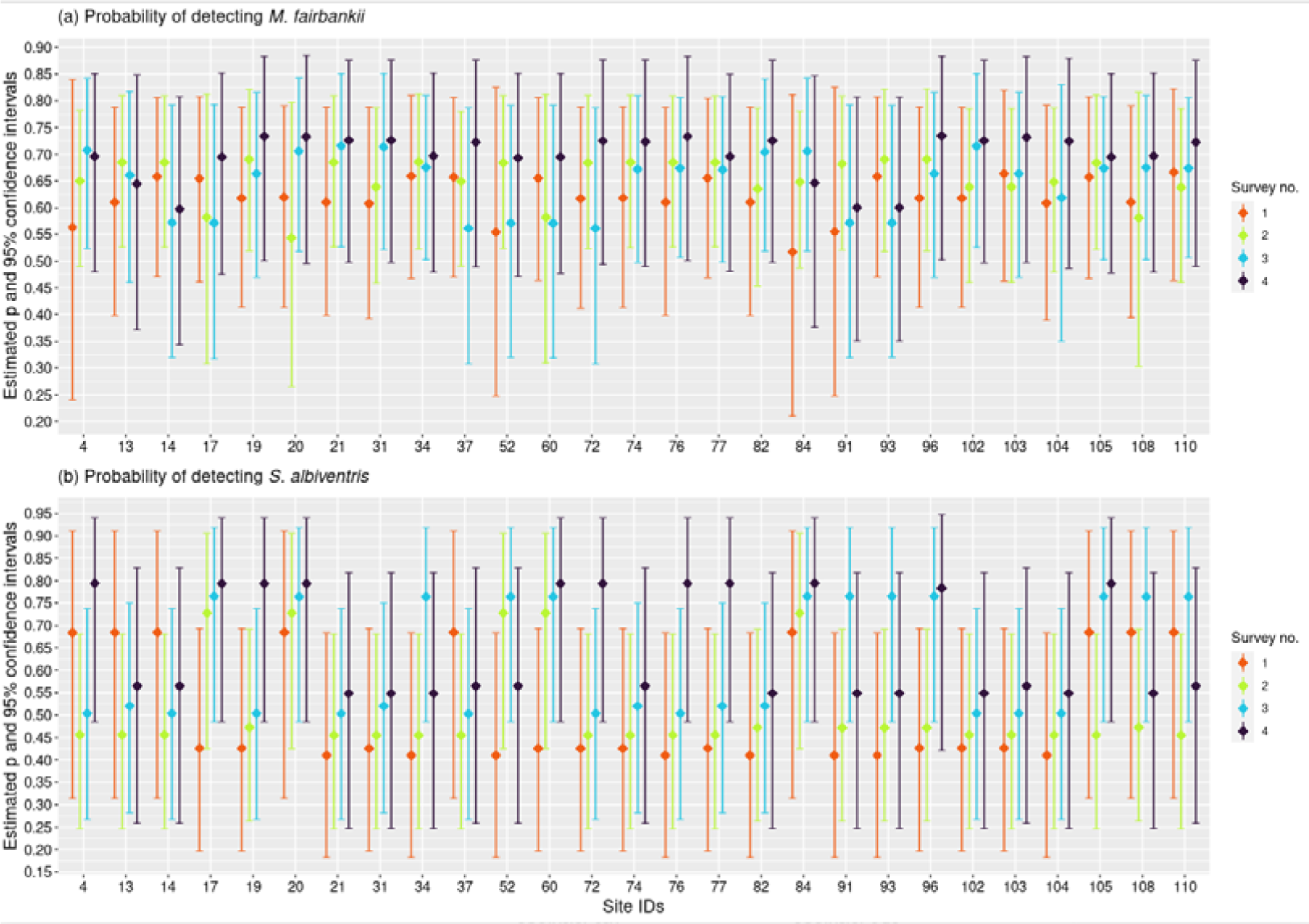
Site- and survey-wise probability of detection (p) of *M. fairbanki* and *S. albiventris*.

### 3.2 Significant models of single-season single-species analyses of *S. albiventris*

Twelve significant models (with very limited possibility for more) are suitable for explaining the occupancy and detectability of *S. albiventris*, out of which only three are top models (see Supporting information). The top models for *S. albiventris* include the covariates Elevation, Neutral, Survey, and *L. camara*.

Naïve occupancy estimate of *S. albiventris* is 0.4444. Here are the mod-avg values for *S. albiventris* (Figure 4, 5b; Supporting information): Mean of ψ = 0.4543, ii) Range of ψ = 0.1358-0.9972, iii) Range of SE of ψ = 0.0294-0.2431, iv) Range of 95% CI of ψ = 4.319817×10^-7^-1, v) Mean of p = 0.5731, vi) Range of p = 0.4102-0.7939, vii) Range of SE of p = 0.1138-0.1714, viii) Range of 95% CI of p = 0.1827-0.9473.

### 3.3 Significant models of single-season two-species analyses

Since none of the models in the set containing significant co-occurrence models had interactive effect on ψ, the species interaction factor/SIF (denoted by phi/φ) given by the formula, φ = ψ^M^ × ψ^MS^/(ψ^M^ × ψ^S^) (1), was found to be equal to 1, providing support to the null hypothesis that there is no significant interaction between the two species i.e there is no evidence to suggest species avoidance (e.g., Bailey et al., 2009) or species convergence (e.g., Richmond et al., 2010)). However, models containing interaction factors of occupancy and detection, on detection probability were found to be significant.

Since the SIF is zero, ψ of co-occurrence models doesn’t give any more information than is already known from the single species models. So, only the results of detection probabilities are shown for two-species models.

#### 3.3.1 MS format: Species A = *M. fairbanki*, Species B = *S. albiventris*

Ten significant models (with limited possibility for more) are suitable for explaining co-occurrence of two species, out of which only six are top models (see Supporting information). The top models include the covariates Elevation, Survey, and *L*. *camara*.

Here are the mod-avg values of r for co-occurrence (Figure S9a, S10a, S11a; Supporting information — table description contains explanation of notations used here): Mean of r^M^ = 0.6722, ii) Range of r^M^ = 0.5784-0.7258, iii) Range of SE of r^M^ = 0.0647-0.1223, iv) Range of 95% CI of r^M^ = 0.3393-0.8609, v) Mean of r^SM^ = 0.6369, vi) Range of r^SM^ = 0.5237-0.6991, vii) Range of SE of r^SM^ = 0.0882-0.1630, viii) Range of 95% CI of r^SM^ = 0.2339-0.8558, ix) Mean of r^Sm^ = 0.4858, x) Range of r^Sm^ = 0.2844-0.5778, xi) Range of SE of r^Sm^ = 0.1304-0.7193, xii) Range of 95% CI of r^Sm^ = 0.0004-0.9975.

#### 3.3.2 SM format: Species A = *S. albiventris*, Species B = *M. fairbanki*

Similar to the MS format, there are twelve significant models (with limited possibility for more), out of which only six are top models (see Supporting information). The top models include the same covariates Elevation, Survey, and *L*. *camara*. In fact, besides the swapping of interaction of occupancy on r with interaction of detection on r in two models, all the top six models are the same.

Here are the mod-avg values of r for co-occurrence (Figure S9b, S10b, S11b; Supporting information — table description contains explanation of notations used here): Mean of r^S^ = 0.6158, ii) Range of r^S^ = 0.5218-0.6637, iii) Range of SE of r^S^ = 0.0786-0.1095, iv) Range of 95% CI of r^S^ = 0.3161-0.8282, v) Mean of r^MS^ = 0.6941, vi) Range of r^MS^ = 0.6087-0.7459, vii) Range of SE of r^MS^ = 0.0683-0.1184, viii) Range of 95% CI of r^MS^ = 0.3699-0.8761, ix) Mean of r^Ms^ = 0.6170, x) Range of r^Ms^ = 0.3658-0.7158, xi) Range of SE of r^Ms^ = 0.1049-0.7456, xii) Range of 95% CI of r^Ms^ = 0.0011-0.9967.

### 3.4 Effects of covariates

Several covariates were found to consistently occupy models with low ΔQAICc in all model types.

#### 3.4.1 Covariates influencing the ψ and p of *M. fairbanki*

The strength and direction of influence of covariates on ψ are as follows (Figure 6, Supporting information): ‘No road’ has a strong negative influence. In other words, presence of roads have a positive influence. Since the covariate Dirt/foot road shows moderate to strong positive influence on ψ, and Metalled road shows very low positive influence, the type of roads increasing ψ are Dirt/foot roads. Elevation and slope also have a moderate to strong influence (the former positive and the latter negative). Visibility show a low to moderate positive influence.

**FIGURE 6.**
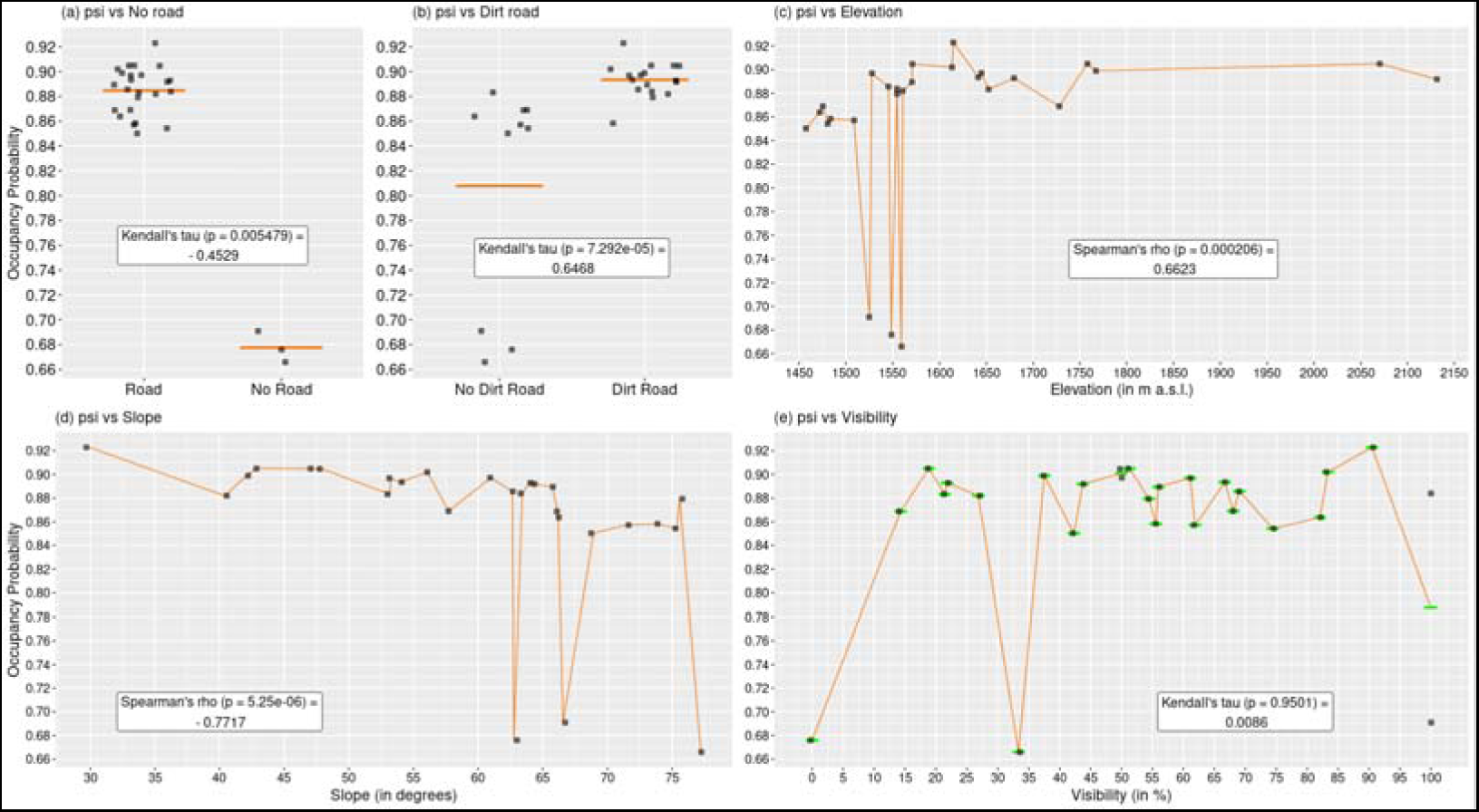
Covariates affecting occupancy of *M. fairbanki*. Jitters are datapoints. Horizontal lines in the plots of binary covariates indicate mean.

Other covariates appearing in the good models for *M. fairbanki* have low to feeble influence: Area and Understorey height have low influence, whereas Burn, Canopy height, Aspect, Lentic and Wet lotic waterbodies, and *R. niveus* have very low influence. Dry lotic shows feeble influence.

The strength and direction of influence of covariates on p are as follows (Figure 7, Figure 8a, Supporting information): The covariate Windy has a strong negative influence; Survey number and Before noon have moderate positive influence; After noon and Clear skies have moderate negative influence; and Overcast skies show low to moderate positive influence. Other covariates appearing in good models have low (Cool), or very low (Visibility, Warm, and Neutral) influence on p.

**FIGURE 7.**
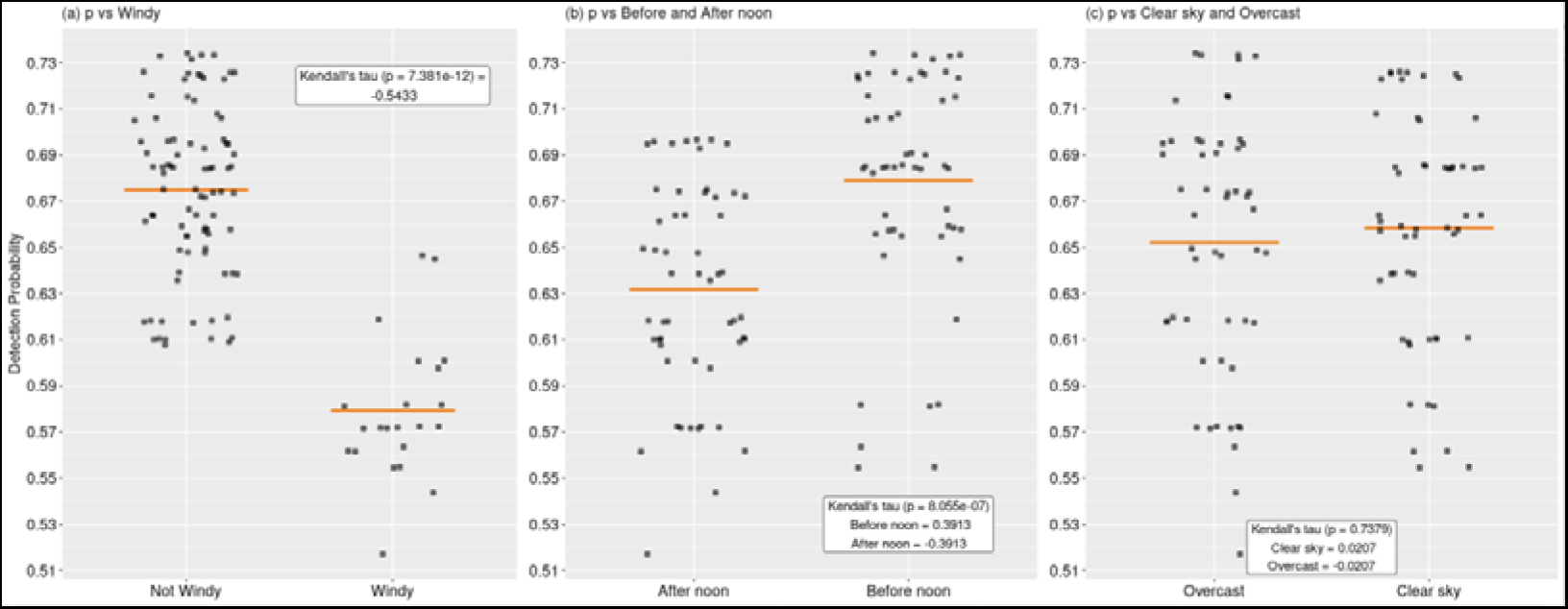
Covariates affecting detection of *M. fairbanki*.

**FIGURE 8.**
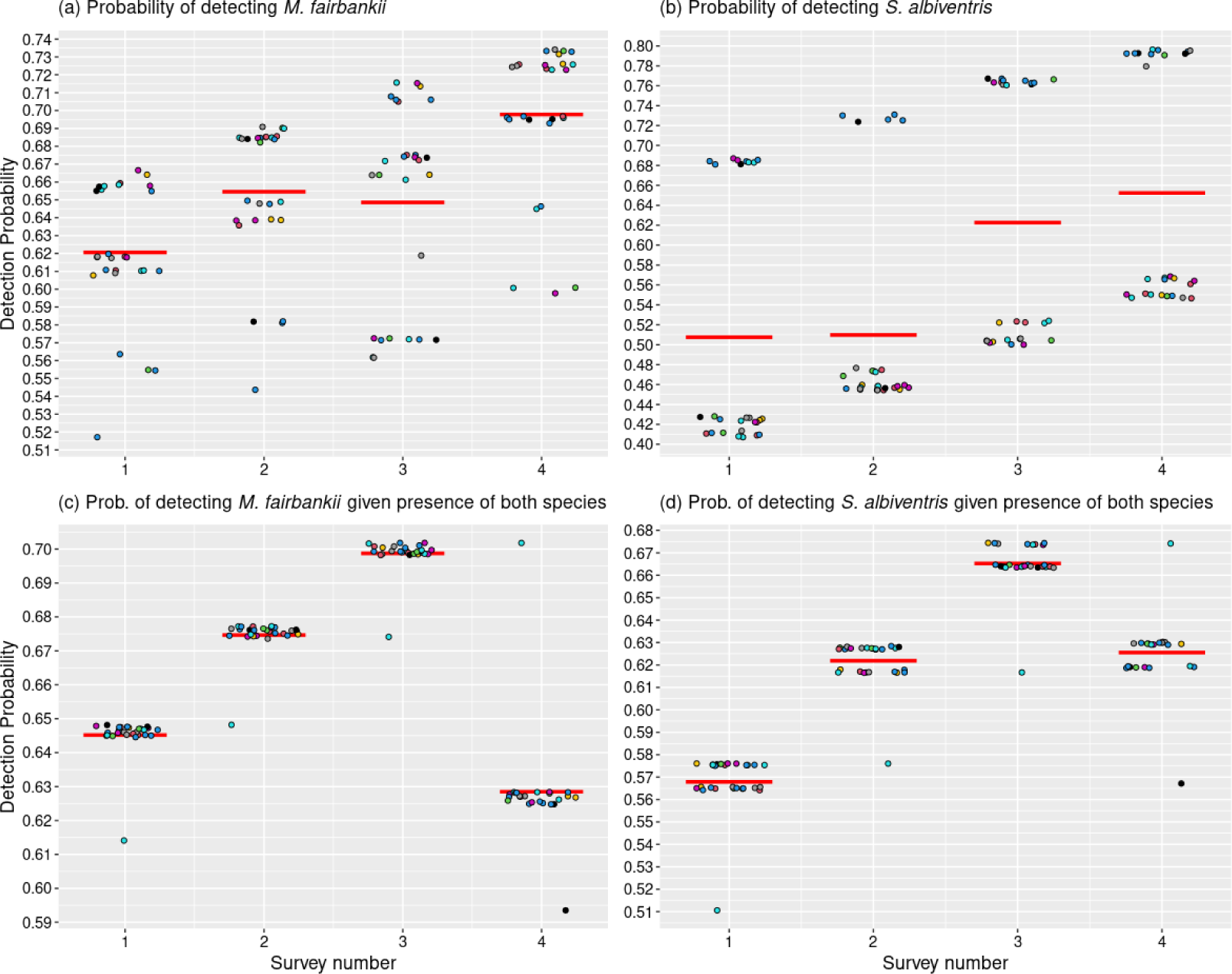
a) Unconditional and b) conditional survey-wise detection probability of *M. fairbanki* and *S. albiventris*. Colour of the jitters indicate site IDs, and the red line is the mean.

#### 3.4.2 Covariates influencing the ψ and p of *S. albiventris*

The strength and direction of influence of covariates on ψ are as follows (Figure 9a,b,c; Supporting information): *L. camara* shows a very strong negative influence; Elevation shows a strong positive influence; and Understorey height shows a moderate to strong negative influence.

**FIGURE 9.**
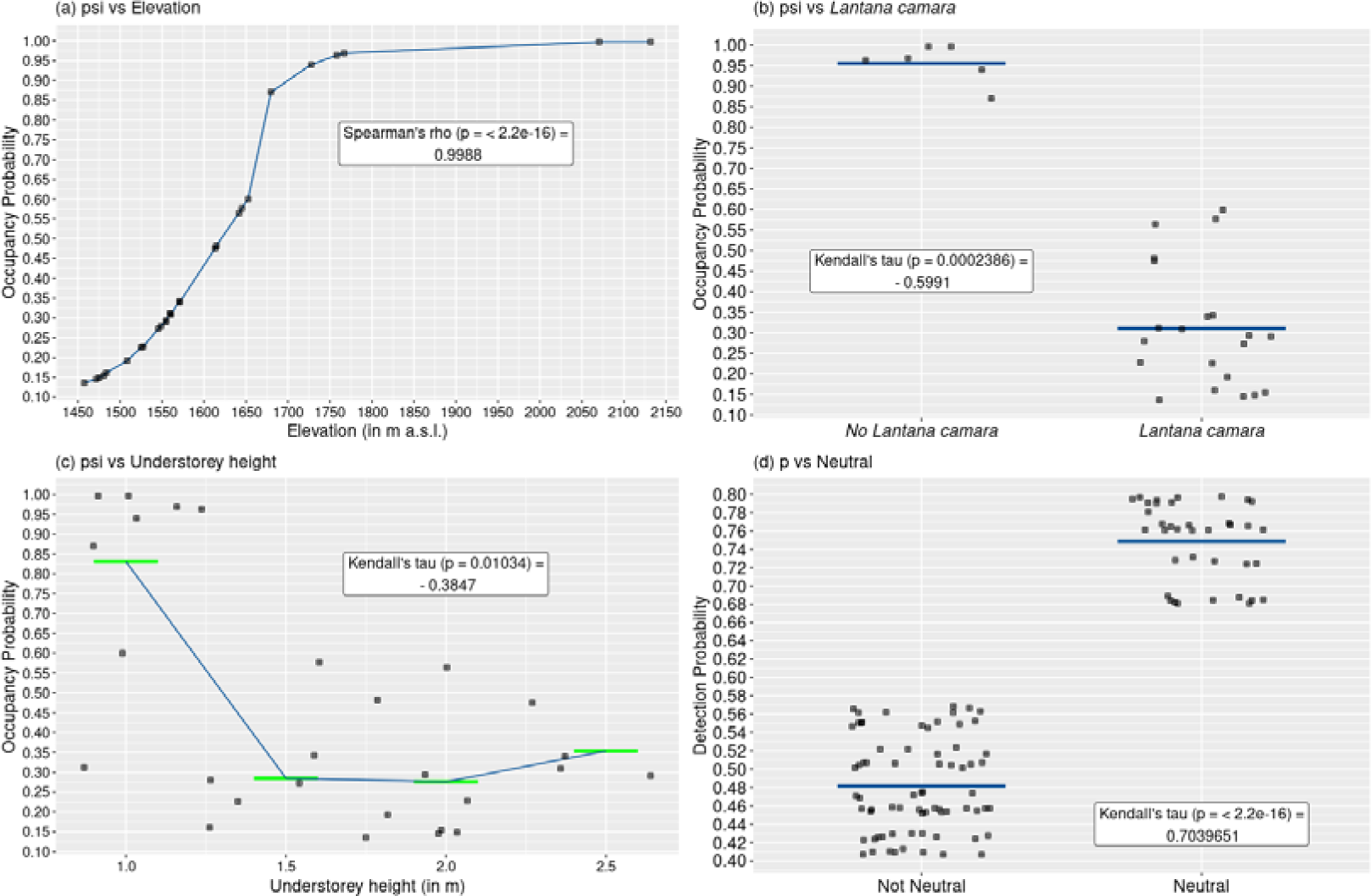
Covariates affecting occupancy and detection of *S. albiventris*.

The strength and direction of influence of covariates on p are as follows (Supporting information): ’Neutral’ weather shows a strong positive influence (Figure 9d); Survey number shows a moderate positive influence (Figure 8b); whereas Warm shows low and Cool shows very low influence.

#### 3.4.1 Covariates influencing the ψ and p of Co-occurrence models

The covariates and their influence on both the MS and SM format in co-occurrence format is nearly the same (Supporting information). The strength and direction of influence of covariates on ψ are as follows (Supporting information): *L. camara* shows a very strong negative influence; Elevation shows a very strong positive influence; ‘No road’ shows a moderate to strong positive influence; and Area shows a moderate to strong positive influence. The covariate Before noon show a low certainty of influence on r (Supporting information).

### 3.5 Post-hoc analyses

Simulation of precision values for the mean mod-avg ψ and p for both species indicate that there is noticeable increase in precision (decrease in standard error and confidence interval) when doing 3 surveys instead of 2, but not much when choosing 4 surveys instead of 3 (Figure 10, 11). The number of sites depended on what level of precision is aimed for. While the precision can be very slightly increased for ψ of both species with slightly lesser number of surveys i.e. 105 (S = 35, K = 3) for *M. fairbanki* and 93 (S = 31, K = 3) for *S. albiventris*, a very small increase in precision of p requires more number of surveys than the current study i.e. 116 (S = 29, K = 4) for both species (see Supporting information).

**FIGURE 10.**
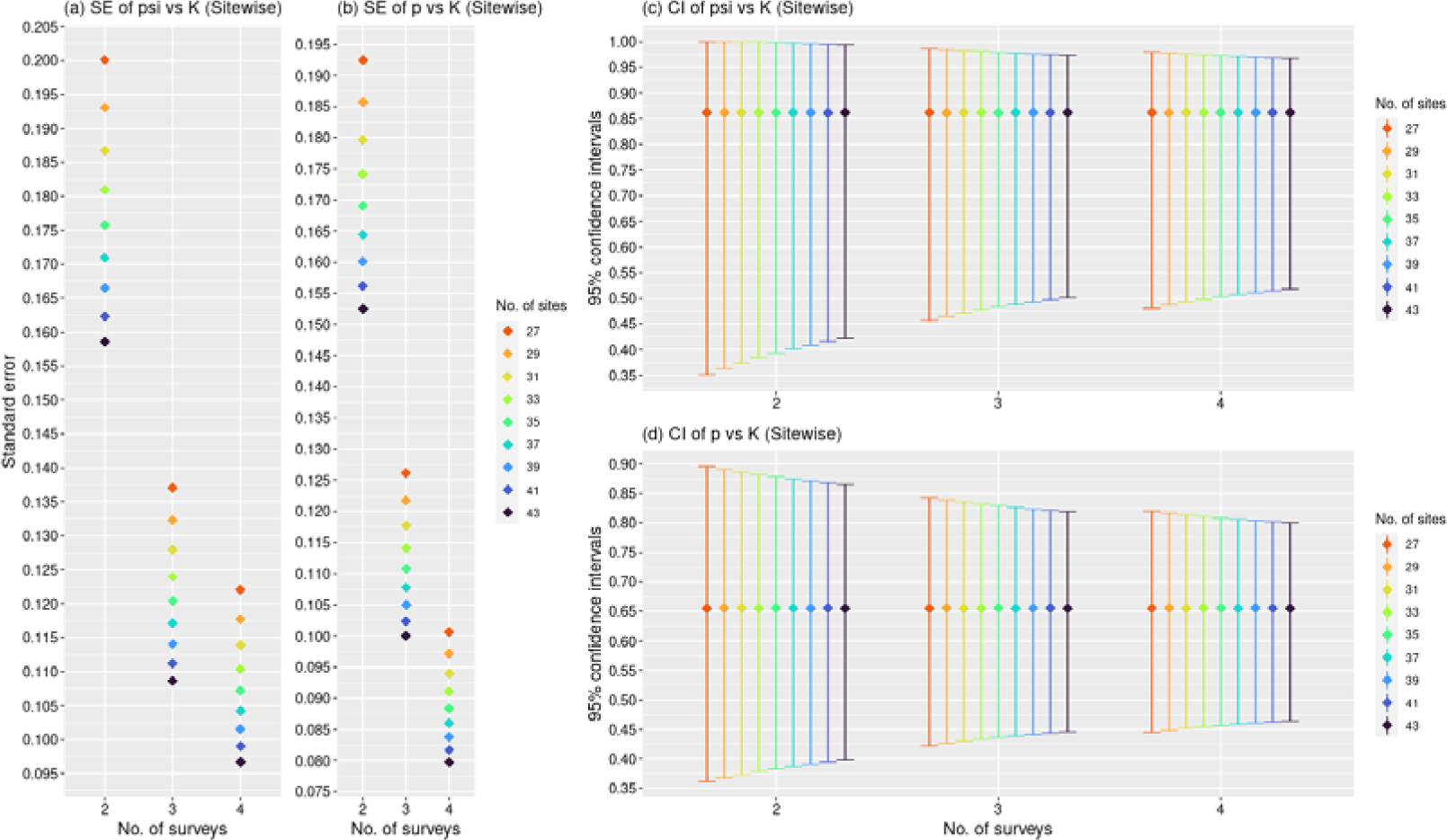
The precision parameters of estimates of ψ and p of *M. fairbanki*

**FIGURE 11.**
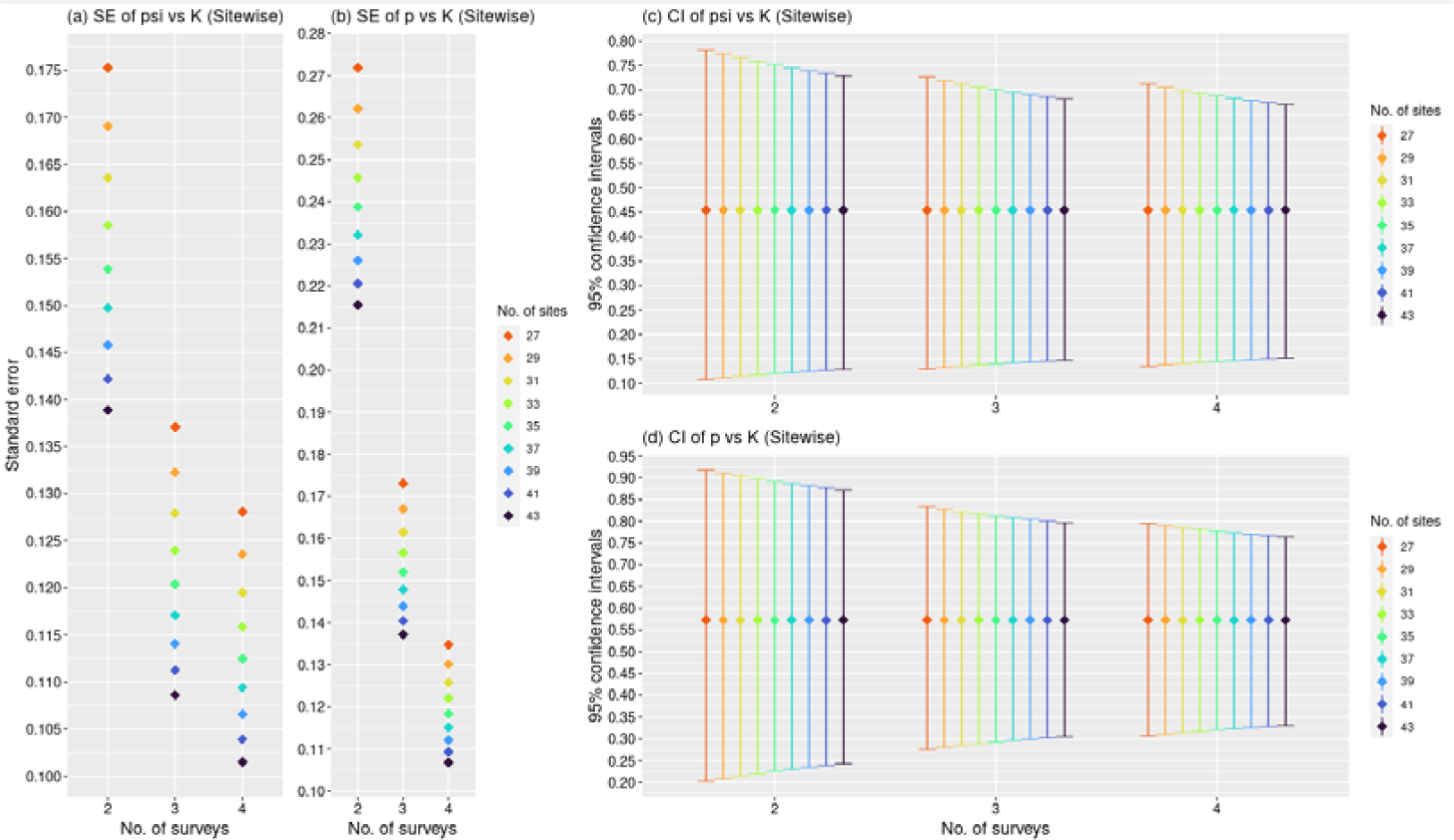
The precision parameters of estimates of ψ and p of *S. albiventris*.

## 4. DISCUSSION

*M. fairbanki* was located in all plots except one, indicating a widespread distribution in Idukki landscape, where *S. albiventris* was observed only in about half of the plots, indicating a rarer distribution status. Occupancy estimates for both the species were lower when carried out using RPresence (mean model-averaged estimates), due to the non-usage of additional data collected before or after the standardized survey times: 10.11% lesser for *M. fairbanki* and 6.42% lesser for *S. albiventris*, indicating the necessity to incorporate informal data for better understanding of distribution. Both ψ and p are generally higher for *M. fairbanki* compared to *S. albiventris*, thus proving the initial hypothesis true.

The larger number of significant models for *M. fairbanki* is due to the higher number of detections of this species across the landscape, making it difficult to pinpoint whether some of the covariates have a significant effect on their occupancy and detection. Conversely, since *S. albiventris* is rarer, the number of significant models is low and it’s easier to pinpoint which covariates are affecting their occupancy and detection in what way.

The estimates in the occupancy models (both MS and SM format) are not very reliable due to the absence of sites where *S. albiventris* is found alone and very few sites where both species are absent. But, it gives us information about what results may be expected with a larger sample size capable of detecting effects. One major result is that while there is no effect on occupancy, presence or detection of one species may result in higher detection rates for the other species. This effect is more pronounced for *S. albiventris*, whose detection is significantly higher when *M. fairbanki* is also present/detected. But this might be simply due to the presence of *M. fairbanki* through much of the landscape. More research is needed on this aspect.

Elevation is a covariate showing significant positive effect on ψ for both species, and while the pattern of its effect on the ψ of *S. albiventris* is clear, it is unclear in the case of *M. fairbanki*. In the former, there is an near exponential curve till about 1750 m a.s.l., after which ψ plateaus, indicating a preference for elevation above 1750 m a.s.l.. This finding is consistent with the knowledge that these species are generally found in high altitudes. *Sholicola* spp. are in greater abundance in the plantations above 1,500 m a.s.l. compared to plantations at the range of 1,000 to 1,500 m a.s.l (Robin & Sukumar, 2002). Same goes for *Montecincla* spp. (Praveen & Nameer, 2012). Thus, the initial hypothesis regarding the effect of elevation is confirmed. Slope is not significantly inversely correlated to elevation (see Supporting information), yet it shows significant negative effects on occupancy. So, *M. fairbanki* likely prefer less uneven terrain.

The absence of foot/dirt road decreased the occupancy of *M. fairbanki*, indicating an adaptation to some level of disturbance, and possibility of them using such pathways for some aspect of their behaviour, such as group cohesion or mating. This species may be positively influenced by visibility, as one would expect in a gregarious species who would need to keep track of other members. While the current study doesn’t find any evidence of the effect of human accessibility on *S. albiventris* occupancy, in an earlier study, encounter rate for *Sholicola* spp. were found to be four times higher in anthropogenically fragmented *shola* forests compared to unfragmented *shola* forests (Robin & Sukumar, 2002).

Presence of the invasive species *L. camara* is found to have a highly significant negative effect on ψ of *S. albiventris*, but doesn’t seem to affect *M. fairbanki*, thus only partially lending support to the initial hypothesis regarding the invasive species. This is concordant with the fact that *M. fairbanki* was observed using *L. camara* as pathways to move around. Findings of a study in Karnataka, at the intersection of Western and Eastern Ghats, indicate that the presence of *L. camara* decrease avian species diversity at community level (Aravind et al., 2010). The current study provide evidence that *L. camara* is doing the same in the *shola* landscapes, at least for rarer species.

*Sholicola* spp. has been found to be associated with streams (Robin & Sukumar, 2002), but the current study doesn’t find any association of the occupancy of *S. albiventris* with such wet lotic systems. Flowing waters have a very weak influence on the occupancy of *M. fairbanki*. So, this study finds very little evidence for the hypothesis that presence of water increases their occupancy. *S. albiventris* has a preference for understorey height of 1 m, and shows low ψ at values higher than 1 m. While understorey height has a significant negative correlation to elevation (Figure S8), the unique patterns indicates that the effects of both are independent.

The detection of *M. fairbanki* is lesser in windy conditions, likely because they would stick to the undergrowth during such times, but they seem to be more active during overcast conditions compared to clear sky conditions. And they are best detected before noon, when compared to the afternoon. *S. albiventris* shows higher detectability when the felt temperature is neither cool nor warm, but is neutral.

Therefore, the hypothesis that cool weather increases the detection probability is rejected. The survey number is a particularly interesting covariate significantly improving the detection of both species. Since the surveys were not conducted in a sequential manner for every sites (i.e. completing survey 1 for all sites before moving on to survey 2), it cannot be the effect of improving detection proficiency of the observer. There is only a slight shift in the means of survey numbers as expected, since surveys are chronological for individual sites (see Supporting information). This means that there is a ‘familiarisation effect’ of the observer to the site, wherein the observer develops a subconscious or conscious bias towards orienting to the direction where they last detected the species, so as to potentially maximise the detection probability.

Statistical significance could not be established for some covariates which were also found in significant models — a) For *M. fairbanki*: Area, Understorey height, Burn, Canopy height, Aspect, Lentic and Wet lotic waterbodies, Dry lotic system, and *R. niveus*, Visibility, and Neutral b) for both species: Warm and Cool. *R. niveus* is recorded to be a food source (fruits) and shelter plant for *M. fairbanki*. So, the non-significance is likely due to the non-dominance of *R. niveus* in their varied diet. More number of surveys can be useful to understand their actual effect.

There are limitations to the current study design. For example, none of the sites studied were fragmented in a way to match the plot size delineated on the map as most of them have connectivity to more plantations. Hence, effect of habitat fragmentation can’t be inferred from the data collected. There is also scope for improvement of data collection and analysis methods. In this present study, covariates that were noted down as categorical could also be collected as continuous variables such as wind in the Beaufort scale (Dawson & Bull, 1975). Other covariates can also be collected such as intensity of noise (Dawson & Bull, 1975) or distance to *shola* forest patches. The confidence in conservative assertions made with the results of the study is based on the fundamental assumption that there are interdepencies between each plots and each recurrent detection is independent of each other, but this may not be true as there maybe spatial autocorrelation and ‘trap-happy’ responses. In analysis, adjustments for spatial autocorrelation and trap response can be incorporated.

In the present study, there have been some observations of convergence failure in logistic regression used in RPresence, probably as a result of the low sample size and/or offending variables/covariates. But, resolving this may not be necessary as the Maximum Likelihood Estimates tend to remain valid (Allison, 2008). In addition, occupancy study of another endemic bird species in the *sholas* has also recorded standard errors larger than estimate for covariates, even with a much larger sample size, revealing the complexity of occupancy modelling (Lele et al., 2020). Another fundamental assumption is that accounting for non-detection will be better than not accounting for it. But a research article posits that the contrary is true (Welsh et al., 2013).

Both the endemic species in this study are facing some level of threat as per the IUCN classification. *M. fairbanki* is classified as Near Threatened whereas *S. albiventris* is classified as Vulnerable (BirdLife International 2016, 2018). Hence, this study is valuable in that it provides baseline information on their occupancy status, and also information on the factors affecting their occupancy. Once a large scale study is carried out over the full range in which these species occur, with a higher number of site replicates enough to be representative of this range, management policies can be devised to ensure their survival.

Under such policies, removal of the highly invasive *L*. *camara* can be prioritized as it affects *S. albiventris* significantly. A study carried out in south-eastern Brazil in *Eucalyptus* plantations have found out that native trees within the plantations, and the native understorey which is in the early successional stage found within mature plantations, had higher correlation to the presence of a subset of bird species found in the larger landscape (Millan et al., 2015). This is similar to the observations made in Idukki landscape (increased presence of *shola* vegetation under plantations showing increased diversity and abundance). Therefore, forestry managers can aid conservation of birds in *Eucalyptus* stands in tropical habitats via maintenance of native trees within the stands and through adoption of techniques triggering regeneration of the understorey during preparation of site, as well as rotation of stand (Millan et al., 2015). This phenomenon is corroborated for the mature plantations in Western Ghats (Arasumani, 2018).

There are other not-so-apparent factors that can affect the species, which needs to be monitored systematically. Climate change, which has a profound effect of the distribution of avian species in mountainous regions throughout the world, is one of them (Freeman et al., 2018). An international assessment indicates that between 184 to 327 avian species (among 1009 species) in the montane regions are expected to lose greater than half of their range, as a result of range contractions which are warming-induced (Pimm et al., 2014). Rampant deforestation is another factor which needs monitoring. A loss summing upto about 50% of the *shola* montane forests since the 1850s till the mid 1990s have been observed (Sukumar et al., 1995). In Idukki, there has been a forest loss of about 2062.3 Km^2^ from 1925 to 2012, constituting about 41.1% of loss. Hydro-electric projects were partly to blame for this, along with agriculture (shows decreasing trend now) and plantations (increasing trend ∼ + 81.1 Km^2^ from 1975 to 2012) (Ramachandran & Reddy, 2017).

The rise in number of plantations (mostly *Eucalyptus*), and the fact that *L. camara* is strongly associated with *Eucalyptus* (Jobin et al., 2023), means that another management effort that needs to be undertaken is to limit the spread/encroachment of *Eucalyptus* plantations into *shola* habitats. Alongside the monitoring in the *shola* forests, these two species would also require monitoring at plantations to understand if they can adapt to the ongoing conversion of habitats. Given the Vulnerable status of *Sholicola albiventris* (compared to the Near Threatened status of *Montecincla fairbanki*), more immediate focus on research may be given to this species. But a landscape level approach is necessary to help safeguard all such range-restricted species in peril, from known and potential threats, in the ancient evolutionary laboratories we know as the *shola* sky islands.

## Supporting information

Data for analyses

Survey details

## ACKNOWLEDGMENTS

Permissions were taken when entering private areas. I am grateful towards Randeep Singh for the guidance given for the project, and Rajan Pilakandy and Moshi Kiran for help on-field. I would like to thank all the locals and workers in the Idukki landscape who have provided minor help during the study. I am grateful towards Therese Donovan, James Hines, and Darryl MacKenzie, for the occupancyTuts, which provides great documentation for the RPresence package. Additional thanks to Mr. Hines who has constantly provided help during the course of writing the code for analyses, by correcting errors, clarifying queries, and even updating the package to make it easier for me (and others).

## DATA AVAILABILITY STATEMENT

The data that support the findings of this study are openly available at https://drive.google.com/drive/folders/1n9xb_Qd_hHT4IZA83cFSalyy4GUAluB4. Supplemental Data is also directly available from the bioRxiv site. These files will be deposited in a dedicated long-term repository after peer-review. The only exception is the shapefile of the study sites, which is from a third party (Arasumani et al., 2018) and can be only be shared with their permission.

## SUPPLEMENTARY INFORMATION

The supplementary file containing all the supplementary figures, methods, and tables is available at https://drive.google.com/drive/folders/1n9xb_Qd_hHT4IZA83cFSalyy4GUAluB4. It is also directly available from the bioRxiv site. This file will be deposited in a dedicated long-term repository after peer-review. The Supplementary Code with extensive documentation for 100% reproducibility, can be found in GitHub at https://github.com/paulvpop/Montecincla-Sholicola-occupancy.git

