## Supporting information for "An investigation into the patterns of occupancy and detection of two endemic montane bird species, in Southern Western Ghats, India"

Paul Pop

**Table of Contents:**

| Title | Page no. |
| --- | --- |
| Figure S1. Landscapesin Idukki. | 3 |
| Figure S2. Covariates | 4 |
| Figure S3. Covariates | 5 |
| Figure S4. Covariates | 6 |
| Figure S5. Continuous site-specific covariates - part A. | 7 |
| Figure S6. Continuous site-specific covariates - part B. | 8 |
| Figure S7. Top: Site-specific categorical covariates;Bottom: Survey-specific categorical covariates. | 9 |
| Figure S8. Correlations between continuous covariates. | 10 |
| Table S1: QAICc table of *Montecincla fairbanki* for single-season, single-species models | 10 |
| Table S2: Model-averaged psi estimates and precision parameters for *M. fairbanki*. | 13 |
| Table S3: Model-averaged p estimates and precision parameters for *M. fairbanki*. | 14 |
| Table S4: QAICc table of *Sholicola albiventris* for single-season, single-species models*.* | 18 |
| Table S5: Model-averaged psi estimates and precision parameters for *S. albiventris*. | 19 |
| Table S6: Model-averaged p estimates and precision parameters for *S. albiventris*. | 20 |
| Table S7. QAICc table of single-season, two-species models: MS format: Species A = *M. fairbanki*, Species B = *S. albiventris*. | 23 |
| Figure S9: Model-averaged rA estimates (det. prob. of *A*, given both species are present) and 95% confidence intervals for Two-species models. | 24 |
| Figure S10: Model-averaged rBA estimates (det. prob. of *A*, given *B* was detected) and 95% confidence intervals for Two-species models. | 24 |
| Figure S11: Model-averaged rBa estimates (det. prob. of *A*, given *B* was not detected) and 95% confidence intervals for Two-species models. | 25 |
| Table S8: Model-averaged r^M^ estimates (det. prob. of *M. fairbakii*, given both species are present) and precision parameters for Two-species models (MS format). | 25 |
| Table S9: Model-averaged r^SM^ estimates (det. prob. of *S. albiventris*, given *M. fairbanki* was detected) and precision parameters for Two-species models (MS format). | 29 |
| Table S10. Model-averaged r^Sm^ estimates (det. prob. of *S. albiventris*, given *M. fairbanki* was not detected) and precision parameters for Two-species models (MS format). | 33 |
| Table S11. QAICc table of single-season, single-species models: SM format: Species A = *S. albiventris*, Species B = *M. fairbanki.* | 36 |
| Table S12: Model-averaged r^S^ estimates (det. prob. of *S. albiventris*, given both species are present) and precision parameters for Two-species models (SM format). | 37 |
| Table S13: Model-averaged r^MS^ estimates (det. prob. of *M. fairbanki*, given *S. albiventris* was detected) and precision parameters for Two-species models (SM format). | 41 |
| Table S14: Model-averaged r^Ms^ estimates (det. prob. of *M. fairbanki*, given *S. albiventris* was not detected) and precision parameters for Two-species models (SM format). | 44 |
| Table S15. The counts and weights of covariates influencing (a) psi and (b) p in the models of the QAICc table of *M. fairbanki*. | 48 |
| Table S16. The counts and weights of covariates influencing (a) psi and (b) p in the models of the QAICc table of *S. albiventris*. | 49 |
| Table S17. The counts and weights of covariates influencing (a) psi and (b) p in the models of the QAICc table of single-season, Two-species models: MS format. | 49 |
| Table S18. The counts and weights of covariates influencing (a) psi and (b) p in the models of the QAICc table of single-season, Two-species models: SM format. | 49 |
| Table S19. Simulated psi and p estimates and associated precision parameters of *M. fairbanki* under varying conditions of no. of sites and number of surveys. | 50 |
| Table S20. Simulated psi and p estimates and associated precision parameters of *S. albiventris* under varying conditions of no. of sites and number of surveys. | 51 |
| Figure S12: Correlation between survey numbers and date numbers. | 52 |

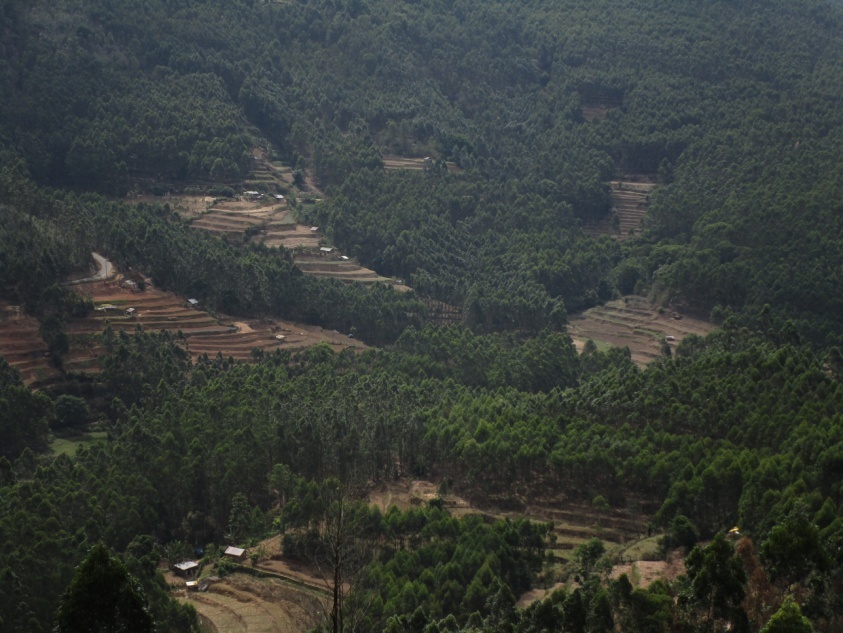

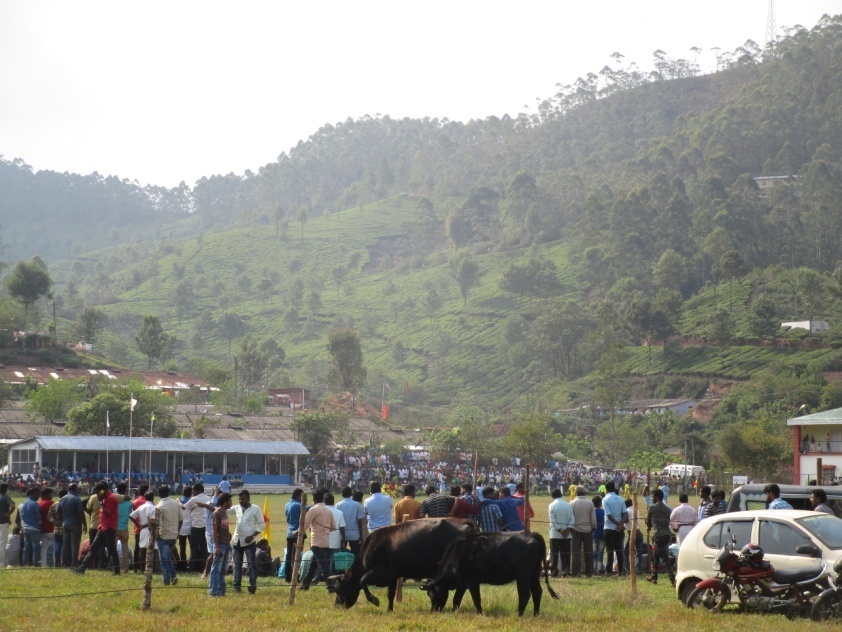

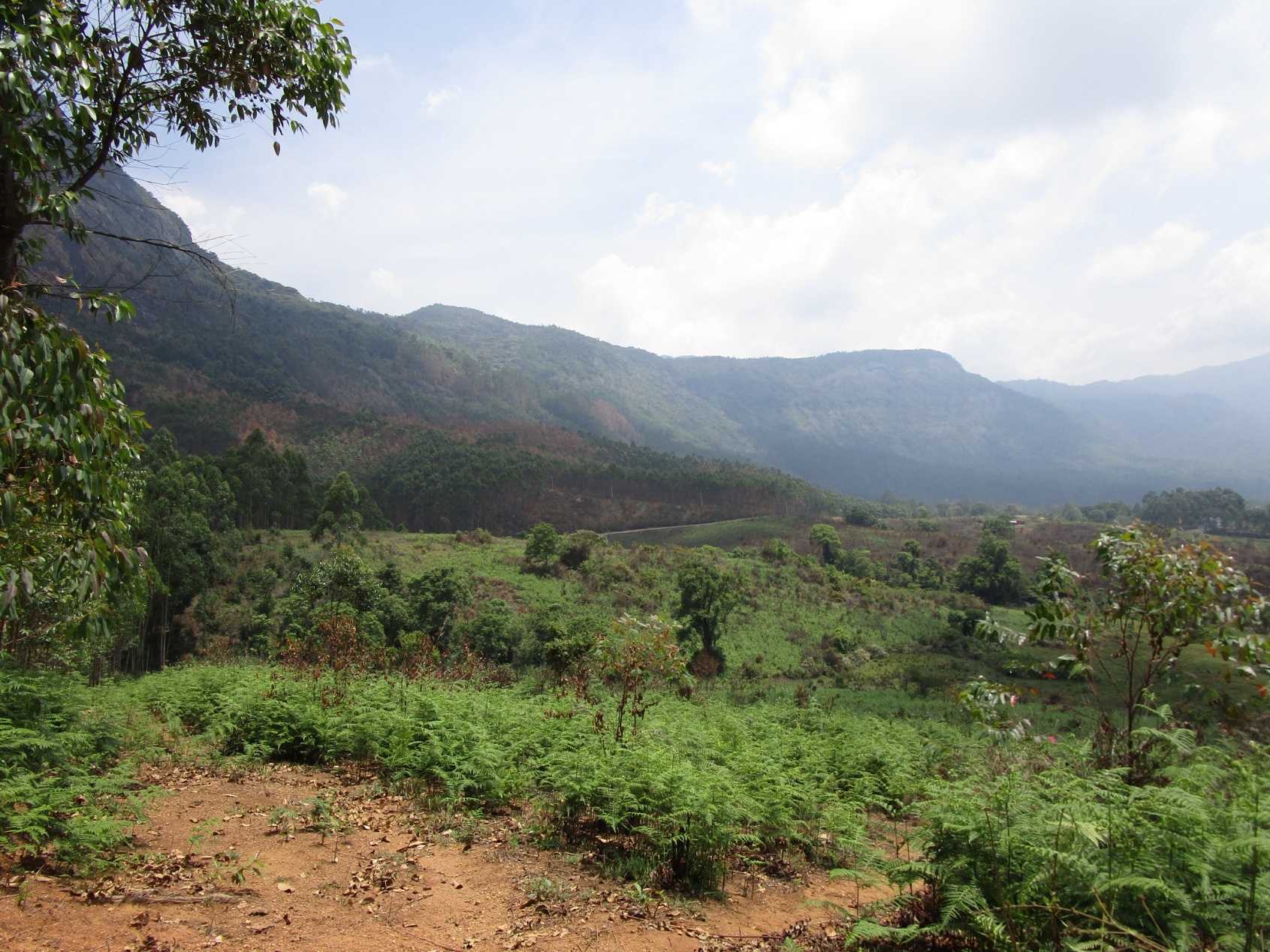
Figure S1. Landscapesin Idukki: Clockwise from bottom left - Munnar; Kanthaloor and Koviloor. Image taken using Canon IXUS 190.

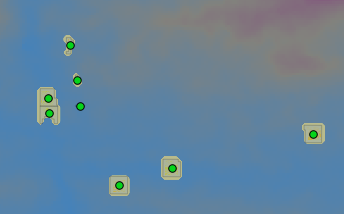

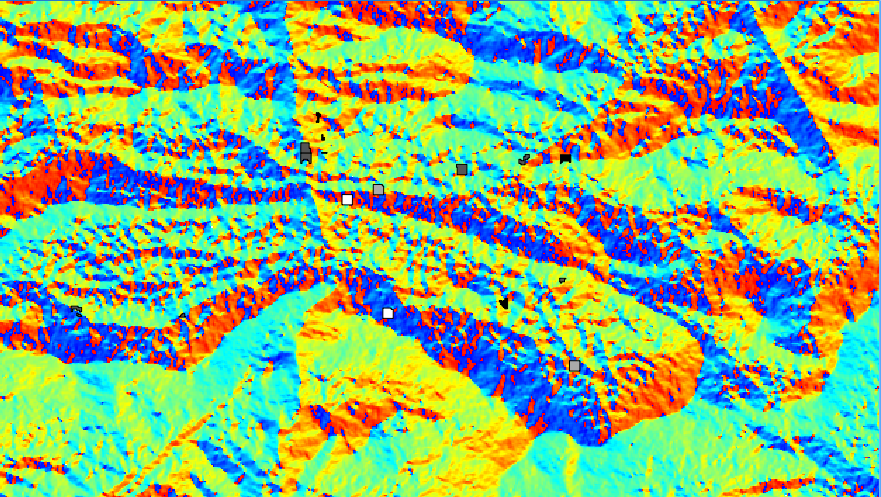

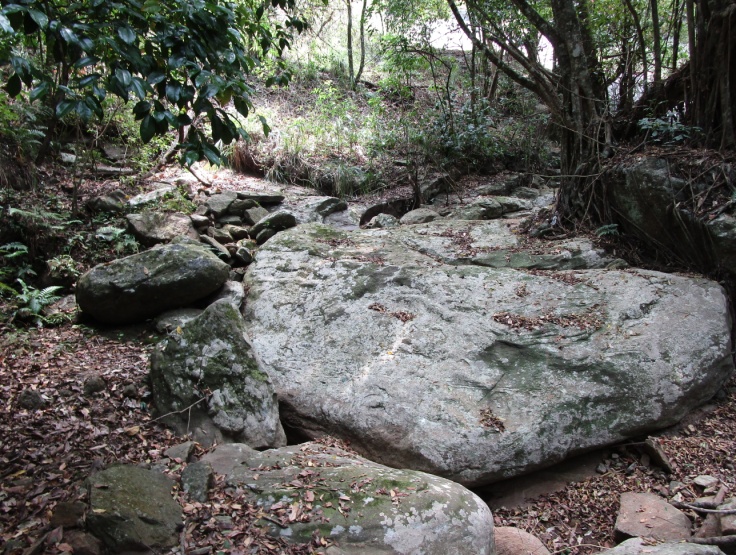

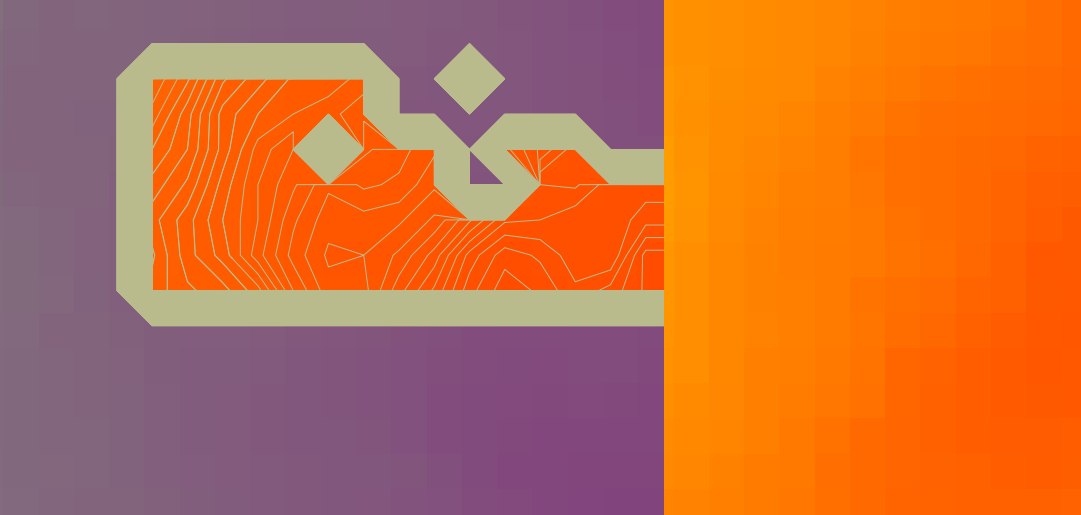

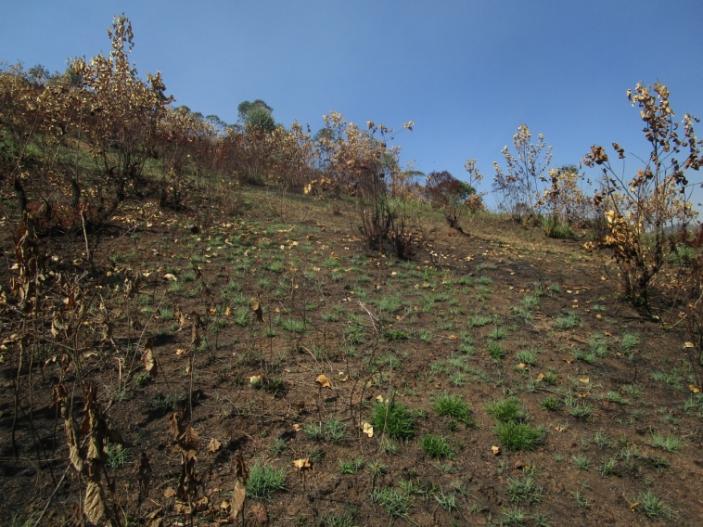

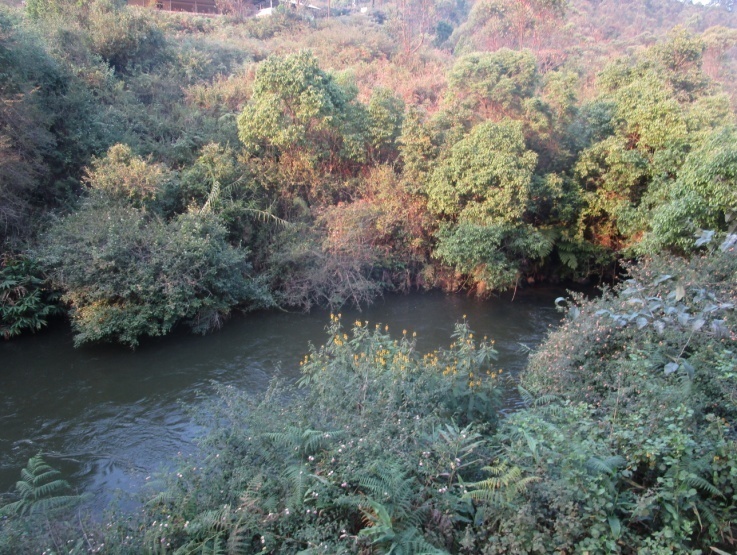

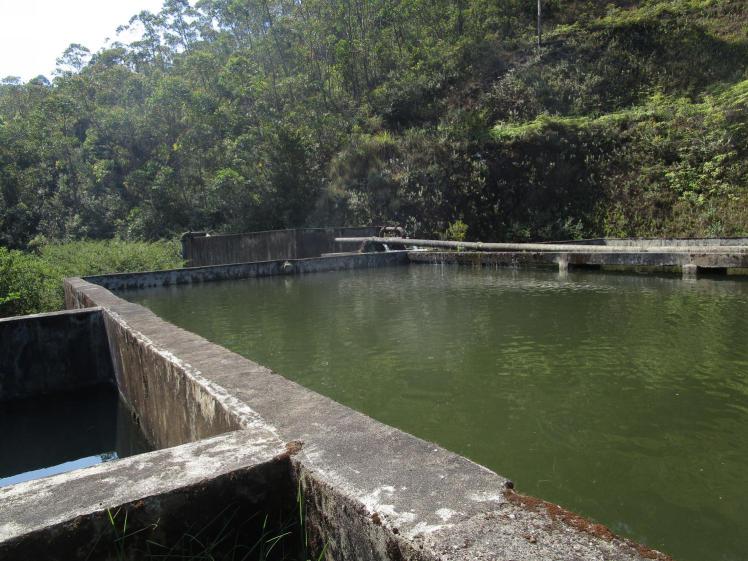

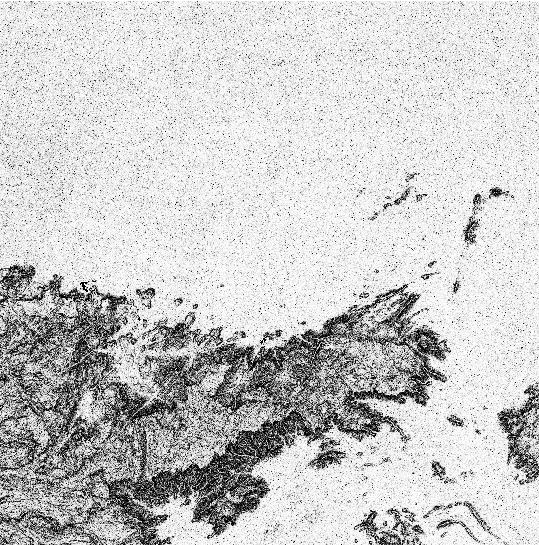
Figure S2. Covariates: Clockwise from top left: Burn; Plot Area; Elevation (derived from contour lines); Slope; Lentic waterbody; Wet lotic waterbody; Dry lotic system; and Aspect. Images taken using Canon IXUS 190 or derived from the USGS ASTER GDEM imagery using Quantum GIS version 1.8.0.

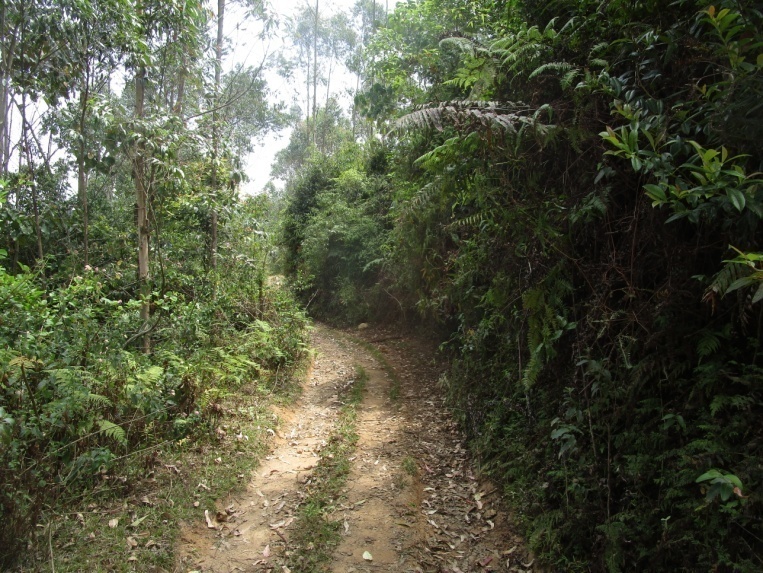

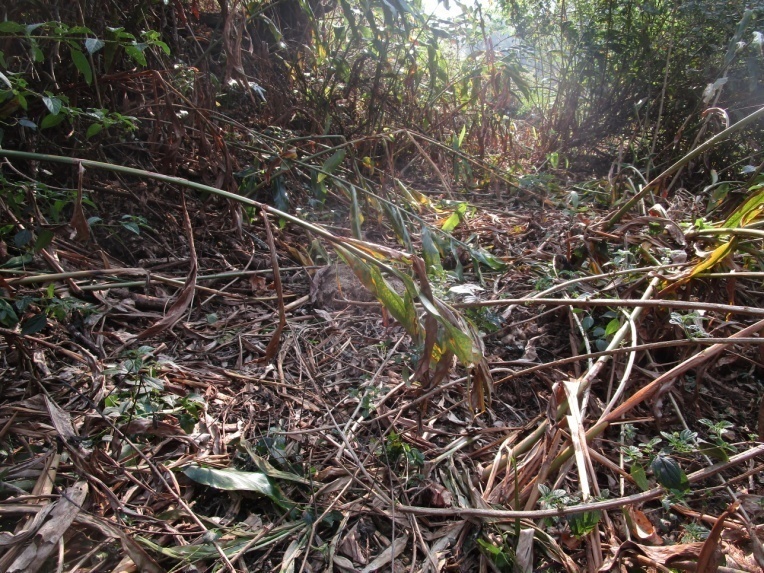

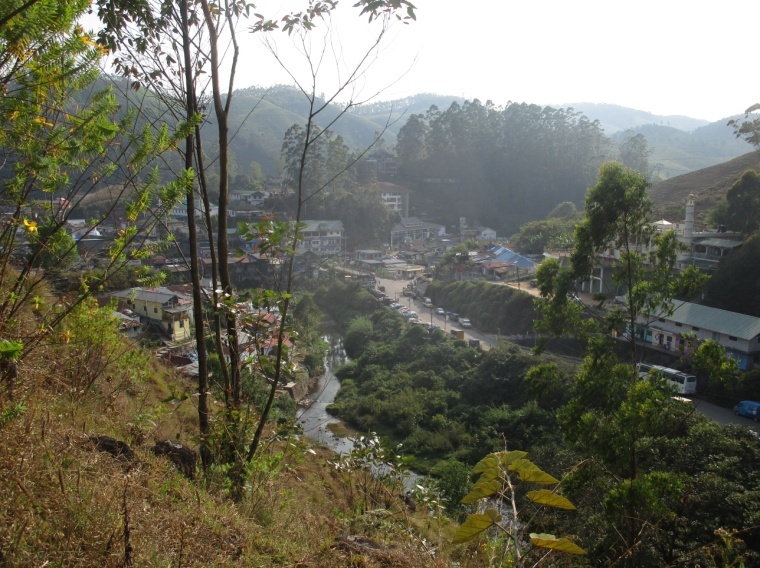

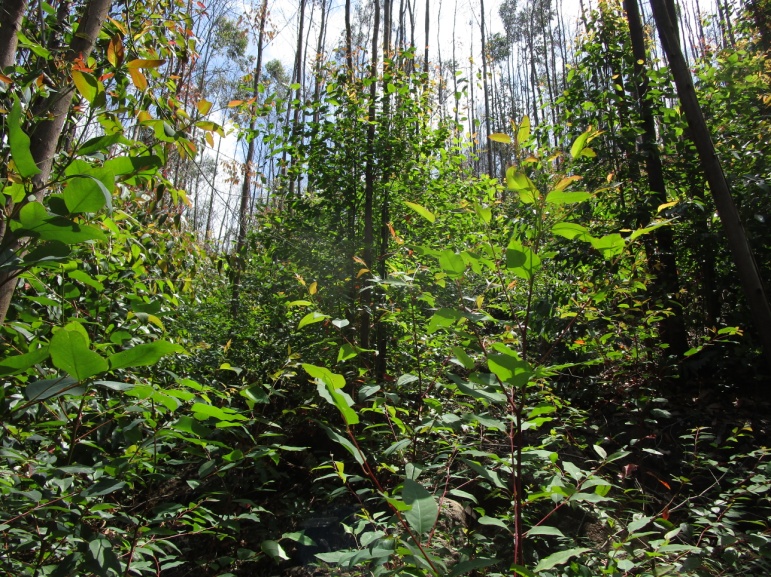

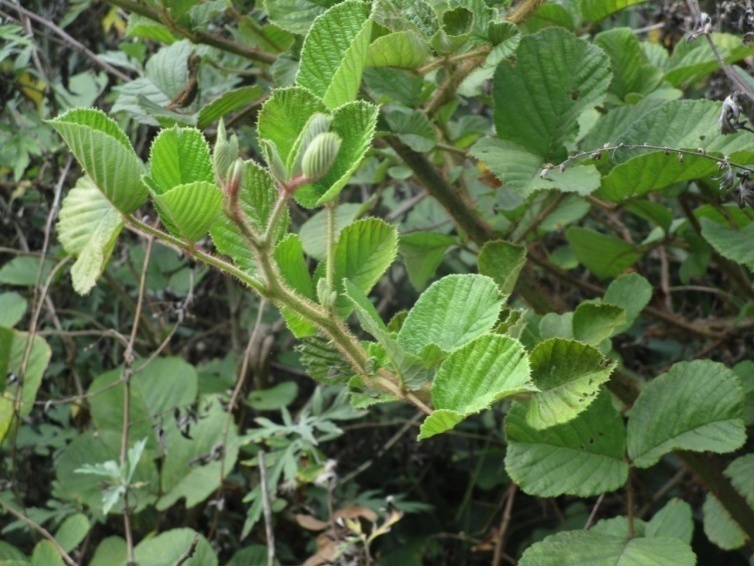

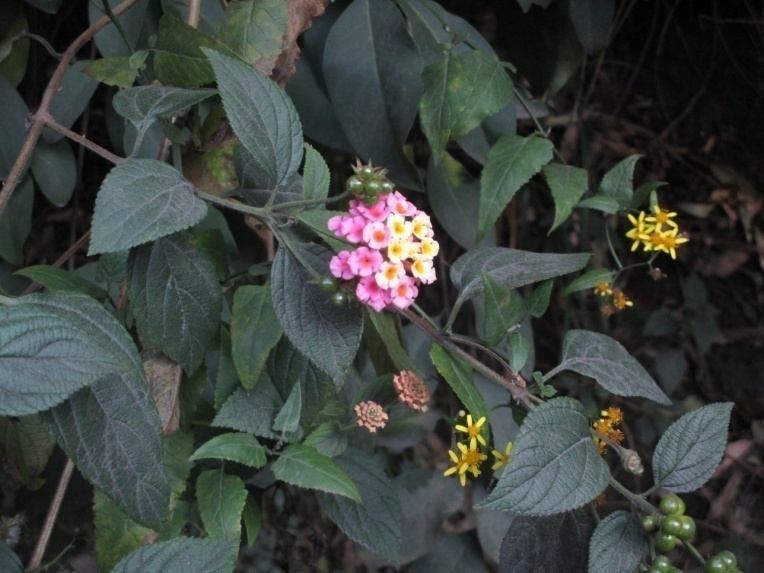
Figure S3. Covariates: Clockwise from top left - No road (Animal (elephant) trail); Dirt/foot road; Metalled road; *Rubus niveus*; *Lantana camara;* and Visibility. Images taken using Canon IXUS 190.

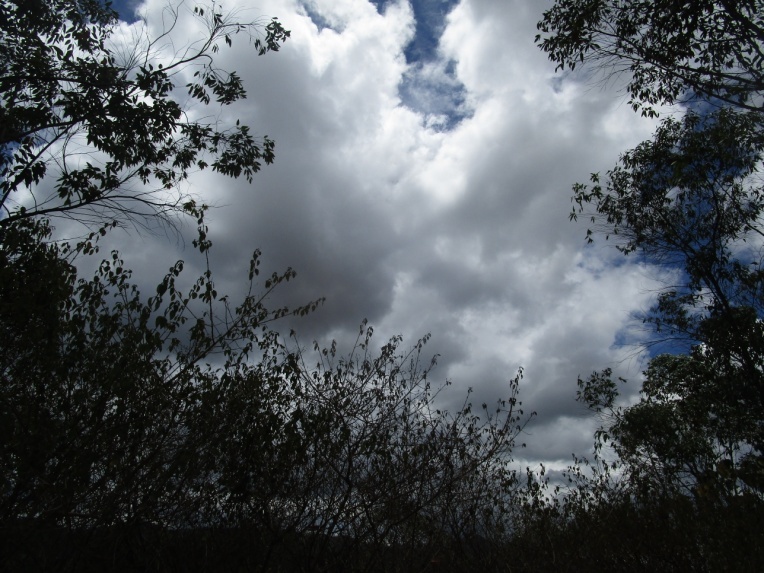

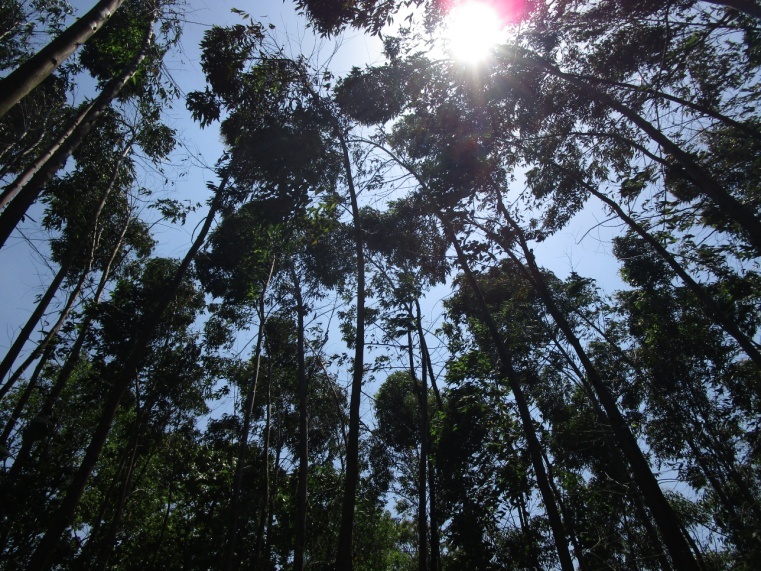

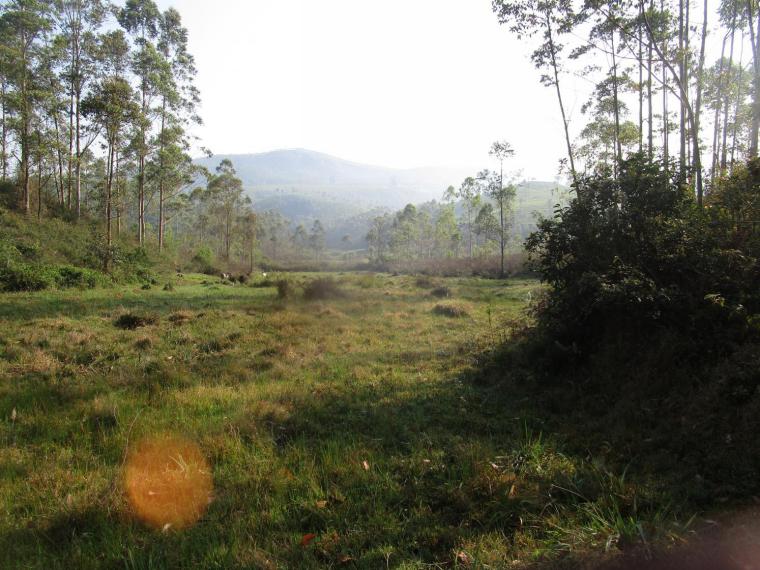

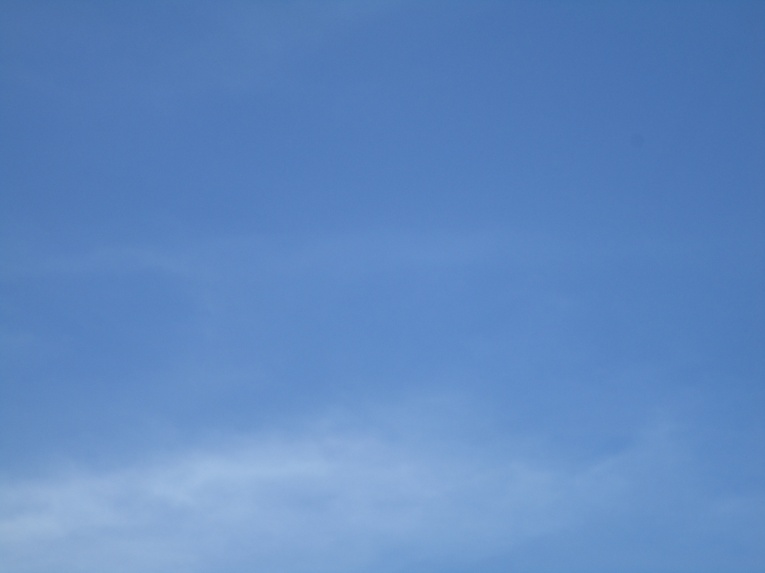

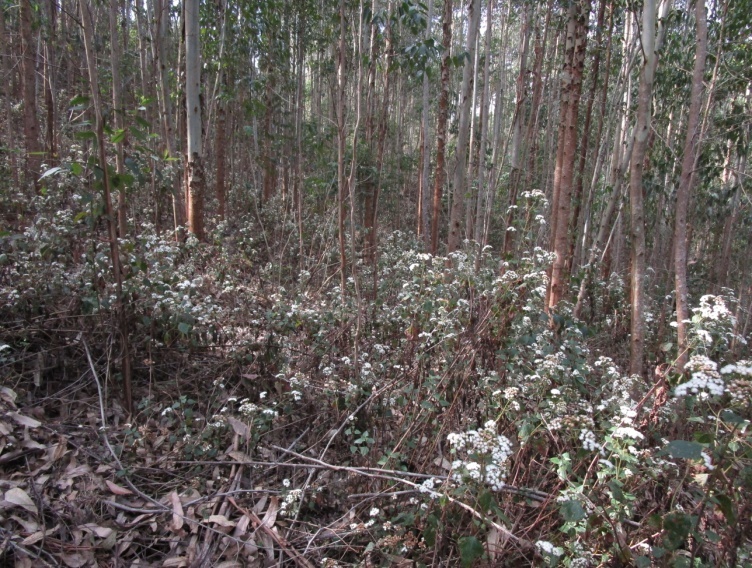

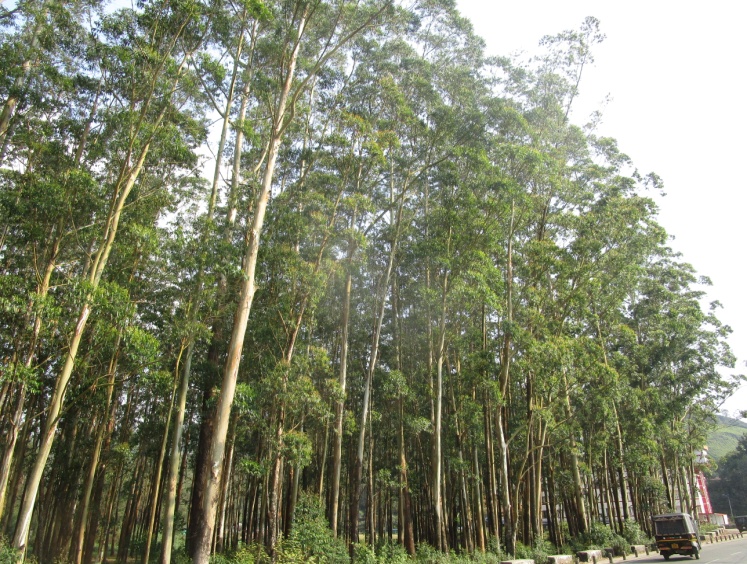

Figure S4. Covariates: Clockwise from top left: Canopy height; Understorey height; Clear skies; Overcast skies; Windy weather; and Sunny weather. Images taken using Canon IXUS 190.

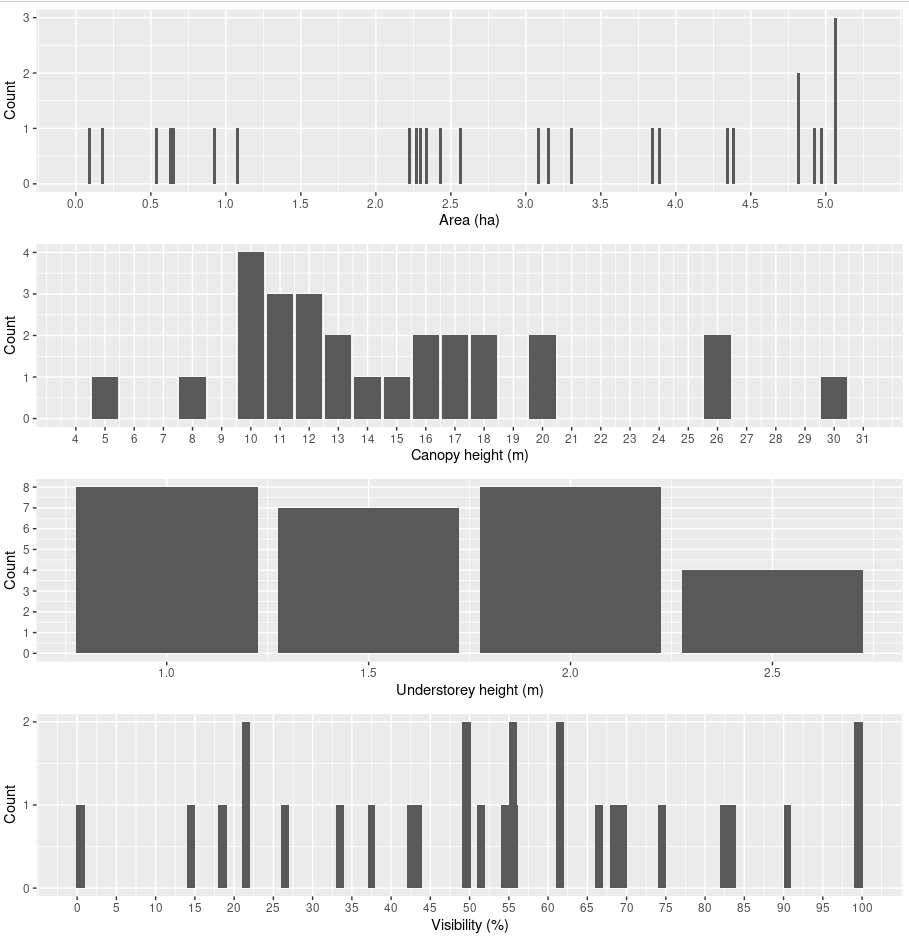

Figure S5. Continuous site-specific covariates - part A.

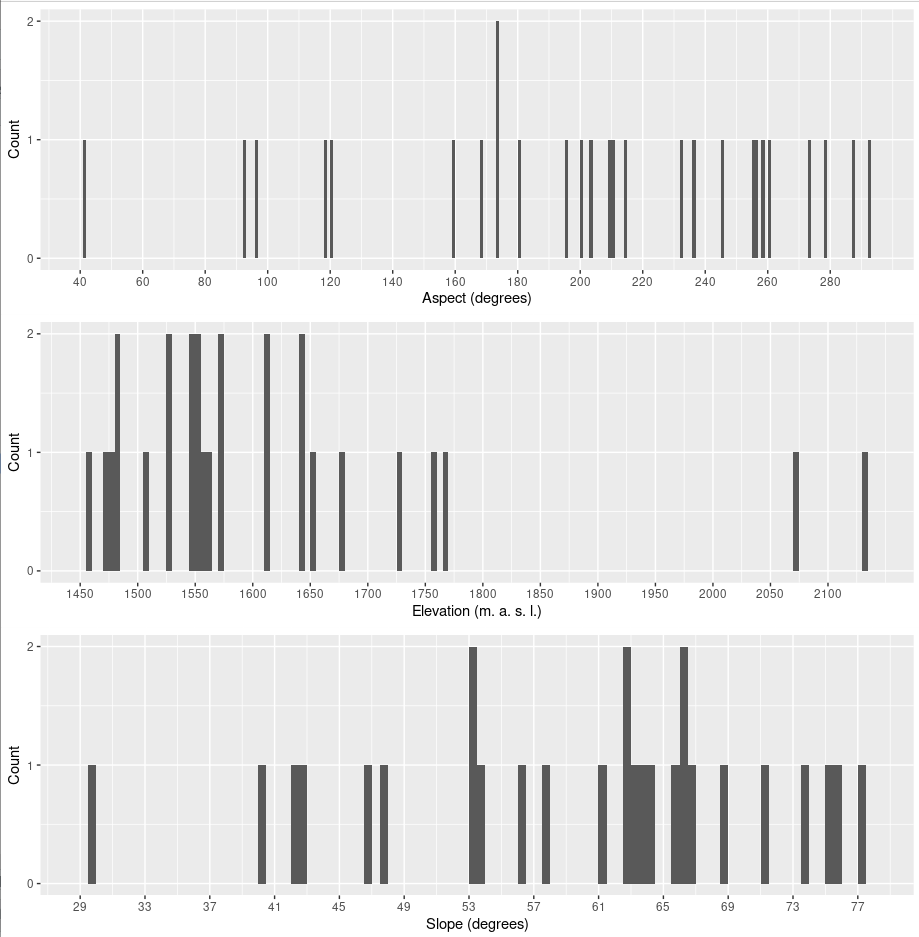

Figure S6. Continuous site-specific covariates - part B.

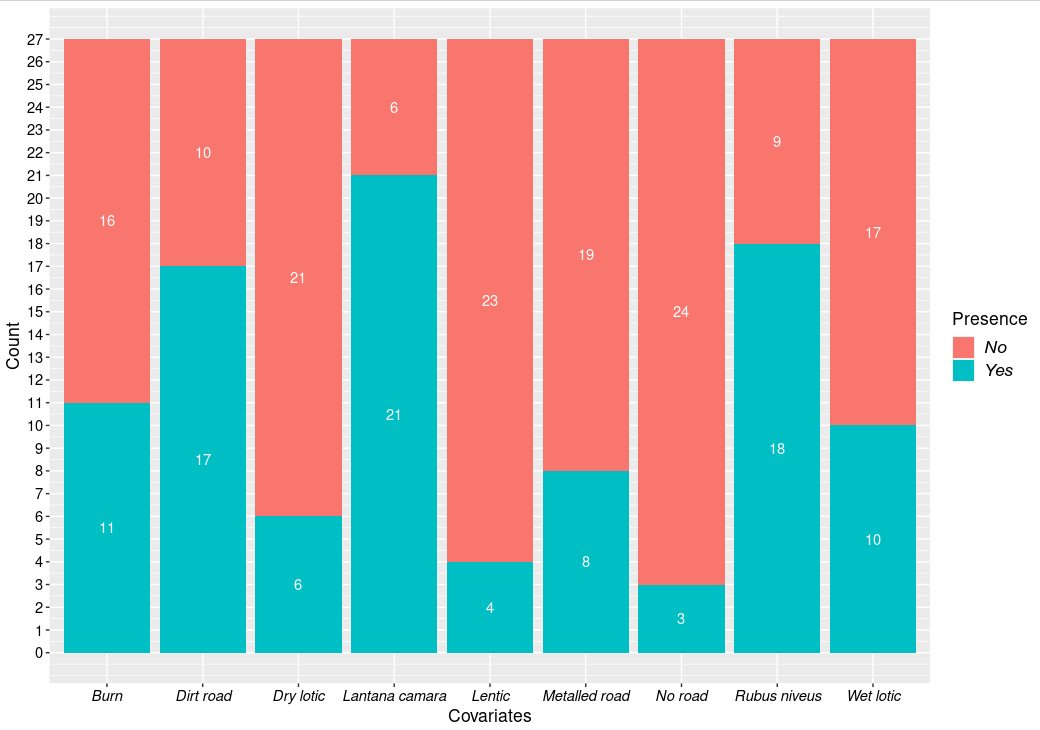

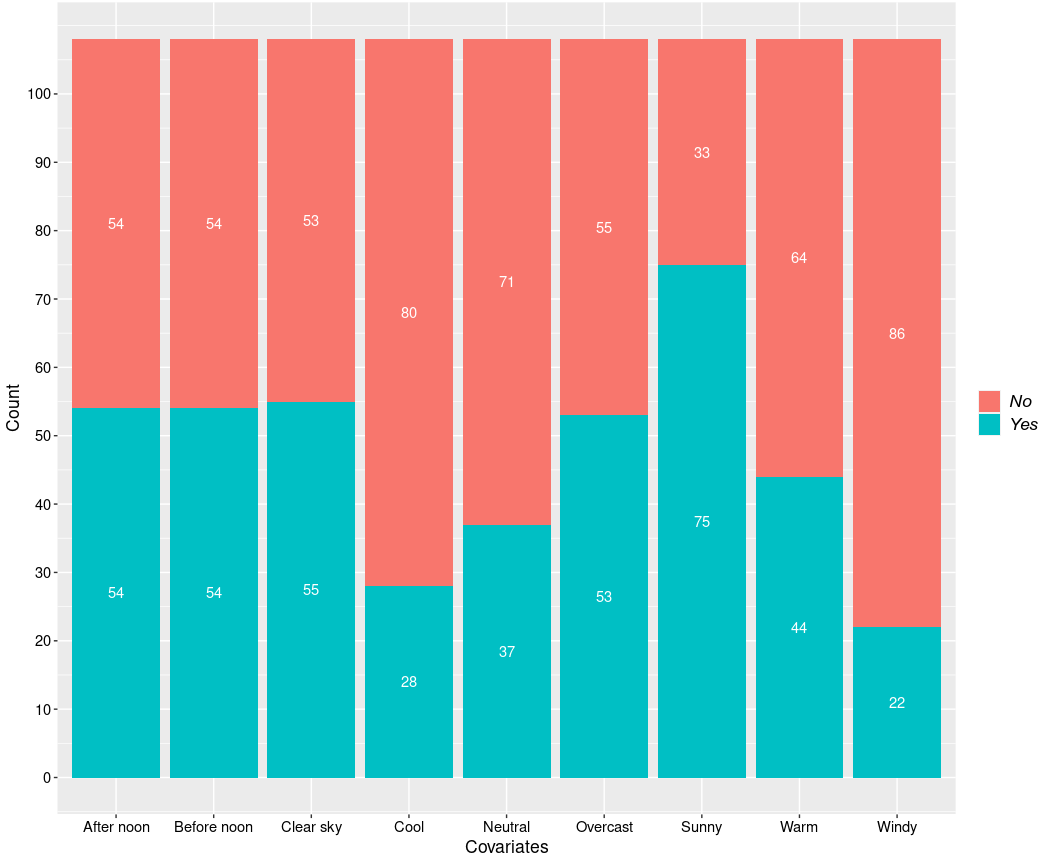

Figure S7. Top: Site-specific categorical covariates; Bottom: Survey-specific categorical covariates.

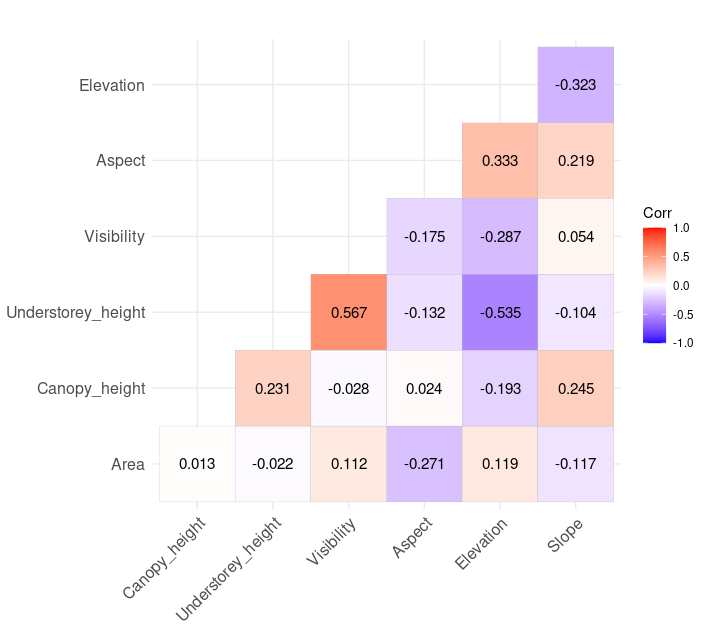

Figure S8. Correlations between continuous covariates.

Table S1: QAICc table of *Montecincla fairbanki* for single-season, single-species models.

| **Model** | **QAICc** | **neg2ll** | **n.**  **par** | **warn.conv** | **DQAICc** | **modlike** | **wgt** |
| --- | --- | --- | --- | --- | --- | --- | --- |
| psi(No road)p(1) | 102.2332 | 134.5317 | 3 | 7.00 | 0.0000 | 1.0000 | 0.0579 |
| psi(No road)p(Windy) | 102.5148 | 131.0082 | 4 | 7.00 | 0.2816 | 0.8687 | 0.0503 |
| psi(No road)p(Windy+survey) | 103.1693 | 127.6382 | 5 | 7.00 | 0.9361 | 0.6262 | 0.0362 |
| psi(1)p(1) | 103.2735 | 139.5966 | 2 | 7.00 | 1.0403 | 0.5944 | 0.0344 |
| psi(1)p(Windy) | 103.2770 | 136.0069 | 3 | 7.00 | 1.0438 | 0.5934 | 0.0343 |
| psi(No road)p(BefNoon) | 103.3555 | 132.1963 | 4 | 7.00 | 1.1223 | 0.5706 | 0.0330 |
| psi(Dirt road)p(1) | 103.8157 | 136.7682 | 3 | 7.00 | 1.5825 | 0.4533 | 0.0262 |
| psi(1)p(survey) | 103.8602 | 136.8311 | 3 | 7.00 | 1.6270 | 0.4433 | 0.0256 |
| psi(Slope)p(1) | 103.8980 | 136.8845 | 3 | 7.00 | 1.6648 | 0.4350 | 0.0252 |
| psi(Elevation)p(1) | 103.9938 | 137.0199 | 3 | 7.00 | 1.7606 | 0.4147 | 0.0240 |
| psi(Dirt road)p(Windy) | 104.1387 | 133.3032 | 4 | 7.00 | 1.9055 | 0.3857 | 0.0223 |
| psi(No road)p(Windy+BefNoon) | 104.1600 | 129.0383 | 5 | 7.00 | 1.9267 | 0.3816 | 0.0221 |
| psi(1)p(BefNoon) | 104.1645 | 137.2612 | 3 | 7.00 | 1.9313 | 0.3807 | 0.0220 |
| psi(1)p(AftNoon) | 104.1645 | 137.2612 | 3 | 7.00 | 1.9313 | 0.3807 | 0.0220 |
| psi(Visibility)p(1) | 104.3185 | 137.4788 | 3 | 7.00 | 2.0853 | 0.3525 | 0.0204 |
| psi(Dry lotic)p(1) | 104.3805 | 137.5664 | 3 | 3.59 | 2.1472 | 0.3418 | 0.0198 |
| psi(Dirt road)p(survey) | 104.6336 | 134.0027 | 4 | 7.00 | 2.4004 | 0.3011 | 0.0174 |
| psi(No road)p(Windy+survey+BefNoon) | 104.7020 | 125.0799 | 6 | 7.00 | 2.4688 | 0.2910 | 0.0168 |
| psi(No road)p(Windy+survey+AftNoon) | 104.7020 | 125.0799 | 6 | 7.00 | 2.4688 | 0.2910 | 0.0168 |
| psi(Slope)p(Windy+survey) | 104.7756 | 129.9083 | 5 | 7.00 | 2.5423 | 0.2805 | 0.0162 |
| psi(No road)p(Visibility) | 104.7989 | 134.2363 | 4 | 7.00 | 2.5657 | 0.2772 | 0.0160 |
| psi(Area)p(1) | 104.8209 | 138.1889 | 3 | 7.00 | 2.5877 | 0.2742 | 0.0159 |
| psi(Elevation)p(Windy+survey) | 104.8555 | 130.0213 | 5 | 7.00 | 2.6223 | 0.2695 | 0.0156 |
| psi(Understorey height)p(1) | 104.9282 | 138.3405 | 3 | 7.00 | 2.6950 | 0.2599 | 0.0150 |
| psi(No road)p(ClrSky+BefNoon) | 105.0035 | 130.2304 | 5 | 7.00 | 2.7702 | 0.2503 | 0.0145 |
| psi(Dry lotic)p(Windy+survey) | 105.2385 | 130.5626 | 5 | 7.00 | 3.0053 | 0.2225 | 0.0129 |
| psi(Visibility)p(Windy+survey) | 105.2627 | 130.5968 | 5 | 7.00 | 3.0295 | 0.2199 | 0.0127 |
| psi(Rubus niveus)p(1) | 105.4212 | 139.0372 | 3 | 7.00 | 3.1879 | 0.2031 | 0.0118 |
| psi(Aspect)p(1) | 105.4671 | 139.1021 | 3 | 7.00 | 3.2338 | 0.1985 | 0.0115 |
| psi(1)p(ClrSky) | 105.4869 | 139.1301 | 3 | 7.00 | 3.2537 | 0.1966 | 0.0114 |
| psi(1)p(Ovrcst) | 105.4869 | 139.1301 | 3 | 7.00 | 3.2537 | 0.1966 | 0.0114 |
| psi(Canopy height)p(1) | 105.4981 | 139.1459 | 3 | 7.00 | 3.2648 | 0.1955 | 0.0113 |
| psi(No road)p(Visibility+Windy) | 105.5532 | 131.0073 | 5 | 7.00 | 3.3199 | 0.1901 | 0.0110 |
| psi(Lentic)p(1) | 105.5753 | 139.2550 | 3 | 7.00 | 3.3420 | 0.1881 | 0.0109 |
| psi(Wet lotic)p(1) | 105.5843 | 139.2677 | 3 | 7.00 | 3.3510 | 0.1872 | 0.0108 |
| psi(1)p(Cool) | 105.6744 | 139.3951 | 3 | 7.00 | 3.4412 | 0.1790 | 0.0104 |
| psi(Burn)p(1) | 105.7007 | 139.4322 | 3 | 7.00 | 3.4674 | 0.1766 | 0.0102 |
| psi(Area)p(Windy+survey) | 105.7167 | 131.2384 | 5 | 7.00 | 3.4834 | 0.1752 | 0.0101 |
| psi(1)p(Sunny) | 105.7510 | 139.5034 | 3 | 7.00 | 3.5178 | 0.1722 | 0.0100 |
| psi(1)p(Warm) | 105.7571 | 139.5120 | 3 | 7.00 | 3.5239 | 0.1717 | 0.0099 |
| psi(Metalled road)p(1) | 105.7819 | 139.5470 | 3 | 7.00 | 3.5486 | 0.1696 | 0.0098 |
| psi(Understorey height)p(Windy+survey) | 105.8046 | 131.3627 | 5 | 7.00 | 3.5714 | 0.1677 | 0.0097 |
| psi(1)p(Neutral) | 105.8101 | 139.5869 | 3 | 7.00 | 3.5769 | 0.1672 | 0.0097 |
| psi(Rubus niveus)p(Windy+survey) | 106.2894 | 132.0478 | 5 | 7.00 | 4.0561 | 0.1316 | 0.0076 |
| psi(Slope)p(Windy+survey+BefNoon) | 106.2957 | 127.3323 | 6 | 7.00 | 4.0625 | 0.1312 | 0.0076 |
| psi(Slope)p(Windy+survey+AftNoon) | 106.2957 | 127.3323 | 6 | 7.00 | 4.0625 | 0.1312 | 0.0076 |
| psi(Area)p(I(2^survey)) | 106.3359 | 136.4085 | 4 | 7.00 | 4.1027 | 0.1286 | 0.0074 |
| psi(Aspect)p(Windy+survey) | 106.3655 | 132.1554 | 5 | 7.00 | 4.1323 | 0.1267 | 0.0073 |
| psi(Elevation)p(Windy+survey+BefNoon) | 106.3811 | 127.4530 | 6 | 7.00 | 4.1479 | 0.1257 | 0.0073 |
| psi(Elevation)p(Windy+survey+AftNoon) | 106.3811 | 127.4530 | 6 | 7.00 | 4.1479 | 0.1257 | 0.0073 |
| psi(No road)p(Visibility+Windy+survey) | 106.5121 | 127.6381 | 6 | 7.00 | 4.2789 | 0.1177 | 0.0068 |
| psi(1)p(BefNoon+Cool) | 106.6737 | 136.8859 | 4 | 4.71 | 4.4405 | 0.1086 | 0.0063 |
| psi(Slope)p(ClrSky+BefNoon) | 106.6787 | 132.5980 | 5 | 7.00 | 4.4454 | 0.1083 | 0.0063 |
| psi(Dirt road)p(ClrSky+BefNoon) | 106.6853 | 132.6073 | 5 | 7.00 | 4.4520 | 0.1080 | 0.0062 |
| psi(1)p(Visibility+Windy+survey) | 106.6965 | 132.6232 | 5 | 7.00 | 4.4633 | 0.1074 | 0.0062 |
| psi(Elevation)p(Warm) | 106.7012 | 136.9247 | 4 | 7.00 | 4.4679 | 0.1071 | 0.0062 |
| psi(Elevation)p(ClrSky+BefNoon) | 106.7369 | 132.6803 | 5 | 7.00 | 4.5037 | 0.1052 | 0.0061 |
| psi(Dry lotic)p(Windy+survey+BefNoon) | 106.7648 | 127.9953 | 6 | 5.20 | 4.5316 | 0.1037 | 0.0060 |
| psi(Dry lotic)p(Windy+survey+AftNoon) | 106.7648 | 127.9953 | 6 | 7.00 | 4.5316 | 0.1037 | 0.0060 |
| psi(Visibility)p(Windy+survey+BefNoon) | 106.7943 | 128.0369 | 6 | 7.00 | 4.5610 | 0.1022 | 0.0059 |
| psi(Visibility)p(Windy+survey+AftNoon) | 106.7943 | 128.0369 | 6 | 7.00 | 4.5610 | 0.1022 | 0.0059 |
| psi(Dry lotic)p(ClrSky+BefNoon) | 107.1073 | 133.2038 | 5 | 3.58 | 4.8741 | 0.0874 | 0.0051 |
| psi(Visibility)p(ClrSky+BefNoon) | 107.1174 | 133.2180 | 5 | 7.00 | 4.8841 | 0.0870 | 0.0050 |
| psi(1)p(BefNoon+Cool+Windy) | 107.1942 | 133.3265 | 5 | 7.00 | 4.9609 | 0.0837 | 0.0048 |
| psi(Area)p(Windy+survey+BefNoon) | 107.2722 | 128.7123 | 6 | 7.00 | 5.0389 | 0.0805 | 0.0047 |
| psi(Area)p(Windy+survey+AftNoon) | 107.2722 | 128.7123 | 6 | 7.00 | 5.0389 | 0.0805 | 0.0047 |
| psi(Area)p(ClrSky+BefNoon) | 107.4891 | 133.7433 | 5 | 7.00 | 5.2558 | 0.0722 | 0.0042 |
| psi(No road)p(Visibility+Windy+AftNoon) | 107.4983 | 129.0319 | 6 | 7.00 | 5.2651 | 0.0719 | 0.0042 |
| psi(1)p(Cool+Windy+Ovrcst) | 107.5706 | 133.8585 | 5 | 7.00 | 5.3373 | 0.0693 | 0.0040 |
| psi(Rubus niveus)p(Windy+survey+BefNoon) | 107.8365 | 129.5099 | 6 | 7.00 | 5.6033 | 0.0607 | 0.0035 |
| psi(Rubus niveus)p(Windy+survey+AftNoon) | 107.8365 | 129.5099 | 6 | 7.00 | 5.6033 | 0.0607 | 0.0035 |
| psi(Elevation)p(BefNoon+Cool) | 107.8759 | 134.2900 | 5 | 7.00 | 5.6426 | 0.0595 | 0.0034 |
| psi(Aspect)p(Windy+survey+BefNoon) | 107.9045 | 129.6060 | 6 | 7.00 | 5.6713 | 0.0587 | 0.0034 |
| psi(Aspect)p(Windy+survey+AftNoon) | 107.9045 | 129.6060 | 6 | 7.00 | 5.6713 | 0.0587 | 0.0034 |
| psi(Area)p(survey+I(survey^2)) | 107.9314 | 134.3685 | 5 | 7.00 | 5.6982 | 0.0579 | 0.0033 |
| psi(Slope)p(Visibility+Windy+survey) | 108.1145 | 129.9027 | 6 | 7.00 | 5.8812 | 0.0528 | 0.0031 |
| psi(Elevation)p(BefNoon+Cool+Windy) | 108.6832 | 130.7065 | 6 | 7.00 | 6.4499 | 0.0398 | 0.0023 |
| psi(Elevation)p(Cool+Windy+Ovrcst) | 109.0897 | 131.2810 | 6 | 7.00 | 6.8564 | 0.0324 | 0.0019 |

Table S2: Model-averaged psi estimates and precision parameters for *M. fairbanki*.

| **Site IDs** | **Estimate** | **Standard Error** | **Lower 95% Confidence Interval** | **Upper 95% Confidence Interval** |
| --- | --- | --- | --- | --- |
| 13 | 0.8542103 | 0.13590618 | 0.4083204 | 0.9802942 |
| 77 | 0.8969234 | 0.08751371 | 0.5764510 | 0.9823424 |
| 82 | 0.8690676 | 0.11766001 | 0.4665752 | 0.9805331 |
| 34 | 0.6909203 | 0.28711186 | 0.1381553 | 0.9689179 |
| 105 | 0.8820558 | 0.10376418 | 0.5142838 | 0.9814204 |
| 108 | 0.8839535 | 0.09492694 | 0.5539564 | 0.9790441 |
| 31 | 0.8637678 | 0.12110251 | 0.4575666 | 0.9794480 |
| 110 | 0.8583428 | 0.12358665 | 0.4525066 | 0.9779842 |
| 103 | 0.8571911 | 0.11271277 | 0.4968808 | 0.9733196 |
| 4 | 0.9229368 | 0.07472200 | 0.6044004 | 0.9894606 |
| 19 | 0.8894673 | 0.08760754 | 0.5839015 | 0.9787893 |
| 96 | 0.8856502 | 0.08467157 | 0.6007049 | 0.9755344 |
| 74 | 0.9020032 | 0.07896981 | 0.6150691 | 0.9814888 |
| 102 | 0.8972254 | 0.07856748 | 0.6217113 | 0.9788910 |
| 76 | 0.8793933 | 0.11420870 | 0.4690800 | 0.9836531 |
| 21 | 0.8934443 | 0.09338733 | 0.5507752 | 0.9828596 |
| 37 | 0.8501526 | 0.13055184 | 0.4322291 | 0.9768956 |
| 72 | 0.6659567 | 0.28516879 | 0.1391319 | 0.9609255 |
| 14 | 0.9046884 | 0.08418764 | 0.5834219 | 0.9846933 |
| 104 | 0.6759544 | 0.28889938 | 0.1358938 | 0.9651188 |
| 17 | 0.8688554 | 0.12276258 | 0.4450366 | 0.9820578 |
| 60 | 0.8928459 | 0.10125310 | 0.5114600 | 0.9851449 |
| 52 | 0.8989191 | 0.09168989 | 0.5516856 | 0.9846785 |
| 91 | 0.9049155 | 0.09026449 | 0.5490818 | 0.9867339 |
| 93 | 0.8918832 | 0.09322999 | 0.5535815 | 0.9821035 |
| 84 | 0.8832803 | 0.11187470 | 0.4742765 | 0.9844912 |
| 20 | 0.9049996 | 0.09847360 | 0.5022921 | 0.9890015 |

Table S3: Model-averaged p estimates and precision parameters for *M. fairbanki*.

| **Site IDs** | **Survey no.** | **Estimate** | **Standard Error** | **Lower 95% Confidence Interval** | **Upper 95% Confidence Interval** |
| --- | --- | --- | --- | --- | --- |
| 13 | 1 | 0.6102463 | 0.10473803 | 0.3977751 | 0.7877540 |
| 77 | 1 | 0.6557339 | 0.08873230 | 0.4685304 | 0.8045102 |
| 82 | 1 | 0.6105095 | 0.10477248 | 0.3979216 | 0.7880216 |
| 34 | 1 | 0.6593133 | 0.09070387 | 0.4672418 | 0.8102586 |
| 105 | 1 | 0.6573749 | 0.08994986 | 0.4672669 | 0.8075790 |
| 108 | 1 | 0.6107713 | 0.10654525 | 0.3946381 | 0.7906700 |
| 31 | 1 | 0.6077246 | 0.10630367 | 0.3926429 | 0.7878028 |
| 110 | 1 | 0.6665451 | 0.09516585 | 0.4634169 | 0.8222687 |
| 103 | 1 | 0.6640844 | 0.09457498 | 0.4627210 | 0.8194301 |
| 4 | 1 | 0.5636037 | 0.17628784 | 0.2406713 | 0.8403191 |
| 19 | 1 | 0.6179444 | 0.09999704 | 0.4135466 | 0.7876783 |
| 96 | 1 | 0.6182233 | 0.10019957 | 0.4133691 | 0.7881955 |
| 74 | 1 | 0.6181870 | 0.10047501 | 0.4127846 | 0.7885464 |
| 102 | 1 | 0.6178161 | 0.10004562 | 0.4133428 | 0.7876371 |
| 76 | 1 | 0.6102189 | 0.10458972 | 0.3980455 | 0.7875266 |
| 21 | 1 | 0.6104801 | 0.10473404 | 0.3979735 | 0.7879441 |
| 37 | 1 | 0.6577129 | 0.08888403 | 0.4698591 | 0.8064253 |
| 72 | 1 | 0.6174517 | 0.10066545 | 0.4117955 | 0.7881884 |
| 14 | 1 | 0.6578789 | 0.08860594 | 0.4706011 | 0.8061903 |
| 104 | 1 | 0.6090112 | 0.10840867 | 0.3895590 | 0.7917459 |
| 17 | 1 | 0.6550491 | 0.09183381 | 0.4612643 | 0.8081246 |
| 60 | 1 | 0.6548812 | 0.09065665 | 0.4636629 | 0.8063922 |
| 52 | 1 | 0.5544714 | 0.16795672 | 0.2471597 | 0.8251052 |
| 91 | 1 | 0.5547709 | 0.16799496 | 0.2472960 | 0.8253493 |
| 93 | 1 | 0.6584761 | 0.08923313 | 0.4697392 | 0.8075568 |
| 84 | 1 | 0.5171142 | 0.17741178 | 0.2101485 | 0.8116852 |
| 20 | 1 | 0.6196895 | 0.10095781 | 0.4130489 | 0.7904826 |
| 13 | 2 | 0.6850851 | 0.07399047 | 0.5262408 | 0.8099094 |
| 77 | 2 | 0.6848084 | 0.07373020 | 0.5265898 | 0.8092983 |
| 82 | 2 | 0.6357081 | 0.08790344 | 0.4533433 | 0.7859593 |
| 34 | 2 | 0.6856001 | 0.07584362 | 0.5224854 | 0.8129441 |
| 105 | 2 | 0.6840745 | 0.07554898 | 0.5218376 | 0.8111830 |
| 108 | 2 | 0.5811077 | 0.14420948 | 0.3028394 | 0.8158467 |
| 31 | 2 | 0.6390753 | 0.08645297 | 0.4592751 | 0.7868376 |
| 110 | 2 | 0.6385218 | 0.08566092 | 0.4604713 | 0.7852207 |
| 103 | 2 | 0.6386562 | 0.08568469 | 0.4605367 | 0.7853727 |
| 4 | 2 | 0.6495617 | 0.07640387 | 0.4898154 | 0.7815923 |
| 19 | 2 | 0.6902545 | 0.07903039 | 0.5191972 | 0.8213898 |
| 96 | 2 | 0.6908631 | 0.07927316 | 0.5191556 | 0.8222483 |
| 74 | 2 | 0.6852666 | 0.07444332 | 0.5253723 | 0.8107035 |
| 102 | 2 | 0.6384031 | 0.08573007 | 0.4602234 | 0.7852155 |
| 76 | 2 | 0.6846680 | 0.07379814 | 0.5263140 | 0.8092683 |
| 21 | 2 | 0.6849245 | 0.07377575 | 0.5265875 | 0.8094656 |
| 37 | 2 | 0.6488868 | 0.07569628 | 0.4907414 | 0.7799436 |
| 72 | 2 | 0.6842157 | 0.07492269 | 0.5233768 | 0.8104370 |
| 14 | 2 | 0.6845738 | 0.07391896 | 0.5259588 | 0.8093536 |
| 104 | 2 | 0.6479504 | 0.08066409 | 0.4792527 | 0.7863598 |
| 17 | 2 | 0.5818100 | 0.14120129 | 0.3084787 | 0.8127010 |
| 60 | 2 | 0.5819790 | 0.14079864 | 0.3092919 | 0.8123324 |
| 52 | 2 | 0.6839086 | 0.07460672 | 0.5238256 | 0.8097229 |
| 91 | 2 | 0.6821565 | 0.07527800 | 0.5207917 | 0.8091020 |
| 93 | 2 | 0.6899949 | 0.07944265 | 0.5180346 | 0.8217167 |
| 84 | 2 | 0.6476976 | 0.07687208 | 0.4871661 | 0.7806084 |
| 20 | 2 | 0.5436631 | 0.15120193 | 0.2651419 | 0.7973170 |
| 13 | 3 | 0.6612485 | 0.09463816 | 0.4602814 | 0.8171170 |
| 77 | 3 | 0.6717440 | 0.08137522 | 0.4981937 | 0.8083606 |
| 82 | 3 | 0.7049854 | 0.08450141 | 0.5186989 | 0.8412387 |
| 34 | 3 | 0.6750897 | 0.08051471 | 0.5029625 | 0.8101127 |
| 105 | 3 | 0.6735736 | 0.08009725 | 0.5025994 | 0.8082049 |
| 108 | 3 | 0.6750897 | 0.08051471 | 0.5029625 | 0.8101127 |
| 31 | 3 | 0.7136022 | 0.08631622 | 0.5212782 | 0.8507789 |
| 110 | 3 | 0.6738524 | 0.07831477 | 0.5068114 | 0.8059765 |
| 103 | 3 | 0.6640675 | 0.09159564 | 0.4692210 | 0.8155101 |
| 4 | 3 | 0.7078805 | 0.08340736 | 0.5236218 | 0.8423299 |
| 19 | 3 | 0.6639432 | 0.09159961 | 0.4691101 | 0.8154094 |
| 96 | 3 | 0.6638182 | 0.09162280 | 0.4689566 | 0.8153335 |
| 74 | 3 | 0.6721994 | 0.08204346 | 0.4970956 | 0.8096775 |
| 102 | 3 | 0.7152683 | 0.08495393 | 0.5258612 | 0.8505198 |
| 76 | 3 | 0.6742263 | 0.07836521 | 0.5070208 | 0.8063773 |
| 21 | 3 | 0.7156140 | 0.08472987 | 0.5266733 | 0.8505374 |
| 37 | 3 | 0.5618118 | 0.13282780 | 0.3081049 | 0.7868483 |
| 72 | 3 | 0.5616128 | 0.13293237 | 0.3077777 | 0.7868346 |
| 14 | 3 | 0.5724889 | 0.13054674 | 0.3200748 | 0.7920710 |
| 104 | 3 | 0.6188269 | 0.13261730 | 0.3503810 | 0.8301244 |
| 17 | 3 | 0.5716302 | 0.13158672 | 0.3176198 | 0.7927763 |
| 60 | 3 | 0.5714651 | 0.13072205 | 0.3189972 | 0.7915092 |
| 52 | 3 | 0.5718163 | 0.13022350 | 0.3201301 | 0.7911225 |
| 91 | 3 | 0.5724542 | 0.13066730 | 0.3198385 | 0.7922031 |
| 93 | 3 | 0.5719544 | 0.13017707 | 0.3203154 | 0.7911683 |
| 84 | 3 | 0.7060487 | 0.08485083 | 0.5187342 | 0.8425823 |
| 20 | 3 | 0.7059934 | 0.08512804 | 0.5180361 | 0.8428822 |
| 13 | 4 | 0.6448947 | 0.13123041 | 0.3713396 | 0.8481056 |
| 77 | 4 | 0.6958997 | 0.09756883 | 0.4810609 | 0.8496027 |
| 82 | 4 | 0.7258153 | 0.09988890 | 0.4974296 | 0.8762371 |
| 34 | 4 | 0.6967362 | 0.09861051 | 0.4792859 | 0.8515122 |
| 105 | 4 | 0.6951814 | 0.09882584 | 0.4776117 | 0.8504996 |
| 108 | 4 | 0.6967362 | 0.09861051 | 0.4792859 | 0.8515122 |
| 31 | 4 | 0.7260938 | 0.10012221 | 0.4970491 | 0.8767050 |
| 110 | 4 | 0.7227823 | 0.10193258 | 0.4903060 | 0.8760333 |
| 103 | 4 | 0.7315800 | 0.10165273 | 0.4970160 | 0.8825957 |
| 4 | 4 | 0.6961539 | 0.09812983 | 0.4799570 | 0.8504728 |
| 19 | 4 | 0.7333847 | 0.10048323 | 0.5011606 | 0.8827863 |
| 96 | 4 | 0.7341469 | 0.10024020 | 0.5022864 | 0.8831274 |
| 74 | 4 | 0.7233398 | 0.10213690 | 0.4901919 | 0.8766860 |
| 102 | 4 | 0.7254471 | 0.10009277 | 0.4966709 | 0.8761651 |
| 76 | 4 | 0.7333537 | 0.10051297 | 0.5010652 | 0.8827931 |
| 21 | 4 | 0.7257873 | 0.09988383 | 0.4974225 | 0.8762096 |
| 37 | 4 | 0.7228431 | 0.10237428 | 0.4892682 | 0.8765492 |
| 72 | 4 | 0.7250366 | 0.10104992 | 0.4940320 | 0.8768618 |
| 14 | 4 | 0.5976809 | 0.12795206 | 0.3436390 | 0.8082597 |
| 104 | 4 | 0.7243843 | 0.10391777 | 0.4865439 | 0.8793689 |
| 17 | 4 | 0.6949109 | 0.09989632 | 0.4749384 | 0.8515348 |
| 60 | 4 | 0.6950728 | 0.09921712 | 0.4766268 | 0.8508709 |
| 52 | 4 | 0.6928489 | 0.10097083 | 0.4709167 | 0.8511195 |
| 91 | 4 | 0.6008077 | 0.12534076 | 0.3508151 | 0.8073872 |
| 93 | 4 | 0.6006432 | 0.12531462 | 0.3507398 | 0.8072252 |
| 84 | 4 | 0.6463596 | 0.12934616 | 0.3761290 | 0.8471165 |
| 20 | 4 | 0.7329422 | 0.10249459 | 0.4958257 | 0.8845152 |

Table S4: QAICc table of *Sholicola albiventris* for single-season, single-species models.

| **Model** | **QAICc** | **neg2ll** | **n. par** | **warn.conv** | **DQAICc** | **modlike** | **wgt** |
| --- | --- | --- | --- | --- | --- | --- | --- |
| psi(Elevation)p(Neutral) | 89.69796 | 82.1244 | 4 | 7.00 | 0.0000 | 1.0000 | 0.2715 |
| psi(Elevation)p(Neutral+survey) | 90.74626 | 80.0778 | 5 | 7.00 | 1.0483 | 0.5921 | 0.1608 |
| psi(Lantana camara)p(Neutral) | 90.87411 | 83.3336 | 4 | 5.13 | 1.1762 | 0.5554 | 0.1508 |
| psi(Elevation)p(survey) | 91.72247 | 84.2058 | 4 | 7.00 | 2.0245 | 0.3634 | 0.0987 |
| psi(Lantana camara)p(Neutral+survey) | 91.88447 | 81.2480 | 5 | 7.00 | 2.1865 | 0.3351 | 0.0910 |
| psi(Elevation)p(1) | 92.77570 | 88.1413 | 3 | 7.00 | 3.0777 | 0.2146 | 0.0583 |
| psi(Elevation)p(Warm) | 93.68045 | 86.2188 | 4 | 7.00 | 3.9825 | 0.1365 | 0.0371 |
| psi(Elevation)p(Neutral+survey+Warm) | 93.85558 | 79.8377 | 6 | 7.00 | 4.1576 | 0.1251 | 0.0340 |
| psi(Elevation)p(Neutral+survey+Cool) | 93.85791 | 79.8401 | 6 | 7.00 | 4.1600 | 0.1249 | 0.0339 |
| psi(Lantana camara)p(1) | 93.86392 | 89.2601 | 3 | 3.83 | 4.1660 | 0.1246 | 0.0338 |
| psi(Lantana camara)p(Neutral+survey+Cool) | 95.07472 | 81.0911 | 6 | 3.19 | 5.3768 | 0.0680 | 0.0185 |
| psi(Understorey height)p(Neutral) | 95.98033 | 88.5833 | 4 | 7.00 | 6.2824 | 0.0432 | 0.0117 |

Table S5: Model-averaged psi estimates and precision parameters for *S. albiventris*.

| **Site IDs** | **Estimate** | **Standard Error** | **Lower 95% Confidence Interval** | **Upper 95% Confidence Interval** |
| --- | --- | --- | --- | --- |
| 13 | 0.1543161 | 0.12847360 | 0.0258153297546 | 0.5568402 |
| 77 | 0.2281509 | 0.11453063 | 0.0763149221861 | 0.5139812 |
| 82 | 0.1486942 | 0.12975882 | 0.0228878267956 | 0.5656778 |
| 34 | 0.2260152 | 0.11958921 | 0.0710387052551 | 0.5272088 |
| 105 | 0.3089538 | 0.11566508 | 0.1339086993142 | 0.5638501 |
| 108 | 0.2910372 | 0.11341591 | 0.1226377213486 | 0.5466096 |
| 31 | 0.1453508 | 0.13053703 | 0.0212252342562 | 0.5715135 |
| 110 | 0.1603117 | 0.13317619 | 0.0267274984931 | 0.5703183 |
| 103 | 0.1924178 | 0.12045528 | 0.0495633032692 | 0.5212169 |
| 4 | 0.4822811 | 0.18371202 | 0.1804958147780 | 0.7975710 |
| 19 | 0.3424049 | 0.12128173 | 0.1533835487157 | 0.5994359 |
| 96 | 0.2727003 | 0.11466110 | 0.1077379939584 | 0.5379585 |
| 74 | 0.4749668 | 0.18345873 | 0.1762271642560 | 0.7927674 |
| 102 | 0.5770305 | 0.23207354 | 0.1746445217755 | 0.8979134 |
| 76 | 0.2935113 | 0.11209748 | 0.1258762862853 | 0.5451617 |
| 21 | 0.5645221 | 0.22916147 | 0.1725757131392 | 0.8895893 |
| 37 | 0.1358074 | 0.13923714 | 0.0151303724716 | 0.6164931 |
| 72 | 0.3120059 | 0.12370113 | 0.1278362552493 | 0.5838773 |
| 14 | 0.3394807 | 0.12236630 | 0.1499262641144 | 0.5996390 |
| 104 | 0.2801025 | 0.11448336 | 0.1133754665203 | 0.5421026 |
| 17 | 0.9397833 | 0.09461150 | 0.3707313645065 | 0.9975870 |
| 60 | 0.8696476 | 0.14870559 | 0.3377765575777 | 0.9886701 |
| 52 | 0.9686965 | 0.06225366 | 0.3562810403338 | 0.9994224 |
| 91 | 0.9971195 | 0.02940619 | 0.0000006673817 | 1.0000000 |
| 93 | 0.9971837 | 0.02940917 | 0.0000004319817 | 1.0000000 |
| 84 | 0.6000361 | 0.24309080 | 0.1708451563475 | 0.9161292 |
| 20 | 0.9635752 | 0.06857622 | 0.3649208778974 | 0.9991796 |

Table S6: Model-averaged p estimates and precision parameters for *S. albiventris*.

| **Site IDs** | **Survey no.** | **Estimate** | **Standard Error** | **Lower 95% Confidence Interval** | **Upper 95% Confidence Interval** |
| --- | --- | --- | --- | --- | --- |
| 13 | 1 | 0.6846808 | 0.1713888 | 0.3141971 | 0.9114366 |
| 77 | 1 | 0.4260318 | 0.1387154 | 0.1962495 | 0.6929148 |
| 82 | 1 | 0.4102089 | 0.1400732 | 0.1827463 | 0.6838771 |
| 34 | 1 | 0.4102089 | 0.1400732 | 0.1827463 | 0.6838771 |
| 105 | 1 | 0.6846808 | 0.1713888 | 0.3141971 | 0.9114366 |
| 108 | 1 | 0.6846808 | 0.1713888 | 0.3141971 | 0.9114366 |
| 31 | 1 | 0.4260318 | 0.1387154 | 0.1962495 | 0.6929148 |
| 110 | 1 | 0.6846808 | 0.1713888 | 0.3141971 | 0.9114366 |
| 103 | 1 | 0.4260318 | 0.1387154 | 0.1962495 | 0.6929148 |
| 4 | 1 | 0.6846808 | 0.1713888 | 0.3141971 | 0.9114366 |
| 19 | 1 | 0.4260318 | 0.1387154 | 0.1962495 | 0.6929148 |
| 96 | 1 | 0.4260318 | 0.1387154 | 0.1962495 | 0.6929148 |
| 74 | 1 | 0.4260318 | 0.1387154 | 0.1962495 | 0.6929148 |
| 102 | 1 | 0.4260318 | 0.1387154 | 0.1962495 | 0.6929148 |
| 76 | 1 | 0.4102089 | 0.1400732 | 0.1827463 | 0.6838771 |
| 21 | 1 | 0.4102089 | 0.1400732 | 0.1827463 | 0.6838771 |
| 37 | 1 | 0.6846808 | 0.1713888 | 0.3141971 | 0.9114366 |
| 72 | 1 | 0.4260318 | 0.1387154 | 0.1962495 | 0.6929148 |
| 14 | 1 | 0.6846808 | 0.1713888 | 0.3141971 | 0.9114366 |
| 104 | 1 | 0.4102089 | 0.1400732 | 0.1827463 | 0.6838771 |
| 17 | 1 | 0.4260318 | 0.1387154 | 0.1962495 | 0.6929148 |
| 60 | 1 | 0.4260318 | 0.1387154 | 0.1962495 | 0.6929148 |
| 52 | 1 | 0.4102089 | 0.1400732 | 0.1827463 | 0.6838771 |
| 91 | 1 | 0.4102089 | 0.1400732 | 0.1827463 | 0.6838771 |
| 93 | 1 | 0.4102089 | 0.1400732 | 0.1827463 | 0.6838771 |
| 84 | 1 | 0.6846808 | 0.1713888 | 0.3141971 | 0.9114366 |
| 20 | 1 | 0.6846808 | 0.1713888 | 0.3141971 | 0.9114366 |
| 13 | 2 | 0.4557572 | 0.1187463 | 0.2468031 | 0.6815415 |
| 77 | 2 | 0.4557572 | 0.1187463 | 0.2468031 | 0.6815415 |
| 82 | 2 | 0.4724848 | 0.1164311 | 0.2639094 | 0.6911292 |
| 34 | 2 | 0.4557572 | 0.1187463 | 0.2468031 | 0.6815415 |
| 105 | 2 | 0.4557572 | 0.1187463 | 0.2468031 | 0.6815415 |
| 108 | 2 | 0.4724848 | 0.1164311 | 0.2639094 | 0.6911292 |
| 31 | 2 | 0.4557572 | 0.1187463 | 0.2468031 | 0.6815415 |
| 110 | 2 | 0.4557572 | 0.1187463 | 0.2468031 | 0.6815415 |
| 103 | 2 | 0.4557572 | 0.1187463 | 0.2468031 | 0.6815415 |
| 4 | 2 | 0.4557572 | 0.1187463 | 0.2468031 | 0.6815415 |
| 19 | 2 | 0.4724848 | 0.1164311 | 0.2639094 | 0.6911292 |
| 96 | 2 | 0.4724848 | 0.1164311 | 0.2639094 | 0.6911292 |
| 74 | 2 | 0.4557572 | 0.1187463 | 0.2468031 | 0.6815415 |
| 102 | 2 | 0.4557572 | 0.1187463 | 0.2468031 | 0.6815415 |
| 76 | 2 | 0.4557572 | 0.1187463 | 0.2468031 | 0.6815415 |
| 21 | 2 | 0.4557572 | 0.1187463 | 0.2468031 | 0.6815415 |
| 37 | 2 | 0.4557572 | 0.1187463 | 0.2468031 | 0.6815415 |
| 72 | 2 | 0.4557572 | 0.1187463 | 0.2468031 | 0.6815415 |
| 14 | 2 | 0.4557572 | 0.1187463 | 0.2468031 | 0.6815415 |
| 104 | 2 | 0.4557572 | 0.1187463 | 0.2468031 | 0.6815415 |
| 17 | 2 | 0.7275809 | 0.1298488 | 0.4251607 | 0.9060549 |
| 60 | 2 | 0.7275809 | 0.1298488 | 0.4251607 | 0.9060549 |
| 52 | 2 | 0.7275809 | 0.1298488 | 0.4251607 | 0.9060549 |
| 91 | 2 | 0.4724848 | 0.1164311 | 0.2639094 | 0.6911292 |
| 93 | 2 | 0.4724848 | 0.1164311 | 0.2639094 | 0.6911292 |
| 84 | 2 | 0.7275809 | 0.1298488 | 0.4251607 | 0.9060549 |
| 20 | 2 | 0.7275809 | 0.1298488 | 0.4251607 | 0.9060549 |
| 13 | 3 | 0.5202682 | 0.1292614 | 0.2821258 | 0.7495428 |
| 77 | 3 | 0.5202682 | 0.1292614 | 0.2821258 | 0.7495428 |
| 82 | 3 | 0.5202682 | 0.1292614 | 0.2821258 | 0.7495428 |
| 34 | 3 | 0.7645736 | 0.1137778 | 0.4847633 | 0.9180999 |
| 105 | 3 | 0.7645736 | 0.1137778 | 0.4847633 | 0.9180999 |
| 108 | 3 | 0.7645736 | 0.1137778 | 0.4847633 | 0.9180999 |
| 31 | 3 | 0.5202682 | 0.1292614 | 0.2821258 | 0.7495428 |
| 110 | 3 | 0.7645736 | 0.1137778 | 0.4847633 | 0.9180999 |
| 103 | 3 | 0.5033134 | 0.1300856 | 0.2676373 | 0.7375262 |
| 4 | 3 | 0.5033134 | 0.1300856 | 0.2676373 | 0.7375262 |
| 19 | 3 | 0.5033134 | 0.1300856 | 0.2676373 | 0.7375262 |
| 96 | 3 | 0.7645736 | 0.1137778 | 0.4847633 | 0.9180999 |
| 74 | 3 | 0.5202682 | 0.1292614 | 0.2821258 | 0.7495428 |
| 102 | 3 | 0.5033134 | 0.1300856 | 0.2676373 | 0.7375262 |
| 76 | 3 | 0.5033134 | 0.1300856 | 0.2676373 | 0.7375262 |
| 21 | 3 | 0.5033134 | 0.1300856 | 0.2676373 | 0.7375262 |
| 37 | 3 | 0.5033134 | 0.1300856 | 0.2676373 | 0.7375262 |
| 72 | 3 | 0.5033134 | 0.1300856 | 0.2676373 | 0.7375262 |
| 14 | 3 | 0.5033134 | 0.1300856 | 0.2676373 | 0.7375262 |
| 104 | 3 | 0.5033134 | 0.1300856 | 0.2676373 | 0.7375262 |
| 17 | 3 | 0.7645736 | 0.1137778 | 0.4847633 | 0.9180999 |
| 60 | 3 | 0.7645736 | 0.1137778 | 0.4847633 | 0.9180999 |
| 52 | 3 | 0.7645736 | 0.1137778 | 0.4847633 | 0.9180999 |
| 91 | 3 | 0.7645736 | 0.1137778 | 0.4847633 | 0.9180999 |
| 93 | 3 | 0.7645736 | 0.1137778 | 0.4847633 | 0.9180999 |
| 84 | 3 | 0.7645736 | 0.1137778 | 0.4847633 | 0.9180999 |
| 20 | 3 | 0.7645736 | 0.1137778 | 0.4847633 | 0.9180999 |
| 13 | 4 | 0.5649269 | 0.1648618 | 0.2585578 | 0.8286149 |
| 77 | 4 | 0.7939474 | 0.1177284 | 0.4846122 | 0.9404388 |
| 82 | 4 | 0.5485009 | 0.1650898 | 0.2475059 | 0.8177518 |
| 34 | 4 | 0.5485009 | 0.1650898 | 0.2475059 | 0.8177518 |
| 105 | 4 | 0.7939474 | 0.1177284 | 0.4846122 | 0.9404388 |
| 108 | 4 | 0.5485009 | 0.1650898 | 0.2475059 | 0.8177518 |
| 31 | 4 | 0.5485009 | 0.1650898 | 0.2475059 | 0.8177518 |
| 110 | 4 | 0.5649269 | 0.1648618 | 0.2585578 | 0.8286149 |
| 103 | 4 | 0.5649269 | 0.1648618 | 0.2585578 | 0.8286149 |
| 4 | 4 | 0.7939474 | 0.1177284 | 0.4846122 | 0.9404388 |
| 19 | 4 | 0.7939474 | 0.1177284 | 0.4846122 | 0.9404388 |
| 96 | 4 | 0.7834976 | 0.1386957 | 0.4215581 | 0.9472857 |
| 74 | 4 | 0.5649269 | 0.1648618 | 0.2585578 | 0.8286149 |
| 102 | 4 | 0.5485009 | 0.1650898 | 0.2475059 | 0.8177518 |
| 76 | 4 | 0.7939474 | 0.1177284 | 0.4846122 | 0.9404388 |
| 21 | 4 | 0.5485009 | 0.1650898 | 0.2475059 | 0.8177518 |
| 37 | 4 | 0.5649269 | 0.1648618 | 0.2585578 | 0.8286149 |
| 72 | 4 | 0.7939474 | 0.1177284 | 0.4846122 | 0.9404388 |
| 14 | 4 | 0.5649269 | 0.1648618 | 0.2585578 | 0.8286149 |
| 104 | 4 | 0.5485009 | 0.1650898 | 0.2475059 | 0.8177518 |
| 17 | 4 | 0.7939474 | 0.1177284 | 0.4846122 | 0.9404388 |
| 60 | 4 | 0.7939474 | 0.1177284 | 0.4846122 | 0.9404388 |
| 52 | 4 | 0.5649269 | 0.1648618 | 0.2585578 | 0.8286149 |
| 91 | 4 | 0.5485009 | 0.1650898 | 0.2475059 | 0.8177518 |
| 93 | 4 | 0.5485009 | 0.1650898 | 0.2475059 | 0.8177518 |
| 84 | 4 | 0.7939474 | 0.1177284 | 0.4846122 | 0.9404388 |
| 20 | 4 | 0.7939474 | 0.1177284 | 0.4846122 | 0.9404388 |

Table S7. QAICc table of single-season, two-species models: MS format: Species A = *M. fairbanki*, Species B = *S. albiventris*

| **Model** | **QAICc** | **neg2ll** | **n. par** | **warn.conv** | **DQAICc** | **modlike** | **wgt** |
| --- | --- | --- | --- | --- | --- | --- | --- |
| psi(SP+Elevation)p(SP+INT_d+survey) | 174.3287 | 218.2615 | 7 | 7.00 | 0.0000 | 1.0000 | 0.1819 |
| psi(SP+Elevation)p(SP+INT_o+survey) | 174.5292 | 218.5449 | 7 | 7.00 | 0.2005 | 0.9046 | 0.1645 |
| psi(SP+Elevation)p(SP+INT_d) | 174.5551 | 223.8033 | 6 | 7.00 | 0.2264 | 0.8930 | 0.1624 |
| psi(SP+Lantana_camara)p(SP+INT_d+survey) | 175.2572 | 219.5738 | 7 | 5.78 | 0.9285 | 0.6286 | 0.1143 |
| psi(SP+Elevation)p(SP+INT_o) | 175.3010 | 224.8575 | 6 | 7.00 | 0.9724 | 0.6150 | 0.1118 |
| psi(SP+Lantana_camara)p(SP+INT_d) | 175.4827 | 225.1142 | 6 | 3.08 | 1.1540 | 0.5616 | 0.1021 |
| psi(SP+Elevation+Lantana_camara)p(SP+INT_d) | 177.1000 | 222.1782 | 7 | 5.32 | 2.7713 | 0.2502 | 0.0455 |
| psi(SP+Elevation+No_road)p(SP+INT_d+BefNoon) | 177.1154 | 216.3980 | 8 | 7.00 | 2.7867 | 0.2482 | 0.0451 |
| psi(SP+Elevation+Area)p(SP+INT_d+BefNoon) | 177.5008 | 216.9427 | 8 | 7.00 | 3.1721 | 0.2047 | 0.0372 |
| psi(SP+Elevation)p(SP+INT_d+BefNoon) | 177.6182 | 222.9106 | 7 | 7.00 | 3.2895 | 0.1931 | 0.0351 |

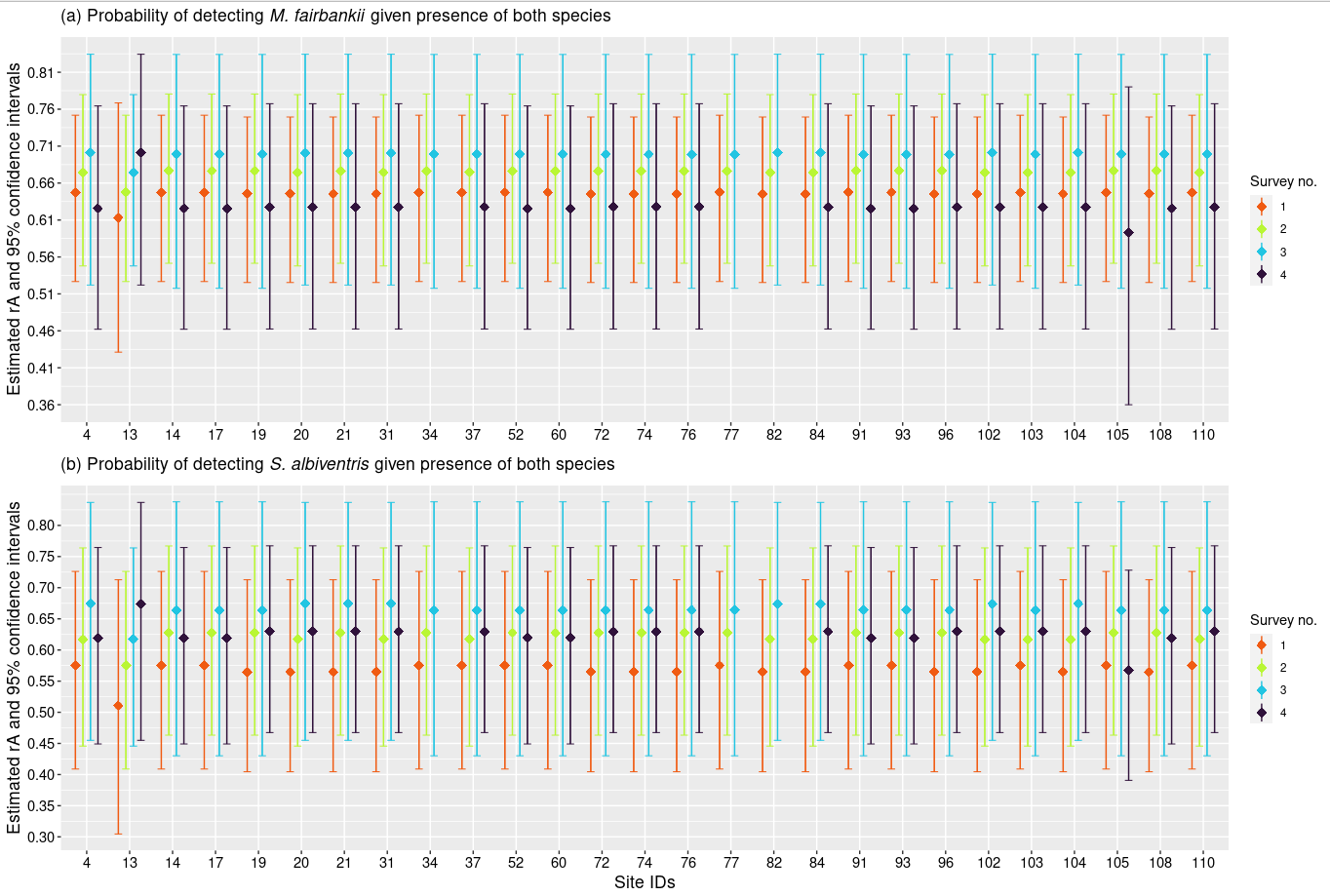

Figure S9: Model-averaged rA estimates (det. prob. of *A*, given both species are present) and 95% confidence intervals for Two-species models.

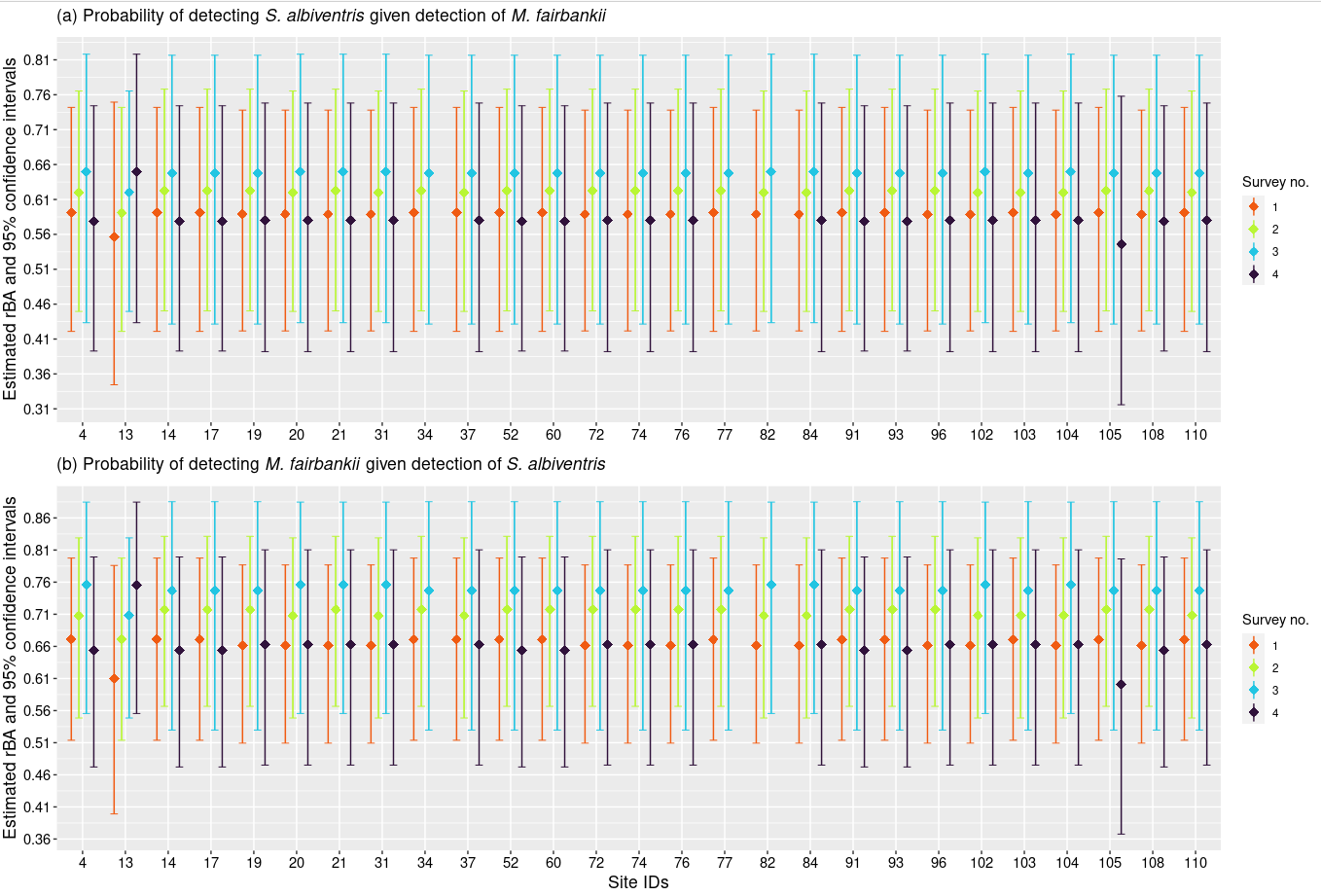

Figure S10: Model-averaged rBA estimates (det. prob. of *A*, given *B* was detected) and 95% confidence intervals for Two-species models.

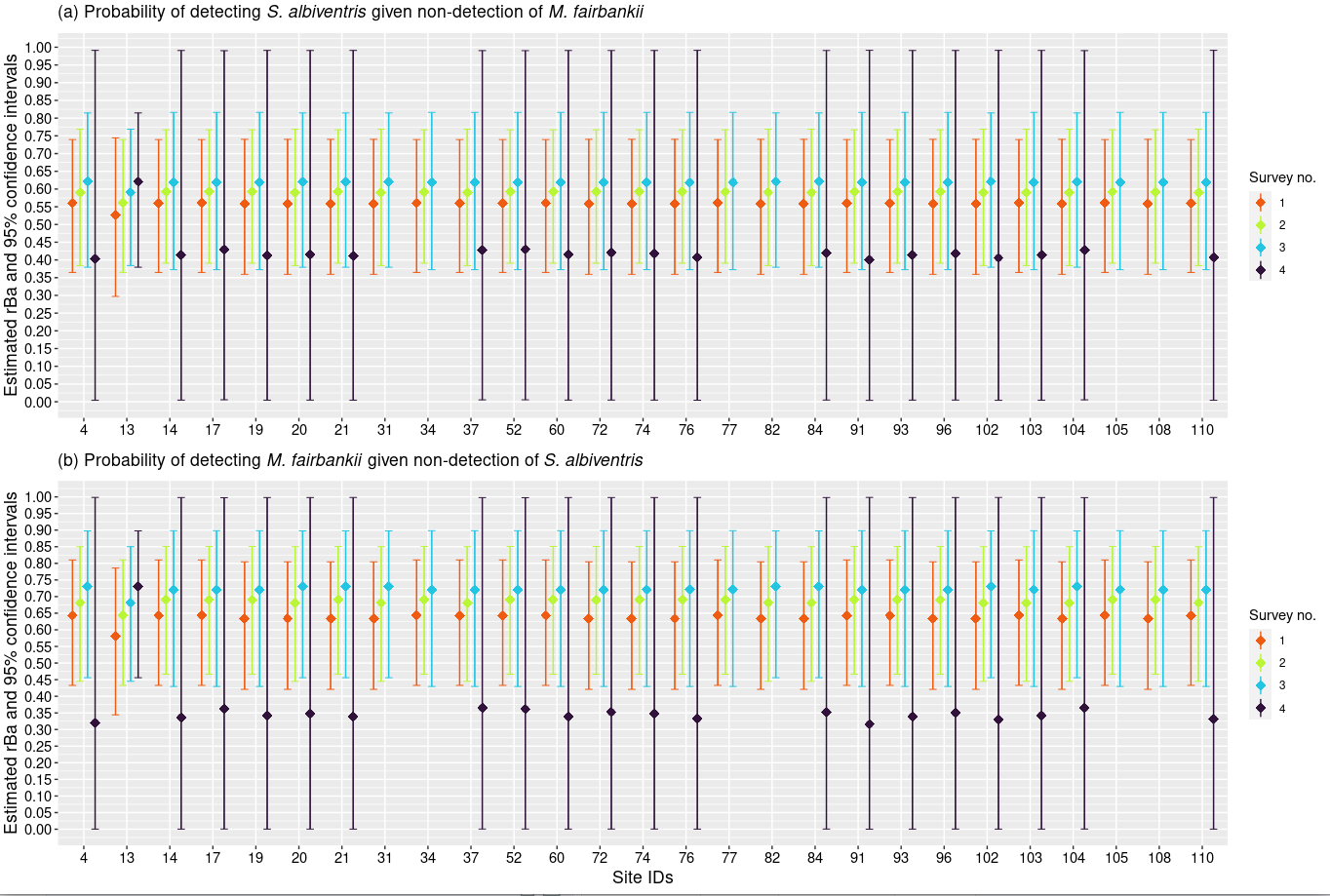

Figure S11: Model-averaged rBa estimates (det. prob. of *A*, given *B* was not detected) and 95% confidence intervals for Two-species models.

Table S8: Model-averaged r^M^ estimates (det. prob. of *M. fairbakii*, given both species are present) and precision parameters for Two-species models (MS format).

| **Site IDs** | **Survey no.** | **Estimate** | **Standard Error** | **Lower 95% Confidence Interval** | **Upper 95% Confidence Interval** |
| --- | --- | --- | --- | --- | --- |
| 13 | 1 | 0.6014822 | 0.09526542 | 0.4091909 | 0.7668485 |
| 77 | 1 | 0.6536526 | 0.06591573 | 0.5161126 | 0.7695541 |
| 82 | 1 | 0.6447922 | 0.06530027 | 0.5093538 | 0.7604286 |
| 34 | 1 | 0.6536526 | 0.06591573 | 0.5161126 | 0.7695541 |
| 105 | 1 | 0.6536526 | 0.06591573 | 0.5161126 | 0.7695541 |
| 108 | 1 | 0.6447922 | 0.06530027 | 0.5093538 | 0.7604286 |
| 31 | 1 | 0.6447922 | 0.06530027 | 0.5093538 | 0.7604286 |
| 110 | 1 | 0.6536526 | 0.06591573 | 0.5161126 | 0.7695541 |
| 103 | 1 | 0.6536526 | 0.06591573 | 0.5161126 | 0.7695541 |
| 4 | 1 | 0.6536526 | 0.06591573 | 0.5161126 | 0.7695541 |
| 19 | 1 | 0.6447922 | 0.06530027 | 0.5093538 | 0.7604286 |
| 96 | 1 | 0.6447922 | 0.06530027 | 0.5093538 | 0.7604286 |
| 74 | 1 | 0.6447922 | 0.06530027 | 0.5093538 | 0.7604286 |
| 102 | 1 | 0.6447922 | 0.06530027 | 0.5093538 | 0.7604286 |
| 76 | 1 | 0.6447922 | 0.06530027 | 0.5093538 | 0.7604286 |
| 21 | 1 | 0.6447922 | 0.06530027 | 0.5093538 | 0.7604286 |
| 37 | 1 | 0.6536526 | 0.06591573 | 0.5161126 | 0.7695541 |
| 72 | 1 | 0.6447922 | 0.06530027 | 0.5093538 | 0.7604286 |
| 14 | 1 | 0.6536526 | 0.06591573 | 0.5161126 | 0.7695541 |
| 104 | 1 | 0.6447922 | 0.06530027 | 0.5093538 | 0.7604286 |
| 17 | 1 | 0.6536526 | 0.06591573 | 0.5161126 | 0.7695541 |
| 60 | 1 | 0.6536526 | 0.06591573 | 0.5161126 | 0.7695541 |
| 52 | 1 | 0.6536526 | 0.06591573 | 0.5161126 | 0.7695541 |
| 91 | 1 | 0.6536526 | 0.06591573 | 0.5161126 | 0.7695541 |
| 93 | 1 | 0.6536526 | 0.06591573 | 0.5161126 | 0.7695541 |
| 84 | 1 | 0.6447922 | 0.06530027 | 0.5093538 | 0.7604286 |
| 20 | 1 | 0.6447922 | 0.06530027 | 0.5093538 | 0.7604286 |
| 13 | 2 | 0.6536526 | 0.06591573 | 0.5161126 | 0.7695541 |
| 77 | 2 | 0.6927212 | 0.06471453 | 0.5540362 | 0.8035690 |
| 82 | 2 | 0.6838607 | 0.06927871 | 0.5358162 | 0.8021256 |
| 34 | 2 | 0.6927212 | 0.06471453 | 0.5540362 | 0.8035690 |
| 105 | 2 | 0.6927212 | 0.06471453 | 0.5540362 | 0.8035690 |
| 108 | 2 | 0.6927212 | 0.06471453 | 0.5540362 | 0.8035690 |
| 31 | 2 | 0.6838607 | 0.06927871 | 0.5358162 | 0.8021256 |
| 110 | 2 | 0.6838607 | 0.06927871 | 0.5358162 | 0.8021256 |
| 103 | 2 | 0.6838607 | 0.06927871 | 0.5358162 | 0.8021256 |
| 4 | 2 | 0.6838607 | 0.06927871 | 0.5358162 | 0.8021256 |
| 19 | 2 | 0.6927212 | 0.06471453 | 0.5540362 | 0.8035690 |
| 96 | 2 | 0.6927212 | 0.06471453 | 0.5540362 | 0.8035690 |
| 74 | 2 | 0.6927212 | 0.06471453 | 0.5540362 | 0.8035690 |
| 102 | 2 | 0.6838607 | 0.06927871 | 0.5358162 | 0.8021256 |
| 76 | 2 | 0.6927212 | 0.06471453 | 0.5540362 | 0.8035690 |
| 21 | 2 | 0.6927212 | 0.06471453 | 0.5540362 | 0.8035690 |
| 37 | 2 | 0.6838607 | 0.06927871 | 0.5358162 | 0.8021256 |
| 72 | 2 | 0.6927212 | 0.06471453 | 0.5540362 | 0.8035690 |
| 14 | 2 | 0.6927212 | 0.06471453 | 0.5540362 | 0.8035690 |
| 104 | 2 | 0.6838607 | 0.06927871 | 0.5358162 | 0.8021256 |
| 17 | 2 | 0.6927212 | 0.06471453 | 0.5540362 | 0.8035690 |
| 60 | 2 | 0.6927212 | 0.06471453 | 0.5540362 | 0.8035690 |
| 52 | 2 | 0.6927212 | 0.06471453 | 0.5540362 | 0.8035690 |
| 91 | 2 | 0.6927212 | 0.06471453 | 0.5540362 | 0.8035690 |
| 93 | 2 | 0.6927212 | 0.06471453 | 0.5540362 | 0.8035690 |
| 84 | 2 | 0.6838607 | 0.06927871 | 0.5358162 | 0.8021256 |
| 20 | 2 | 0.6838607 | 0.06927871 | 0.5358162 | 0.8021256 |
| 13 | 3 | 0.6838607 | 0.06927871 | 0.5358162 | 0.8021256 |
| 77 | 3 | 0.7169307 | 0.09253302 | 0.5089046 | 0.8609200 |
| 82 | 3 | 0.7257911 | 0.08581819 | 0.5320115 | 0.8603888 |
| 34 | 3 | 0.7169307 | 0.09253302 | 0.5089046 | 0.8609200 |
| 105 | 3 | 0.7169307 | 0.09253302 | 0.5089046 | 0.8609200 |
| 108 | 3 | 0.7169307 | 0.09253302 | 0.5089046 | 0.8609200 |
| 31 | 3 | 0.7257911 | 0.08581819 | 0.5320115 | 0.8603888 |
| 110 | 3 | 0.7169307 | 0.09253302 | 0.5089046 | 0.8609200 |
| 103 | 3 | 0.7169307 | 0.09253302 | 0.5089046 | 0.8609200 |
| 4 | 3 | 0.7257911 | 0.08581819 | 0.5320115 | 0.8603888 |
| 19 | 3 | 0.7169307 | 0.09253302 | 0.5089046 | 0.8609200 |
| 96 | 3 | 0.7169307 | 0.09253302 | 0.5089046 | 0.8609200 |
| 74 | 3 | 0.7169307 | 0.09253302 | 0.5089046 | 0.8609200 |
| 102 | 3 | 0.7257911 | 0.08581819 | 0.5320115 | 0.8603888 |
| 76 | 3 | 0.7169307 | 0.09253302 | 0.5089046 | 0.8609200 |
| 21 | 3 | 0.7257911 | 0.08581819 | 0.5320115 | 0.8603888 |
| 37 | 3 | 0.7169307 | 0.09253302 | 0.5089046 | 0.8609200 |
| 72 | 3 | 0.7169307 | 0.09253302 | 0.5089046 | 0.8609200 |
| 14 | 3 | 0.7169307 | 0.09253302 | 0.5089046 | 0.8609200 |
| 104 | 3 | 0.7257911 | 0.08581819 | 0.5320115 | 0.8603888 |
| 17 | 3 | 0.7169307 | 0.09253302 | 0.5089046 | 0.8609200 |
| 60 | 3 | 0.7169307 | 0.09253302 | 0.5089046 | 0.8609200 |
| 52 | 3 | 0.7169307 | 0.09253302 | 0.5089046 | 0.8609200 |
| 91 | 3 | 0.7169307 | 0.09253302 | 0.5089046 | 0.8609200 |
| 93 | 3 | 0.7169307 | 0.09253302 | 0.5089046 | 0.8609200 |
| 84 | 3 | 0.7257911 | 0.08581819 | 0.5320115 | 0.8603888 |
| 20 | 3 | 0.7257911 | 0.08581819 | 0.5320115 | 0.8603888 |
| 13 | 4 | 0.7257911 | 0.08581819 | 0.5320115 | 0.8603888 |
| 77 | 4 | NA | NA | NA | NA |
| 82 | 4 | NA | NA | NA | NA |
| 34 | 4 | NA | NA | NA | NA |
| 105 | 4 | 0.5784086 | 0.12225747 | 0.3393030 | 0.7856481 |
| 108 | 4 | 0.6227173 | 0.08614293 | 0.4458285 | 0.7720170 |
| 31 | 4 | 0.6315777 | 0.08884002 | 0.4478595 | 0.7836908 |
| 110 | 4 | 0.6315777 | 0.08884002 | 0.4478595 | 0.7836908 |
| 103 | 4 | 0.6315777 | 0.08884002 | 0.4478595 | 0.7836908 |
| 4 | 4 | 0.6227173 | 0.08614293 | 0.4458285 | 0.7720170 |
| 19 | 4 | 0.6315777 | 0.08884002 | 0.4478595 | 0.7836908 |
| 96 | 4 | 0.6315777 | 0.08884002 | 0.4478595 | 0.7836908 |
| 74 | 4 | 0.6315777 | 0.08884002 | 0.4478595 | 0.7836908 |
| 102 | 4 | 0.6315777 | 0.08884002 | 0.4478595 | 0.7836908 |
| 76 | 4 | 0.6315777 | 0.08884002 | 0.4478595 | 0.7836908 |
| 21 | 4 | 0.6315777 | 0.08884002 | 0.4478595 | 0.7836908 |
| 37 | 4 | 0.6315777 | 0.08884002 | 0.4478595 | 0.7836908 |
| 72 | 4 | 0.6315777 | 0.08884002 | 0.4478595 | 0.7836908 |
| 14 | 4 | 0.6227173 | 0.08614293 | 0.4458285 | 0.7720170 |
| 104 | 4 | 0.6315777 | 0.08884002 | 0.4478595 | 0.7836908 |
| 17 | 4 | 0.6227173 | 0.08614293 | 0.4458285 | 0.7720170 |
| 60 | 4 | 0.6227173 | 0.08614293 | 0.4458285 | 0.7720170 |
| 52 | 4 | 0.6227173 | 0.08614293 | 0.4458285 | 0.7720170 |
| 91 | 4 | 0.6227173 | 0.08614293 | 0.4458285 | 0.7720170 |
| 93 | 4 | 0.6227173 | 0.08614293 | 0.4458285 | 0.7720170 |
| 84 | 4 | 0.6315777 | 0.08884002 | 0.4478595 | 0.7836908 |
| 20 | 4 | 0.6315777 | 0.08884002 | 0.4478595 | 0.7836908 |

Table S9: Model-averaged r^SM^ estimates (det. prob. of *S. albiventris*, given *M. fairbanki* was detected) and precision parameters for Two-species models (MS format).

| **Site IDs** | **Survey no.** | **Estimate** | **Standard Error** | **Lower 95% Confidence Interval** | **Upper 95% Confidence Interval** |
| --- | --- | --- | --- | --- | --- |
| 13 | 1 | 0.5680243 | 0.12545932 | 0.3255598 | 0.7817556 |
| 77 | 1 | 0.6212138 | 0.09786822 | 0.4205591 | 0.7874940 |
| 82 | 1 | 0.6123330 | 0.09557189 | 0.4177614 | 0.7766464 |
| 34 | 1 | 0.6212138 | 0.09786822 | 0.4205591 | 0.7874940 |
| 105 | 1 | 0.6212138 | 0.09786822 | 0.4205591 | 0.7874940 |
| 108 | 1 | 0.6123330 | 0.09557189 | 0.4177614 | 0.7766464 |
| 31 | 1 | 0.6123330 | 0.09557189 | 0.4177614 | 0.7766464 |
| 110 | 1 | 0.6212138 | 0.09786822 | 0.4205591 | 0.7874940 |
| 103 | 1 | 0.6212138 | 0.09786822 | 0.4205591 | 0.7874940 |
| 4 | 1 | 0.6212138 | 0.09786822 | 0.4205591 | 0.7874940 |
| 19 | 1 | 0.6123330 | 0.09557189 | 0.4177614 | 0.7766464 |
| 96 | 1 | 0.6123330 | 0.09557189 | 0.4177614 | 0.7766464 |
| 74 | 1 | 0.6123330 | 0.09557189 | 0.4177614 | 0.7766464 |
| 102 | 1 | 0.6123330 | 0.09557189 | 0.4177614 | 0.7766464 |
| 76 | 1 | 0.6123330 | 0.09557189 | 0.4177614 | 0.7766464 |
| 21 | 1 | 0.6123330 | 0.09557189 | 0.4177614 | 0.7766464 |
| 37 | 1 | 0.6212138 | 0.09786822 | 0.4205591 | 0.7874940 |
| 72 | 1 | 0.6123330 | 0.09557189 | 0.4177614 | 0.7766464 |
| 14 | 1 | 0.6212138 | 0.09786822 | 0.4205591 | 0.7874940 |
| 104 | 1 | 0.6123330 | 0.09557189 | 0.4177614 | 0.7766464 |
| 17 | 1 | 0.6212138 | 0.09786822 | 0.4205591 | 0.7874940 |
| 60 | 1 | 0.6212138 | 0.09786822 | 0.4205591 | 0.7874940 |
| 52 | 1 | 0.6212138 | 0.09786822 | 0.4205591 | 0.7874940 |
| 91 | 1 | 0.6212138 | 0.09786822 | 0.4205591 | 0.7874940 |
| 93 | 1 | 0.6212138 | 0.09786822 | 0.4205591 | 0.7874940 |
| 84 | 1 | 0.6123330 | 0.09557189 | 0.4177614 | 0.7766464 |
| 20 | 1 | 0.6123330 | 0.09557189 | 0.4177614 | 0.7766464 |
| 13 | 2 | 0.6212138 | 0.09786822 | 0.4205591 | 0.7874940 |
| 77 | 2 | 0.6627764 | 0.08823564 | 0.4755021 | 0.8099142 |
| 82 | 2 | 0.6538955 | 0.08988630 | 0.4645012 | 0.8044981 |
| 34 | 2 | 0.6627764 | 0.08823564 | 0.4755021 | 0.8099142 |
| 105 | 2 | 0.6627764 | 0.08823564 | 0.4755021 | 0.8099142 |
| 108 | 2 | 0.6627764 | 0.08823564 | 0.4755021 | 0.8099142 |
| 31 | 2 | 0.6538955 | 0.08988630 | 0.4645012 | 0.8044981 |
| 110 | 2 | 0.6538955 | 0.08988630 | 0.4645012 | 0.8044981 |
| 103 | 2 | 0.6538955 | 0.08988630 | 0.4645012 | 0.8044981 |
| 4 | 2 | 0.6538955 | 0.08988630 | 0.4645012 | 0.8044981 |
| 19 | 2 | 0.6627764 | 0.08823564 | 0.4755021 | 0.8099142 |
| 96 | 2 | 0.6627764 | 0.08823564 | 0.4755021 | 0.8099142 |
| 74 | 2 | 0.6627764 | 0.08823564 | 0.4755021 | 0.8099142 |
| 102 | 2 | 0.6538955 | 0.08988630 | 0.4645012 | 0.8044981 |
| 76 | 2 | 0.6627764 | 0.08823564 | 0.4755021 | 0.8099142 |
| 21 | 2 | 0.6627764 | 0.08823564 | 0.4755021 | 0.8099142 |
| 37 | 2 | 0.6538955 | 0.08988630 | 0.4645012 | 0.8044981 |
| 72 | 2 | 0.6627764 | 0.08823564 | 0.4755021 | 0.8099142 |
| 14 | 2 | 0.6627764 | 0.08823564 | 0.4755021 | 0.8099142 |
| 104 | 2 | 0.6538955 | 0.08988630 | 0.4645012 | 0.8044981 |
| 17 | 2 | 0.6627764 | 0.08823564 | 0.4755021 | 0.8099142 |
| 60 | 2 | 0.6627764 | 0.08823564 | 0.4755021 | 0.8099142 |
| 52 | 2 | 0.6627764 | 0.08823564 | 0.4755021 | 0.8099142 |
| 91 | 2 | 0.6627764 | 0.08823564 | 0.4755021 | 0.8099142 |
| 93 | 2 | 0.6627764 | 0.08823564 | 0.4755021 | 0.8099142 |
| 84 | 2 | 0.6538955 | 0.08988630 | 0.4645012 | 0.8044981 |
| 20 | 2 | 0.6538955 | 0.08988630 | 0.4645012 | 0.8044981 |
| 13 | 3 | 0.6538955 | 0.08988630 | 0.4645012 | 0.8044981 |
| 77 | 3 | 0.6902559 | 0.10521831 | 0.4592811 | 0.8539432 |
| 82 | 3 | 0.6991368 | 0.10065312 | 0.4763471 | 0.8558282 |
| 34 | 3 | 0.6902559 | 0.10521831 | 0.4592811 | 0.8539432 |
| 105 | 3 | 0.6902559 | 0.10521831 | 0.4592811 | 0.8539432 |
| 108 | 3 | 0.6902559 | 0.10521831 | 0.4592811 | 0.8539432 |
| 31 | 3 | 0.6991368 | 0.10065312 | 0.4763471 | 0.8558282 |
| 110 | 3 | 0.6902559 | 0.10521831 | 0.4592811 | 0.8539432 |
| 103 | 3 | 0.6902559 | 0.10521831 | 0.4592811 | 0.8539432 |
| 4 | 3 | 0.6991368 | 0.10065312 | 0.4763471 | 0.8558282 |
| 19 | 3 | 0.6902559 | 0.10521831 | 0.4592811 | 0.8539432 |
| 96 | 3 | 0.6902559 | 0.10521831 | 0.4592811 | 0.8539432 |
| 74 | 3 | 0.6902559 | 0.10521831 | 0.4592811 | 0.8539432 |
| 102 | 3 | 0.6991368 | 0.10065312 | 0.4763471 | 0.8558282 |
| 76 | 3 | 0.6902559 | 0.10521831 | 0.4592811 | 0.8539432 |
| 21 | 3 | 0.6991368 | 0.10065312 | 0.4763471 | 0.8558282 |
| 37 | 3 | 0.6902559 | 0.10521831 | 0.4592811 | 0.8539432 |
| 72 | 3 | 0.6902559 | 0.10521831 | 0.4592811 | 0.8539432 |
| 14 | 3 | 0.6902559 | 0.10521831 | 0.4592811 | 0.8539432 |
| 104 | 3 | 0.6991368 | 0.10065312 | 0.4763471 | 0.8558282 |
| 17 | 3 | 0.6902559 | 0.10521831 | 0.4592811 | 0.8539432 |
| 60 | 3 | 0.6902559 | 0.10521831 | 0.4592811 | 0.8539432 |
| 52 | 3 | 0.6902559 | 0.10521831 | 0.4592811 | 0.8539432 |
| 91 | 3 | 0.6902559 | 0.10521831 | 0.4592811 | 0.8539432 |
| 93 | 3 | 0.6902559 | 0.10521831 | 0.4592811 | 0.8539432 |
| 84 | 3 | 0.6991368 | 0.10065312 | 0.4763471 | 0.8558282 |
| 20 | 3 | 0.6991368 | 0.10065312 | 0.4763471 | 0.8558282 |
| 13 | 4 | 0.6991368 | 0.10065312 | 0.4763471 | 0.8558282 |
| 77 | 4 | NA | NA | NA | NA |
| 82 | 4 | NA | NA | NA | NA |
| 34 | 4 | NA | NA | NA | NA |
| 105 | 4 | 0.5237279 | 0.16304345 | 0.2339502 | 0.7983648 |
| 108 | 4 | 0.5673965 | 0.13128008 | 0.3149611 | 0.7890984 |
| 31 | 4 | 0.5762773 | 0.13592943 | 0.3135554 | 0.8019567 |
| 110 | 4 | 0.5762773 | 0.13592943 | 0.3135554 | 0.8019567 |
| 103 | 4 | 0.5762773 | 0.13592943 | 0.3135554 | 0.8019567 |
| 4 | 4 | 0.5673965 | 0.13128008 | 0.3149611 | 0.7890984 |
| 19 | 4 | 0.5762773 | 0.13592943 | 0.3135554 | 0.8019567 |
| 96 | 4 | 0.5762773 | 0.13592943 | 0.3135554 | 0.8019567 |
| 74 | 4 | 0.5762773 | 0.13592943 | 0.3135554 | 0.8019567 |
| 102 | 4 | 0.5762773 | 0.13592943 | 0.3135554 | 0.8019567 |
| 76 | 4 | 0.5762773 | 0.13592943 | 0.3135554 | 0.8019567 |
| 21 | 4 | 0.5762773 | 0.13592943 | 0.3135554 | 0.8019567 |
| 37 | 4 | 0.5762773 | 0.13592943 | 0.3135554 | 0.8019567 |
| 72 | 4 | 0.5762773 | 0.13592943 | 0.3135554 | 0.8019567 |
| 14 | 4 | 0.5673965 | 0.13128008 | 0.3149611 | 0.7890984 |
| 104 | 4 | 0.5762773 | 0.13592943 | 0.3135554 | 0.8019567 |
| 17 | 4 | 0.5673965 | 0.13128008 | 0.3149611 | 0.7890984 |
| 60 | 4 | 0.5673965 | 0.13128008 | 0.3149611 | 0.7890984 |
| 52 | 4 | 0.5673965 | 0.13128008 | 0.3149611 | 0.7890984 |
| 91 | 4 | 0.5673965 | 0.13128008 | 0.3149611 | 0.7890984 |
| 93 | 4 | 0.5673965 | 0.13128008 | 0.3149611 | 0.7890984 |
| 84 | 4 | 0.5762773 | 0.13592943 | 0.3135554 | 0.8019567 |
| 20 | 4 | 0.5762773 | 0.13592943 | 0.3135554 | 0.8019567 |

Table S10. Model-averaged r^Sm^ estimates (det. prob. of *S. albiventris*, given *M. fairbanki* was not detected) and precision parameters for Two-species models (MS format).

| **Site IDs** | **Survey no.** | **Estimate** | **Standard Error** | **Lower 95% Confidence Interval** | **Upper 95% Confidence Interval** |
| --- | --- | --- | --- | --- | --- |
| 13 | 1 | 0.4389893 | 0.1380387 | 0.2068816869 | 0.7012581 |
| 77 | 1 | 0.4924549 | 0.1304260 | 0.2586615213 | 0.7295955 |
| 82 | 1 | 0.4826579 | 0.1307930 | 0.2504845522 | 0.7225683 |
| 34 | 1 | 0.4924549 | 0.1304260 | 0.2586615213 | 0.7295955 |
| 105 | 1 | 0.4924549 | 0.1304260 | 0.2586615213 | 0.7295955 |
| 108 | 1 | 0.4826579 | 0.1307930 | 0.2504845522 | 0.7225683 |
| 31 | 1 | 0.4826579 | 0.1307930 | 0.2504845522 | 0.7225683 |
| 110 | 1 | 0.4924549 | 0.1304260 | 0.2586615213 | 0.7295955 |
| 103 | 1 | 0.4924549 | 0.1304260 | 0.2586615213 | 0.7295955 |
| 4 | 1 | 0.4924549 | 0.1304260 | 0.2586615213 | 0.7295955 |
| 19 | 1 | 0.4826579 | 0.1307930 | 0.2504845522 | 0.7225683 |
| 96 | 1 | 0.4826579 | 0.1307930 | 0.2504845522 | 0.7225683 |
| 74 | 1 | 0.4826579 | 0.1307930 | 0.2504845522 | 0.7225683 |
| 102 | 1 | 0.4826579 | 0.1307930 | 0.2504845522 | 0.7225683 |
| 76 | 1 | 0.4826579 | 0.1307930 | 0.2504845522 | 0.7225683 |
| 21 | 1 | 0.4826579 | 0.1307930 | 0.2504845522 | 0.7225683 |
| 37 | 1 | 0.4924549 | 0.1304260 | 0.2586615213 | 0.7295955 |
| 72 | 1 | 0.4826579 | 0.1307930 | 0.2504845522 | 0.7225683 |
| 14 | 1 | 0.4924549 | 0.1304260 | 0.2586615213 | 0.7295955 |
| 104 | 1 | 0.4826579 | 0.1307930 | 0.2504845522 | 0.7225683 |
| 17 | 1 | 0.4924549 | 0.1304260 | 0.2586615213 | 0.7295955 |
| 60 | 1 | 0.4924549 | 0.1304260 | 0.2586615213 | 0.7295955 |
| 52 | 1 | 0.4924549 | 0.1304260 | 0.2586615213 | 0.7295955 |
| 91 | 1 | 0.4924549 | 0.1304260 | 0.2586615213 | 0.7295955 |
| 93 | 1 | 0.4924549 | 0.1304260 | 0.2586615213 | 0.7295955 |
| 84 | 1 | 0.4826579 | 0.1307930 | 0.2504845522 | 0.7225683 |
| 20 | 1 | 0.4826579 | 0.1307930 | 0.2504845522 | 0.7225683 |
| 13 | 2 | 0.4924549 | 0.1304260 | 0.2586615213 | 0.7295955 |
| 77 | 2 | 0.5365053 | 0.1384590 | 0.2798845757 | 0.7751455 |
| 82 | 2 | 0.5267083 | 0.1418798 | 0.2672607877 | 0.7724917 |
| 34 | 2 | 0.5365053 | 0.1384590 | 0.2798845757 | 0.7751455 |
| 105 | 2 | 0.5365053 | 0.1384590 | 0.2798845757 | 0.7751455 |
| 108 | 2 | 0.5365053 | 0.1384590 | 0.2798845757 | 0.7751455 |
| 31 | 2 | 0.5267083 | 0.1418798 | 0.2672607877 | 0.7724917 |
| 110 | 2 | 0.5267083 | 0.1418798 | 0.2672607877 | 0.7724917 |
| 103 | 2 | 0.5267083 | 0.1418798 | 0.2672607877 | 0.7724917 |
| 4 | 2 | 0.5267083 | 0.1418798 | 0.2672607877 | 0.7724917 |
| 19 | 2 | 0.5365053 | 0.1384590 | 0.2798845757 | 0.7751455 |
| 96 | 2 | 0.5365053 | 0.1384590 | 0.2798845757 | 0.7751455 |
| 74 | 2 | 0.5365053 | 0.1384590 | 0.2798845757 | 0.7751455 |
| 102 | 2 | 0.5267083 | 0.1418798 | 0.2672607877 | 0.7724917 |
| 76 | 2 | 0.5365053 | 0.1384590 | 0.2798845757 | 0.7751455 |
| 21 | 2 | 0.5365053 | 0.1384590 | 0.2798845757 | 0.7751455 |
| 37 | 2 | 0.5267083 | 0.1418798 | 0.2672607877 | 0.7724917 |
| 72 | 2 | 0.5365053 | 0.1384590 | 0.2798845757 | 0.7751455 |
| 14 | 2 | 0.5365053 | 0.1384590 | 0.2798845757 | 0.7751455 |
| 104 | 2 | 0.5267083 | 0.1418798 | 0.2672607877 | 0.7724917 |
| 17 | 2 | 0.5365053 | 0.1384590 | 0.2798845757 | 0.7751455 |
| 60 | 2 | 0.5365053 | 0.1384590 | 0.2798845757 | 0.7751455 |
| 52 | 2 | 0.5365053 | 0.1384590 | 0.2798845757 | 0.7751455 |
| 91 | 2 | 0.5365053 | 0.1384590 | 0.2798845757 | 0.7751455 |
| 93 | 2 | 0.5365053 | 0.1384590 | 0.2798845757 | 0.7751455 |
| 84 | 2 | 0.5267083 | 0.1418798 | 0.2672607877 | 0.7724917 |
| 20 | 2 | 0.5267083 | 0.1418798 | 0.2672607877 | 0.7724917 |
| 13 | 3 | 0.5267083 | 0.1418798 | 0.2672607877 | 0.7724917 |
| 77 | 3 | 0.5680193 | 0.1650571 | 0.2602540482 | 0.8309251 |
| 82 | 3 | 0.5778163 | 0.1596102 | 0.2751663440 | 0.8314863 |
| 34 | 3 | 0.5680193 | 0.1650571 | 0.2602540482 | 0.8309251 |
| 105 | 3 | 0.5680193 | 0.1650571 | 0.2602540482 | 0.8309251 |
| 108 | 3 | 0.5680193 | 0.1650571 | 0.2602540482 | 0.8309251 |
| 31 | 3 | 0.5778163 | 0.1596102 | 0.2751663440 | 0.8314863 |
| 110 | 3 | 0.5680193 | 0.1650571 | 0.2602540482 | 0.8309251 |
| 103 | 3 | 0.5680193 | 0.1650571 | 0.2602540482 | 0.8309251 |
| 4 | 3 | 0.5778163 | 0.1596102 | 0.2751663440 | 0.8314863 |
| 19 | 3 | 0.5680193 | 0.1650571 | 0.2602540482 | 0.8309251 |
| 96 | 3 | 0.5680193 | 0.1650571 | 0.2602540482 | 0.8309251 |
| 74 | 3 | 0.5680193 | 0.1650571 | 0.2602540482 | 0.8309251 |
| 102 | 3 | 0.5778163 | 0.1596102 | 0.2751663440 | 0.8314863 |
| 76 | 3 | 0.5680193 | 0.1650571 | 0.2602540482 | 0.8309251 |
| 21 | 3 | 0.5778163 | 0.1596102 | 0.2751663440 | 0.8314863 |
| 37 | 3 | 0.5680193 | 0.1650571 | 0.2602540482 | 0.8309251 |
| 72 | 3 | 0.5680193 | 0.1650571 | 0.2602540482 | 0.8309251 |
| 14 | 3 | 0.5680193 | 0.1650571 | 0.2602540482 | 0.8309251 |
| 104 | 3 | 0.5778163 | 0.1596102 | 0.2751663440 | 0.8314863 |
| 17 | 3 | 0.5680193 | 0.1650571 | 0.2602540482 | 0.8309251 |
| 60 | 3 | 0.5680193 | 0.1650571 | 0.2602540482 | 0.8309251 |
| 52 | 3 | 0.5680193 | 0.1650571 | 0.2602540482 | 0.8309251 |
| 91 | 3 | 0.5680193 | 0.1650571 | 0.2602540482 | 0.8309251 |
| 93 | 3 | 0.5680193 | 0.1650571 | 0.2602540482 | 0.8309251 |
| 84 | 3 | 0.5778163 | 0.1596102 | 0.2751663440 | 0.8314863 |
| 20 | 3 | 0.5778163 | 0.1596102 | 0.2751663440 | 0.8314863 |
| 13 | 4 | 0.5778163 | 0.1596102 | 0.2751663440 | 0.8314863 |
| 77 | 4 | NA | NA | NA | NA |
| 82 | 4 | NA | NA | NA | NA |
| 34 | 4 | NA | NA | NA | NA |
| 105 | 4 | NA | NA | NA | NA |
| 108 | 4 | NA | NA | NA | NA |
| 31 | 4 | NA | NA | NA | NA |
| 110 | 4 | 0.2988429 | 0.7186936 | 0.0005126738 | 0.9971843 |
| 103 | 4 | 0.3071555 | 0.7160162 | 0.0006060901 | 0.9969238 |
| 4 | 4 | 0.2880068 | 0.7164502 | 0.0004292873 | 0.9973822 |
| 19 | 4 | 0.3068092 | 0.7161244 | 0.0006020030 | 0.9969345 |
| 96 | 4 | 0.3125154 | 0.7143817 | 0.0006714969 | 0.9967588 |
| 74 | 4 | 0.3113118 | 0.7147422 | 0.0006564578 | 0.9967956 |
| 102 | 4 | 0.2971111 | 0.7192730 | 0.0004944358 | 0.9972390 |
| 76 | 4 | 0.2993711 | 0.7185182 | 0.0005183211 | 0.9971676 |
| 21 | 4 | 0.3040383 | 0.7170001 | 0.0005699166 | 0.9970210 |
| 37 | 4 | 0.3244735 | 0.7109953 | 0.0008319652 | 0.9964039 |
| 72 | 4 | 0.3144291 | 0.7138152 | 0.0006958338 | 0.9967006 |
| 14 | 4 | 0.3001294 | 0.7127245 | 0.0005544296 | 0.9969926 |
| 104 | 4 | 0.3241272 | 0.7110882 | 0.0008270383 | 0.9964139 |
| 17 | 4 | 0.3198719 | 0.7074544 | 0.0008017708 | 0.9963854 |
| 60 | 4 | 0.3016793 | 0.7122751 | 0.0005718942 | 0.9969433 |
| 52 | 4 | 0.3195256 | 0.7075383 | 0.0007969709 | 0.9963956 |
| 91 | 4 | 0.2843787 | 0.7176364 | 0.0003957452 | 0.9974992 |
| 93 | 4 | 0.3023721 | 0.7120761 | 0.0005798084 | 0.9969213 |
| 84 | 4 | 0.3144291 | 0.7138152 | 0.0006958338 | 0.9967006 |
| 20 | 4 | 0.3104373 | 0.7150070 | 0.0006456549 | 0.9968224 |

Table S11. QAICc table of single-season, single-species models: SM format: Species A = *S. albiventris*, Species B = *M. fairbanki*

| **Model** | **QAICc** | **neg2ll** | **n. par** | **warn.conv** | **DQAICc** | **mod like** | **wgt** |
| --- | --- | --- | --- | --- | --- | --- | --- |
| psi(SP+Elevation)p(SP+INT_o+survey) | 174.5292 | 218.5449 | 7 | 7.00 | 0.0000 | 1.0000 | 0.1858 |
| psi(SP+Elevation)p(SP+INT_d+survey) | 174.7960 | 218.9219 | 7 | 7.00 | 0.2668 | 0.8751 | 0.1626 |
| psi(SP+Elevation)p(SP+INT_o) | 175.3010 | 224.8575 | 6 | 7.00 | 0.7718 | 0.6798 | 0.1263 |
| psi(SP+Elevation)p(SP+INT_d) | 175.3034 | 224.8608 | 6 | 7.00 | 0.7742 | 0.6790 | 0.1262 |
| psi(SP+Lantana_camara)p(SP+INT_o+survey) | 175.5002 | 219.9172 | 7 | 7.00 | 0.9710 | 0.6154 | 0.1144 |
| psi(SP+Lantana_camara)p(SP+INT_o) | 176.2848 | 226.2479 | 6 | 3.97 | 1.7556 | 0.4157 | 0.0772 |
| psi(SP+Elevation+No_road)  p(SP+INT_d+BefNoon) | 177.6821 | 217.1989 | 8 | 7.00 | 3.1529 | 0.2067 | 0.0384 |
| psi(SP+Elevation+No_road)  p(SP+INT_o+BefNoon) | 177.7993 | 217.3645 | 8 | 7.00 | 3.2701 | 0.1949 | 0.0362 |
| psi(SP+Elevation+Lantana_camara)  p(SP+INT_d) | 177.8403 | 223.2244 | 7 | 3.20 | 3.3110 | 0.1910 | 0.0355 |
| psi(SP+Elevation+Lantana_camara)  p(SP+INT_o) | 177.8933 | 223.2994 | 7 | 2.30 | 3.3641 | 0.1860 | 0.0346 |
| psi(SP+Elevation+Area)p(SP+INT_d+BefNoon) | 178.0625 | 217.7366 | 8 | 7.00 | 3.5333 | 0.1709 | 0.0318 |
| psi(SP+Elevation+Area)p(SP+INT_o+BefNoon) | 178.1133 | 217.8083 | 8 | 7.00 | 3.5841 | 0.1666 | 0.0310 |

Table S12: Model-averaged r^S^ estimates (det. prob. of *S. albiventris*, given both species are present) and precision parameters for Two-species models (SM format).

| **Site IDs** | **Survey no.** | **Estimate** | **Standard Error** | **Lower 95% Confidence Interval** | **Upper 95% Confidence Interval** |
| --- | --- | --- | --- | --- | --- |
| 13 | 1 | 0.5217848 | 0.10935560 | 0.3160967 | 0.7203417 |
| 77 | 1 | 0.5801077 | 0.08359606 | 0.4135203 | 0.7302444 |
| 82 | 1 | 0.5678288 | 0.08117750 | 0.4072481 | 0.7153170 |
| 34 | 1 | 0.5801077 | 0.08359606 | 0.4135203 | 0.7302444 |
| 105 | 1 | 0.5801077 | 0.08359606 | 0.4135203 | 0.7302444 |
| 108 | 1 | 0.5678288 | 0.08117750 | 0.4072481 | 0.7153170 |
| 31 | 1 | 0.5678288 | 0.08117750 | 0.4072481 | 0.7153170 |
| 110 | 1 | 0.5801077 | 0.08359606 | 0.4135203 | 0.7302444 |
| 103 | 1 | 0.5801077 | 0.08359606 | 0.4135203 | 0.7302444 |
| 4 | 1 | 0.5801077 | 0.08359606 | 0.4135203 | 0.7302444 |
| 19 | 1 | 0.5678288 | 0.08117750 | 0.4072481 | 0.7153170 |
| 96 | 1 | 0.5678288 | 0.08117750 | 0.4072481 | 0.7153170 |
| 74 | 1 | 0.5678288 | 0.08117750 | 0.4072481 | 0.7153170 |
| 102 | 1 | 0.5678288 | 0.08117750 | 0.4072481 | 0.7153170 |
| 76 | 1 | 0.5678288 | 0.08117750 | 0.4072481 | 0.7153170 |
| 21 | 1 | 0.5678288 | 0.08117750 | 0.4072481 | 0.7153170 |
| 37 | 1 | 0.5801077 | 0.08359606 | 0.4135203 | 0.7302444 |
| 72 | 1 | 0.5678288 | 0.08117750 | 0.4072481 | 0.7153170 |
| 14 | 1 | 0.5801077 | 0.08359606 | 0.4135203 | 0.7302444 |
| 104 | 1 | 0.5678288 | 0.08117750 | 0.4072481 | 0.7153170 |
| 17 | 1 | 0.5801077 | 0.08359606 | 0.4135203 | 0.7302444 |
| 60 | 1 | 0.5801077 | 0.08359606 | 0.4135203 | 0.7302444 |
| 52 | 1 | 0.5801077 | 0.08359606 | 0.4135203 | 0.7302444 |
| 91 | 1 | 0.5801077 | 0.08359606 | 0.4135203 | 0.7302444 |
| 93 | 1 | 0.5801077 | 0.08359606 | 0.4135203 | 0.7302444 |
| 84 | 1 | 0.5678288 | 0.08117750 | 0.4072481 | 0.7153170 |
| 20 | 1 | 0.5678288 | 0.08117750 | 0.4072481 | 0.7153170 |
| 13 | 2 | 0.5801077 | 0.08359606 | 0.4135203 | 0.7302444 |
| 77 | 2 | 0.6243491 | 0.08006613 | 0.4598265 | 0.7644332 |
| 82 | 2 | 0.6120702 | 0.08425283 | 0.4404226 | 0.7597834 |
| 34 | 2 | 0.6243491 | 0.08006613 | 0.4598265 | 0.7644332 |
| 105 | 2 | 0.6243491 | 0.08006613 | 0.4598265 | 0.7644332 |
| 108 | 2 | 0.6243491 | 0.08006613 | 0.4598265 | 0.7644332 |
| 31 | 2 | 0.6120702 | 0.08425283 | 0.4404226 | 0.7597834 |
| 110 | 2 | 0.6120702 | 0.08425283 | 0.4404226 | 0.7597834 |
| 103 | 2 | 0.6120702 | 0.08425283 | 0.4404226 | 0.7597834 |
| 4 | 2 | 0.6120702 | 0.08425283 | 0.4404226 | 0.7597834 |
| 19 | 2 | 0.6243491 | 0.08006613 | 0.4598265 | 0.7644332 |
| 96 | 2 | 0.6243491 | 0.08006613 | 0.4598265 | 0.7644332 |
| 74 | 2 | 0.6243491 | 0.08006613 | 0.4598265 | 0.7644332 |
| 102 | 2 | 0.6120702 | 0.08425283 | 0.4404226 | 0.7597834 |
| 76 | 2 | 0.6243491 | 0.08006613 | 0.4598265 | 0.7644332 |
| 21 | 2 | 0.6243491 | 0.08006613 | 0.4598265 | 0.7644332 |
| 37 | 2 | 0.6120702 | 0.08425283 | 0.4404226 | 0.7597834 |
| 72 | 2 | 0.6243491 | 0.08006613 | 0.4598265 | 0.7644332 |
| 14 | 2 | 0.6243491 | 0.08006613 | 0.4598265 | 0.7644332 |
| 104 | 2 | 0.6120702 | 0.08425283 | 0.4404226 | 0.7597834 |
| 17 | 2 | 0.6243491 | 0.08006613 | 0.4598265 | 0.7644332 |
| 60 | 2 | 0.6243491 | 0.08006613 | 0.4598265 | 0.7644332 |
| 52 | 2 | 0.6243491 | 0.08006613 | 0.4598265 | 0.7644332 |
| 91 | 2 | 0.6243491 | 0.08006613 | 0.4598265 | 0.7644332 |
| 93 | 2 | 0.6243491 | 0.08006613 | 0.4598265 | 0.7644332 |
| 84 | 2 | 0.6120702 | 0.08425283 | 0.4404226 | 0.7597834 |
| 20 | 2 | 0.6120702 | 0.08425283 | 0.4404226 | 0.7597834 |
| 13 | 3 | 0.6120702 | 0.08425283 | 0.4404226 | 0.7597834 |
| 77 | 3 | 0.6514546 | 0.10954074 | 0.4206510 | 0.8279232 |
| 82 | 3 | 0.6637335 | 0.10170559 | 0.4469120 | 0.8282263 |
| 34 | 3 | 0.6514546 | 0.10954074 | 0.4206510 | 0.8279232 |
| 105 | 3 | 0.6514546 | 0.10954074 | 0.4206510 | 0.8279232 |
| 108 | 3 | 0.6514546 | 0.10954074 | 0.4206510 | 0.8279232 |
| 31 | 3 | 0.6637335 | 0.10170559 | 0.4469120 | 0.8282263 |
| 110 | 3 | 0.6514546 | 0.10954074 | 0.4206510 | 0.8279232 |
| 103 | 3 | 0.6514546 | 0.10954074 | 0.4206510 | 0.8279232 |
| 4 | 3 | 0.6637335 | 0.10170559 | 0.4469120 | 0.8282263 |
| 19 | 3 | 0.6514546 | 0.10954074 | 0.4206510 | 0.8279232 |
| 96 | 3 | 0.6514546 | 0.10954074 | 0.4206510 | 0.8279232 |
| 74 | 3 | 0.6514546 | 0.10954074 | 0.4206510 | 0.8279232 |
| 102 | 3 | 0.6637335 | 0.10170559 | 0.4469120 | 0.8282263 |
| 76 | 3 | 0.6514546 | 0.10954074 | 0.4206510 | 0.8279232 |
| 21 | 3 | 0.6637335 | 0.10170559 | 0.4469120 | 0.8282263 |
| 37 | 3 | 0.6514546 | 0.10954074 | 0.4206510 | 0.8279232 |
| 72 | 3 | 0.6514546 | 0.10954074 | 0.4206510 | 0.8279232 |
| 14 | 3 | 0.6514546 | 0.10954074 | 0.4206510 | 0.8279232 |
| 104 | 3 | 0.6637335 | 0.10170559 | 0.4469120 | 0.8282263 |
| 17 | 3 | 0.6514546 | 0.10954074 | 0.4206510 | 0.8279232 |
| 60 | 3 | 0.6514546 | 0.10954074 | 0.4206510 | 0.8279232 |
| 52 | 3 | 0.6514546 | 0.10954074 | 0.4206510 | 0.8279232 |
| 91 | 3 | 0.6514546 | 0.10954074 | 0.4206510 | 0.8279232 |
| 93 | 3 | 0.6514546 | 0.10954074 | 0.4206510 | 0.8279232 |
| 84 | 3 | 0.6637335 | 0.10170559 | 0.4469120 | 0.8282263 |
| 20 | 3 | 0.6637335 | 0.10170559 | 0.4469120 | 0.8282263 |
| 13 | 4 | 0.6637335 | 0.10170559 | 0.4469120 | 0.8282263 |
| 77 | 4 | NA | NA | NA | NA |
| 82 | 4 | NA | NA | NA | NA |
| 34 | 4 | NA | NA | NA | NA |
| 105 | 4 | 0.5684089 | 0.08817917 | 0.3943324 | 0.7270808 |
| 108 | 4 | 0.6125704 | 0.08293353 | 0.4435482 | 0.7582361 |
| 31 | 4 | 0.6248493 | 0.07859856 | 0.4633177 | 0.7626672 |
| 110 | 4 | 0.6248493 | 0.07859856 | 0.4633177 | 0.7626672 |
| 103 | 4 | 0.6248493 | 0.07859856 | 0.4633177 | 0.7626672 |
| 4 | 4 | 0.6125704 | 0.08293353 | 0.4435482 | 0.7582361 |
| 19 | 4 | 0.6248493 | 0.07859856 | 0.4633177 | 0.7626672 |
| 96 | 4 | 0.6248493 | 0.07859856 | 0.4633177 | 0.7626672 |
| 74 | 4 | 0.6248493 | 0.07859856 | 0.4633177 | 0.7626672 |
| 102 | 4 | 0.6248493 | 0.07859856 | 0.4633177 | 0.7626672 |
| 76 | 4 | 0.6248493 | 0.07859856 | 0.4633177 | 0.7626672 |
| 21 | 4 | 0.6248493 | 0.07859856 | 0.4633177 | 0.7626672 |
| 37 | 4 | 0.6248493 | 0.07859856 | 0.4633177 | 0.7626672 |
| 72 | 4 | 0.6248493 | 0.07859856 | 0.4633177 | 0.7626672 |
| 14 | 4 | 0.6125704 | 0.08293353 | 0.4435482 | 0.7582361 |
| 104 | 4 | 0.6248493 | 0.07859856 | 0.4633177 | 0.7626672 |
| 17 | 4 | 0.6125704 | 0.08293353 | 0.4435482 | 0.7582361 |
| 60 | 4 | 0.6125704 | 0.08293353 | 0.4435482 | 0.7582361 |
| 52 | 4 | 0.6125704 | 0.08293353 | 0.4435482 | 0.7582361 |
| 91 | 4 | 0.6125704 | 0.08293353 | 0.4435482 | 0.7582361 |
| 93 | 4 | 0.6125704 | 0.08293353 | 0.4435482 | 0.7582361 |
| 84 | 4 | 0.6248493 | 0.07859856 | 0.4633177 | 0.7626672 |
| 20 | 4 | 0.6248493 | 0.07859856 | 0.4633177 | 0.7626672 |

Table S13: Model-averaged r^MS^ estimates (det. prob. of *M. fairbanki*, given *S. albiventris* was detected) and precision parameters for Two-species models (SM format).

| **Site IDs** | **Survey no.** | **Estimate** | **Standard Error** | **Lower 95% Confidence Interval** | **Upper 95% Confidence Interval** |
| --- | --- | --- | --- | --- | --- |
| 13 | 1 | 0.6187873 | 0.10266120 | 0.4088819 | 0.7920616 |
| 77 | 1 | 0.6739053 | 0.07374709 | 0.5170353 | 0.7995743 |
| 82 | 1 | 0.6629488 | 0.07205658 | 0.5111035 | 0.7872629 |
| 34 | 1 | 0.6739053 | 0.07374709 | 0.5170353 | 0.7995743 |
| 105 | 1 | 0.6739053 | 0.07374709 | 0.5170353 | 0.7995743 |
| 108 | 1 | 0.6629488 | 0.07205658 | 0.5111035 | 0.7872629 |
| 31 | 1 | 0.6629488 | 0.07205658 | 0.5111035 | 0.7872629 |
| 110 | 1 | 0.6739053 | 0.07374709 | 0.5170353 | 0.7995743 |
| 103 | 1 | 0.6739053 | 0.07374709 | 0.5170353 | 0.7995743 |
| 4 | 1 | 0.6739053 | 0.07374709 | 0.5170353 | 0.7995743 |
| 19 | 1 | 0.6629488 | 0.07205658 | 0.5111035 | 0.7872629 |
| 96 | 1 | 0.6629488 | 0.07205658 | 0.5111035 | 0.7872629 |
| 74 | 1 | 0.6629488 | 0.07205658 | 0.5111035 | 0.7872629 |
| 102 | 1 | 0.6629488 | 0.07205658 | 0.5111035 | 0.7872629 |
| 76 | 1 | 0.6629488 | 0.07205658 | 0.5111035 | 0.7872629 |
| 21 | 1 | 0.6629488 | 0.07205658 | 0.5111035 | 0.7872629 |
| 37 | 1 | 0.6739053 | 0.07374709 | 0.5170353 | 0.7995743 |
| 72 | 1 | 0.6629488 | 0.07205658 | 0.5111035 | 0.7872629 |
| 14 | 1 | 0.6739053 | 0.07374709 | 0.5170353 | 0.7995743 |
| 104 | 1 | 0.6629488 | 0.07205658 | 0.5111035 | 0.7872629 |
| 17 | 1 | 0.6739053 | 0.07374709 | 0.5170353 | 0.7995743 |
| 60 | 1 | 0.6739053 | 0.07374709 | 0.5170353 | 0.7995743 |
| 52 | 1 | 0.6739053 | 0.07374709 | 0.5170353 | 0.7995743 |
| 91 | 1 | 0.6739053 | 0.07374709 | 0.5170353 | 0.7995743 |
| 93 | 1 | 0.6739053 | 0.07374709 | 0.5170353 | 0.7995743 |
| 84 | 1 | 0.6629488 | 0.07205658 | 0.5111035 | 0.7872629 |
| 20 | 1 | 0.6629488 | 0.07205658 | 0.5111035 | 0.7872629 |
| 13 | 2 | 0.6739053 | 0.07374709 | 0.5170353 | 0.7995743 |
| 77 | 2 | 0.7131875 | 0.06833524 | 0.5636878 | 0.8271682 |
| 82 | 2 | 0.7022310 | 0.07269121 | 0.5440341 | 0.8233622 |
| 34 | 2 | 0.7131875 | 0.06833524 | 0.5636878 | 0.8271682 |
| 105 | 2 | 0.7131875 | 0.06833524 | 0.5636878 | 0.8271682 |
| 108 | 2 | 0.7131875 | 0.06833524 | 0.5636878 | 0.8271682 |
| 31 | 2 | 0.7022310 | 0.07269121 | 0.5440341 | 0.8233622 |
| 110 | 2 | 0.7022310 | 0.07269121 | 0.5440341 | 0.8233622 |
| 103 | 2 | 0.7022310 | 0.07269121 | 0.5440341 | 0.8233622 |
| 4 | 2 | 0.7022310 | 0.07269121 | 0.5440341 | 0.8233622 |
| 19 | 2 | 0.7131875 | 0.06833524 | 0.5636878 | 0.8271682 |
| 96 | 2 | 0.7131875 | 0.06833524 | 0.5636878 | 0.8271682 |
| 74 | 2 | 0.7131875 | 0.06833524 | 0.5636878 | 0.8271682 |
| 102 | 2 | 0.7022310 | 0.07269121 | 0.5440341 | 0.8233622 |
| 76 | 2 | 0.7131875 | 0.06833524 | 0.5636878 | 0.8271682 |
| 21 | 2 | 0.7131875 | 0.06833524 | 0.5636878 | 0.8271682 |
| 37 | 2 | 0.7022310 | 0.07269121 | 0.5440341 | 0.8233622 |
| 72 | 2 | 0.7131875 | 0.06833524 | 0.5636878 | 0.8271682 |
| 14 | 2 | 0.7131875 | 0.06833524 | 0.5636878 | 0.8271682 |
| 104 | 2 | 0.7022310 | 0.07269121 | 0.5440341 | 0.8233622 |
| 17 | 2 | 0.7131875 | 0.06833524 | 0.5636878 | 0.8271682 |
| 60 | 2 | 0.7131875 | 0.06833524 | 0.5636878 | 0.8271682 |
| 52 | 2 | 0.7131875 | 0.06833524 | 0.5636878 | 0.8271682 |
| 91 | 2 | 0.7131875 | 0.06833524 | 0.5636878 | 0.8271682 |
| 93 | 2 | 0.7131875 | 0.06833524 | 0.5636878 | 0.8271682 |
| 84 | 2 | 0.7022310 | 0.07269121 | 0.5440341 | 0.8233622 |
| 20 | 2 | 0.7022310 | 0.07269121 | 0.5440341 | 0.8233622 |
| 13 | 3 | 0.7022310 | 0.07269121 | 0.5440341 | 0.8233622 |
| 77 | 3 | 0.7349698 | 0.09251243 | 0.5222673 | 0.8755386 |
| 82 | 3 | 0.7459263 | 0.08501080 | 0.5493006 | 0.8761168 |
| 34 | 3 | 0.7349698 | 0.09251243 | 0.5222673 | 0.8755386 |
| 105 | 3 | 0.7349698 | 0.09251243 | 0.5222673 | 0.8755386 |
| 108 | 3 | 0.7349698 | 0.09251243 | 0.5222673 | 0.8755386 |
| 31 | 3 | 0.7459263 | 0.08501080 | 0.5493006 | 0.8761168 |
| 110 | 3 | 0.7349698 | 0.09251243 | 0.5222673 | 0.8755386 |
| 103 | 3 | 0.7349698 | 0.09251243 | 0.5222673 | 0.8755386 |
| 4 | 3 | 0.7459263 | 0.08501080 | 0.5493006 | 0.8761168 |
| 19 | 3 | 0.7349698 | 0.09251243 | 0.5222673 | 0.8755386 |
| 96 | 3 | 0.7349698 | 0.09251243 | 0.5222673 | 0.8755386 |
| 74 | 3 | 0.7349698 | 0.09251243 | 0.5222673 | 0.8755386 |
| 102 | 3 | 0.7459263 | 0.08501080 | 0.5493006 | 0.8761168 |
| 76 | 3 | 0.7349698 | 0.09251243 | 0.5222673 | 0.8755386 |
| 21 | 3 | 0.7459263 | 0.08501080 | 0.5493006 | 0.8761168 |
| 37 | 3 | 0.7349698 | 0.09251243 | 0.5222673 | 0.8755386 |
| 72 | 3 | 0.7349698 | 0.09251243 | 0.5222673 | 0.8755386 |
| 14 | 3 | 0.7349698 | 0.09251243 | 0.5222673 | 0.8755386 |
| 104 | 3 | 0.7459263 | 0.08501080 | 0.5493006 | 0.8761168 |
| 17 | 3 | 0.7349698 | 0.09251243 | 0.5222673 | 0.8755386 |
| 60 | 3 | 0.7349698 | 0.09251243 | 0.5222673 | 0.8755386 |
| 52 | 3 | 0.7349698 | 0.09251243 | 0.5222673 | 0.8755386 |
| 91 | 3 | 0.7349698 | 0.09251243 | 0.5222673 | 0.8755386 |
| 93 | 3 | 0.7349698 | 0.09251243 | 0.5222673 | 0.8755386 |
| 84 | 3 | 0.7459263 | 0.08501080 | 0.5493006 | 0.8761168 |
| 20 | 3 | 0.7459263 | 0.08501080 | 0.5493006 | 0.8761168 |
| 13 | 4 | 0.7459263 | 0.08501080 | 0.5493006 | 0.8761168 |
| 77 | 4 | NA | NA | NA | NA |
| 82 | 4 | NA | NA | NA | NA |
| 34 | 4 | NA | NA | NA | NA |
| 105 | 4 | 0.6087046 | 0.11840312 | 0.3699466 | 0.8047399 |
| 108 | 4 | 0.6532890 | 0.08829235 | 0.4674261 | 0.8017927 |
| 31 | 4 | 0.6642455 | 0.09084986 | 0.4709998 | 0.8146743 |
| 110 | 4 | 0.6642455 | 0.09084986 | 0.4709998 | 0.8146743 |
| 103 | 4 | 0.6642455 | 0.09084986 | 0.4709998 | 0.8146743 |
| 4 | 4 | 0.6532890 | 0.08829235 | 0.4674261 | 0.8017927 |
| 19 | 4 | 0.6642455 | 0.09084986 | 0.4709998 | 0.8146743 |
| 96 | 4 | 0.6642455 | 0.09084986 | 0.4709998 | 0.8146743 |
| 74 | 4 | 0.6642455 | 0.09084986 | 0.4709998 | 0.8146743 |
| 102 | 4 | 0.6642455 | 0.09084986 | 0.4709998 | 0.8146743 |
| 76 | 4 | 0.6642455 | 0.09084986 | 0.4709998 | 0.8146743 |
| 21 | 4 | 0.6642455 | 0.09084986 | 0.4709998 | 0.8146743 |
| 37 | 4 | 0.6642455 | 0.09084986 | 0.4709998 | 0.8146743 |
| 72 | 4 | 0.6642455 | 0.09084986 | 0.4709998 | 0.8146743 |
| 14 | 4 | 0.6532890 | 0.08829235 | 0.4674261 | 0.8017927 |
| 104 | 4 | 0.6642455 | 0.09084986 | 0.4709998 | 0.8146743 |
| 17 | 4 | 0.6532890 | 0.08829235 | 0.4674261 | 0.8017927 |
| 60 | 4 | 0.6532890 | 0.08829235 | 0.4674261 | 0.8017927 |
| 52 | 4 | 0.6532890 | 0.08829235 | 0.4674261 | 0.8017927 |
| 91 | 4 | 0.6532890 | 0.08829235 | 0.4674261 | 0.8017927 |
| 93 | 4 | 0.6532890 | 0.08829235 | 0.4674261 | 0.8017927 |
| 84 | 4 | 0.6642455 | 0.09084986 | 0.4709998 | 0.8146743 |
| 20 | 4 | 0.6642455 | 0.09084986 | 0.4709998 | 0.8146743 |

Table S14: Model-averaged r^Ms^ estimates (det. prob. of *M. fairbanki*, given *S. albiventris* was not detected) and precision parameters for Two-species models (SM format).

| **Site IDs** | **Survey no.** | **Estimate** | **Standard Error** | **Lower 95% Confidence Interval** | **Upper 95% Confidence Interval** |
| --- | --- | --- | --- | --- | --- |
| 13 | 1 | 0.5850134 | 0.1244445 | 0.340451443 | 0.7938123 |
| 77 | 1 | 0.6412636 | 0.1048587 | 0.422495837 | 0.8137013 |
| 82 | 1 | 0.6295977 | 0.1066546 | 0.409535508 | 0.8064130 |
| 34 | 1 | 0.6412636 | 0.1048587 | 0.422495837 | 0.8137013 |
| 105 | 1 | 0.6412636 | 0.1048587 | 0.422495837 | 0.8137013 |
| 108 | 1 | 0.6295977 | 0.1066546 | 0.409535508 | 0.8064130 |
| 31 | 1 | 0.6295977 | 0.1066546 | 0.409535508 | 0.8064130 |
| 110 | 1 | 0.6412636 | 0.1048587 | 0.422495837 | 0.8137013 |
| 103 | 1 | 0.6412636 | 0.1048587 | 0.422495837 | 0.8137013 |
| 4 | 1 | 0.6412636 | 0.1048587 | 0.422495837 | 0.8137013 |
| 19 | 1 | 0.6295977 | 0.1066546 | 0.409535508 | 0.8064130 |
| 96 | 1 | 0.6295977 | 0.1066546 | 0.409535508 | 0.8064130 |
| 74 | 1 | 0.6295977 | 0.1066546 | 0.409535508 | 0.8064130 |
| 102 | 1 | 0.6295977 | 0.1066546 | 0.409535508 | 0.8064130 |
| 76 | 1 | 0.6295977 | 0.1066546 | 0.409535508 | 0.8064130 |
| 21 | 1 | 0.6295977 | 0.1066546 | 0.409535508 | 0.8064130 |
| 37 | 1 | 0.6412636 | 0.1048587 | 0.422495837 | 0.8137013 |
| 72 | 1 | 0.6295977 | 0.1066546 | 0.409535508 | 0.8064130 |
| 14 | 1 | 0.6412636 | 0.1048587 | 0.422495837 | 0.8137013 |
| 104 | 1 | 0.6295977 | 0.1066546 | 0.409535508 | 0.8064130 |
| 17 | 1 | 0.6412636 | 0.1048587 | 0.422495837 | 0.8137013 |
| 60 | 1 | 0.6412636 | 0.1048587 | 0.422495837 | 0.8137013 |
| 52 | 1 | 0.6412636 | 0.1048587 | 0.422495837 | 0.8137013 |
| 91 | 1 | 0.6412636 | 0.1048587 | 0.422495837 | 0.8137013 |
| 93 | 1 | 0.6412636 | 0.1048587 | 0.422495837 | 0.8137013 |
| 84 | 1 | 0.6295977 | 0.1066546 | 0.409535508 | 0.8064130 |
| 20 | 1 | 0.6295977 | 0.1066546 | 0.409535508 | 0.8064130 |
| 13 | 2 | 0.6412636 | 0.1048587 | 0.422495837 | 0.8137013 |
| 77 | 2 | 0.6816191 | 0.1051168 | 0.453103619 | 0.8469120 |
| 82 | 2 | 0.6699532 | 0.1112248 | 0.430963880 | 0.8447320 |
| 34 | 2 | 0.6816191 | 0.1051168 | 0.453103619 | 0.8469120 |
| 105 | 2 | 0.6816191 | 0.1051168 | 0.453103619 | 0.8469120 |
| 108 | 2 | 0.6816191 | 0.1051168 | 0.453103619 | 0.8469120 |
| 31 | 2 | 0.6699532 | 0.1112248 | 0.430963880 | 0.8447320 |
| 110 | 2 | 0.6699532 | 0.1112248 | 0.430963880 | 0.8447320 |
| 103 | 2 | 0.6699532 | 0.1112248 | 0.430963880 | 0.8447320 |
| 4 | 2 | 0.6699532 | 0.1112248 | 0.430963880 | 0.8447320 |
| 19 | 2 | 0.6816191 | 0.1051168 | 0.453103619 | 0.8469120 |
| 96 | 2 | 0.6816191 | 0.1051168 | 0.453103619 | 0.8469120 |
| 74 | 2 | 0.6816191 | 0.1051168 | 0.453103619 | 0.8469120 |
| 102 | 2 | 0.6699532 | 0.1112248 | 0.430963880 | 0.8447320 |
| 76 | 2 | 0.6816191 | 0.1051168 | 0.453103619 | 0.8469120 |
| 21 | 2 | 0.6816191 | 0.1051168 | 0.453103619 | 0.8469120 |
| 37 | 2 | 0.6699532 | 0.1112248 | 0.430963880 | 0.8447320 |
| 72 | 2 | 0.6816191 | 0.1051168 | 0.453103619 | 0.8469120 |
| 14 | 2 | 0.6816191 | 0.1051168 | 0.453103619 | 0.8469120 |
| 104 | 2 | 0.6699532 | 0.1112248 | 0.430963880 | 0.8447320 |
| 17 | 2 | 0.6816191 | 0.1051168 | 0.453103619 | 0.8469120 |
| 60 | 2 | 0.6816191 | 0.1051168 | 0.453103619 | 0.8469120 |
| 52 | 2 | 0.6816191 | 0.1051168 | 0.453103619 | 0.8469120 |
| 91 | 2 | 0.6816191 | 0.1051168 | 0.453103619 | 0.8469120 |
| 93 | 2 | 0.6816191 | 0.1051168 | 0.453103619 | 0.8469120 |
| 84 | 2 | 0.6699532 | 0.1112248 | 0.430963880 | 0.8447320 |
| 20 | 2 | 0.6699532 | 0.1112248 | 0.430963880 | 0.8447320 |
| 13 | 3 | 0.6699532 | 0.1112248 | 0.430963880 | 0.8447320 |
| 77 | 3 | 0.7041232 | 0.1282143 | 0.416002479 | 0.8882735 |
| 82 | 3 | 0.7157891 | 0.1196672 | 0.442938104 | 0.8886063 |
| 34 | 3 | 0.7041232 | 0.1282143 | 0.416002479 | 0.8882735 |
| 105 | 3 | 0.7041232 | 0.1282143 | 0.416002479 | 0.8882735 |
| 108 | 3 | 0.7041232 | 0.1282143 | 0.416002479 | 0.8882735 |
| 31 | 3 | 0.7157891 | 0.1196672 | 0.442938104 | 0.8886063 |
| 110 | 3 | 0.7041232 | 0.1282143 | 0.416002479 | 0.8882735 |
| 103 | 3 | 0.7041232 | 0.1282143 | 0.416002479 | 0.8882735 |
| 4 | 3 | 0.7157891 | 0.1196672 | 0.442938104 | 0.8886063 |
| 19 | 3 | 0.7041232 | 0.1282143 | 0.416002479 | 0.8882735 |
| 96 | 3 | 0.7041232 | 0.1282143 | 0.416002479 | 0.8882735 |
| 74 | 3 | 0.7041232 | 0.1282143 | 0.416002479 | 0.8882735 |
| 102 | 3 | 0.7157891 | 0.1196672 | 0.442938104 | 0.8886063 |
| 76 | 3 | 0.7041232 | 0.1282143 | 0.416002479 | 0.8882735 |
| 21 | 3 | 0.7157891 | 0.1196672 | 0.442938104 | 0.8886063 |
| 37 | 3 | 0.7041232 | 0.1282143 | 0.416002479 | 0.8882735 |
| 72 | 3 | 0.7041232 | 0.1282143 | 0.416002479 | 0.8882735 |
| 14 | 3 | 0.7041232 | 0.1282143 | 0.416002479 | 0.8882735 |
| 104 | 3 | 0.7157891 | 0.1196672 | 0.442938104 | 0.8886063 |
| 17 | 3 | 0.7041232 | 0.1282143 | 0.416002479 | 0.8882735 |
| 60 | 3 | 0.7041232 | 0.1282143 | 0.416002479 | 0.8882735 |
| 52 | 3 | 0.7041232 | 0.1282143 | 0.416002479 | 0.8882735 |
| 91 | 3 | 0.7041232 | 0.1282143 | 0.416002479 | 0.8882735 |
| 93 | 3 | 0.7041232 | 0.1282143 | 0.416002479 | 0.8882735 |
| 84 | 3 | 0.7157891 | 0.1196672 | 0.442938104 | 0.8886063 |
| 20 | 3 | 0.7157891 | 0.1196672 | 0.442938104 | 0.8886063 |
| 13 | 4 | 0.7157891 | 0.1196672 | 0.442938104 | 0.8886063 |
| 77 | 4 | NA | NA | NA | NA |
| 82 | 4 | NA | NA | NA | NA |
| 34 | 4 | NA | NA | NA | NA |
| 105 | 4 | NA | NA | NA | NA |
| 108 | 4 | NA | NA | NA | NA |
| 31 | 4 | NA | NA | NA | NA |
| 110 | 4 | 0.3822254 | 0.7448109 | 0.001276528 | 0.9966722 |
| 103 | 4 | 0.3905884 | 0.7412730 | 0.001430096 | 0.9965258 |
| 4 | 4 | 0.3695142 | 0.7414381 | 0.001144219 | 0.9966761 |
| 19 | 4 | 0.3902399 | 0.7414173 | 0.001423526 | 0.9965316 |
| 96 | 4 | 0.3960010 | 0.7390688 | 0.001533973 | 0.9964386 |
| 74 | 4 | 0.3947698 | 0.7395640 | 0.001510050 | 0.9964579 |
| 102 | 4 | 0.3804832 | 0.7455679 | 0.001245644 | 0.9967044 |
| 76 | 4 | 0.3827365 | 0.7445900 | 0.001285664 | 0.9966628 |
| 21 | 4 | 0.3874523 | 0.7425811 | 0.001371495 | 0.9965791 |
| 37 | 4 | 0.4080111 | 0.7344198 | 0.001775931 | 0.9962687 |
| 72 | 4 | 0.3979059 | 0.7383088 | 0.001571331 | 0.9964095 |
| 14 | 4 | 0.3817101 | 0.7364696 | 0.001360649 | 0.9964379 |
| 104 | 4 | 0.4076627 | 0.7345499 | 0.001768705 | 0.9962732 |
| 17 | 4 | 0.4015720 | 0.7291223 | 0.001751413 | 0.9961189 |
| 60 | 4 | 0.3832898 | 0.7358515 | 0.001390065 | 0.9964093 |
| 52 | 4 | 0.4012236 | 0.7292431 | 0.001744195 | 0.9961237 |
| 91 | 4 | 0.3658438 | 0.7430000 | 0.001082947 | 0.9967531 |
| 93 | 4 | 0.3839867 | 0.7355806 | 0.001403141 | 0.9963968 |
| 84 | 4 | 0.3979059 | 0.7383088 | 0.001571331 | 0.9964095 |
| 20 | 4 | 0.3939103 | 0.7399123 | 0.001493445 | 0.9964716 |

Table S15. The counts and weights of covariates influencing (a) psi and (b) p in the models of the QAICc table of *M. fairbanki*.

| **Covariates** | **Count** | **Sum weight** |
| --- | --- | --- |
| Warm | 2 | 0.0161 |
| survey | 33 | 0.3118 |
| Before noon | 22 | 0.1965 |
| Cool | 7 | 0.0331 |
| Windy | 38 | 0.412 |
| Overcast | 3 | 0.0173 |
| After noon | 10 | 0.0814 |
| Clear sky | 8 | 0.0588 |
| Sunny | 1 | 0.01 |
| Neutral | 1 | 0.0097 |
| Visibility | 11 | 0.0972 |

(a)	(b)

| **Covariates** | **Count** | **Sum weight** |
| --- | --- | --- |
| Area | 7 | 0.0503 |
| Aspect | 4 | 0.0256 |
| Burn | 1 | 0.0102 |
| Canopy height | 1 | 0.0113 |
| Dry lotic | 5 | 0.0498 |
| Dirt road | 4 | 0.0721 |
| Elevation | 9 | 0.0741 |
| Lentic | 1 | 0.0109 |
| Metalled road | 1 | 0.0098 |
| No road | 12 | 0.2856 |
| *Rubus niveus* | 4 | 0.0264 |
| Slope | 6 | 0.066 |
| Understorey height | 2 | 0.0247 |
| Visibility | 11 | 0.0972 |
| Wet lotic | 1 | 0.0108 |

Table S16. The counts and weights of covariates influencing (a) psi and (b) p in the models of the QAICc table of *S. albiventris*.

(a)	(b)

| **Covariates** | **Count** | **Sum weight** |
| --- | --- | --- |
| Elevation | 8 | 0.7835 |
| No road | 1 | 0.0451 |
| *Lantana camara* | 3 | 0.2619 |
| Area | 1 | 0.0372 |

| **Covariates** | **Count** | **Sum weight** |
| --- | --- | --- |
| Before noon | 3 | 0.1174 |

Table S17. The counts and weights of covariates influencing (a) psi and (b) p in the models of the QAICc table of single-season, Two-species models: MS format: Species A = *M. fairbanki*, Species B = *S. albiventris*.

(a)	(b)

| **Covariates** | **Count** | **Sum weight** |
| --- | --- | --- |
| Elevation | 7 | 0.6943 |
| *Lantana camara* | 4 | 0.2941 |
| Understorey height | 1 | 0.0117 |

| **Covariates** | **Count** | **Sum weight** |
| --- | --- | --- |
| Neutral | 8 | 0.7722 |
| survey | 6 | 0.4369 |
| Warm | 2 | 0.0711 |
| Cool | 2 | 0.0524 |

Table S18. The counts and weights of covariates influencing (a) psi and (b) p in the models of the QAICc table of single-season, Two-species models: SM format: Species A = *S. albiventris*, Species B = *M. fairbanki*.

(a)	(b)

| **Covariates** | **Count** | **Sum weight** |
| --- | --- | --- |
| Before noon | 4 | 0.1374 |

| **Covariates** | **Count** | **Sum weight** |
| --- | --- | --- |
| Elevation | 10 | 0.8084 |
| No_road | 2 | 0.0746 |
| *Lantana_camara* | 4 | 0.2617 |
| Area | 2 | 0.0628 |

Table S19. Simulated psi and p estimates and associated precision parameters of *M. fairbanki* under varying conditions of no. of sites and number of surveys.

| **No. of Sites** | **No. of surveys** | **Estimate (psi)** | **Standard Error (psi)** | **Lower 95% Confidence Interval (psi)** | **Upper 95% Confidence Interval (psi)** | **Estimate (p)** | **Standard Error (p)** | **Lower 95% Confidence Interval (p)** | **Upper 95% Confidence Interval (p)** |
| --- | --- | --- | --- | --- | --- | --- | --- | --- | --- |
| 27 | 2 | 0.861815 | 0.20007417 | 0.3515051 | 1.0000000 | 0.6553877 | 0.19247433 | 0.3614891 | 0.8959096 |
| 29 | 2 | 0.861815 | 0.19305189 | 0.3635594 | 1.0000000 | 0.6553877 | 0.18571869 | 0.3676021 | 0.8909299 |
| 31 | 2 | 0.861815 | 0.18672058 | 0.3744271 | 1.0000000 | 0.6553877 | 0.17962797 | 0.3731288 | 0.8863579 |
| 33 | 2 | 0.861815 | 0.18097392 | 0.3842739 | 1.0000000 | 0.6553877 | 0.17409973 | 0.3781560 | 0.8821403 |
| 35 | 2 | 0.861815 | 0.17572723 | 0.3932368 | 0.9989881 | 0.6553877 | 0.16905217 | 0.3827538 | 0.8782330 |
| 37 | 2 | 0.861815 | 0.17091187 | 0.4014306 | 0.9976160 | 0.6553877 | 0.16441991 | 0.3869787 | 0.8745995 |
| 39 | 2 | 0.861815 | 0.16647185 | 0.4089508 | 0.9962977 | 0.6553877 | 0.16014832 | 0.3908785 | 0.8712085 |
| 41 | 2 | 0.861815 | 0.16236071 | 0.4158785 | 0.9950300 | 0.6553877 | 0.15619348 | 0.3944917 | 0.8680343 |
| 43 | 2 | 0.861815 | 0.15854002 | 0.4222816 | 0.9938099 | 0.6553877 | 0.15251779 | 0.3978516 | 0.8650543 |
| 27 | 3 | 0.861815 | 0.13706434 | 0.4574648 | 0.9861360 | 0.6553877 | 0.12614942 | 0.4219622 | 0.8428333 |
| 29 | 3 | 0.861815 | 0.13225368 | 0.4651177 | 0.9842113 | 0.6553877 | 0.12172163 | 0.4260049 | 0.8389571 |
| 31 | 3 | 0.861815 | 0.12791625 | 0.4719344 | 0.9824060 | 0.6553877 | 0.11773032 | 0.4296459 | 0.8354276 |
| 33 | 3 | 0.861815 | 0.12397944 | 0.4780493 | 0.9807082 | 0.6553877 | 0.11410652 | 0.4329484 | 0.8321939 |
| 35 | 3 | 0.861815 | 0.12038509 | 0.4835696 | 0.9791072 | 0.6553877 | 0.11079809 | 0.4359605 | 0.8292176 |
| 37 | 3 | 0.861815 | 0.11708617 | 0.4885816 | 0.9775939 | 0.6553877 | 0.10776273 | 0.4387211 | 0.8264668 |
| 39 | 3 | 0.861815 | 0.11404438 | 0.4931552 | 0.9761604 | 0.6553877 | 0.10496266 | 0.4412651 | 0.8239121 |
| 41 | 3 | 0.861815 | 0.11122817 | 0.4973478 | 0.9747999 | 0.6553877 | 0.10237084 | 0.4436175 | 0.8215329 |
| 43 | 3 | 0.861815 | 0.10861066 | 0.5012076 | 0.9735059 | 0.6553877 | 0.09996169 | 0.4458018 | 0.8193088 |
| 27 | 4 | 0.861815 | 0.12202870 | 0.4810529 | 0.9798454 | 0.6553877 | 0.10065900 | 0.4451698 | 0.8199538 |
| 29 | 4 | 0.861815 | 0.11774581 | 0.4875837 | 0.9778999 | 0.6553877 | 0.09712630 | 0.4483697 | 0.8166759 |
| 31 | 4 | 0.861815 | 0.11388421 | 0.4933948 | 0.9760839 | 0.6553877 | 0.09394028 | 0.4512511 | 0.8136975 |
| 33 | 4 | 0.861815 | 0.11037915 | 0.4986037 | 0.9743833 | 0.6553877 | 0.09104978 | 0.4538614 | 0.8109773 |
| 35 | 4 | 0.861815 | 0.10717914 | 0.5033033 | 0.9727861 | 0.6553877 | 0.08840972 | 0.4562420 | 0.8084778 |
| 37 | 4 | 0.861815 | 0.10424214 | 0.5075685 | 0.9712817 | 0.6553877 | 0.08598690 | 0.4584238 | 0.8061715 |
| 39 | 4 | 0.861815 | 0.10153415 | 0.5114595 | 0.9698614 | 0.6553877 | 0.08375370 | 0.4604321 | 0.8040351 |
| 41 | 4 | 0.861815 | 0.09902673 | 0.5150261 | 0.9685173 | 0.6553877 | 0.08168502 | 0.4622901 | 0.8020471 |
| 43 | 4 | 0.861815 | 0.09669623 | 0.5183094 | 0.9672425 | 0.6553877 | 0.07976272 | 0.4640145 | 0.8001920 |

Table S20. Simulated psi and p estimates and associated precision parameters of *S. albiventris* under varying conditions of no. of sites and number of surveys.

| **No. of Sites** | **No. of surveys** | **Estimate (psi)** | **Standard Error (psi)** | **Lower 95% Confidence Interval (psi)** | **Upper 95% Confidence Interval (psi)** | **Estimate (p)** | **Standard Error (p)** | **Lower 95% Confidence Interval (p)** | **Upper 95% Confidence Interval (p)** |
| --- | --- | --- | --- | --- | --- | --- | --- | --- | --- |
| 27 | 2 | 0.4543001 | 0.1752448 | 0.1086455 | 0.7821791 | 0.5730774 | 0.2717967 | 0.2020831 | 0.9177764 |
| 29 | 2 | 0.4543001 | 0.1690940 | 0.1118747 | 0.7735290 | 0.5730774 | 0.2622573 | 0.2086361 | 0.9105941 |
| 31 | 2 | 0.4543001 | 0.1635487 | 0.1148435 | 0.7656166 | 0.5730774 | 0.2536561 | 0.2146531 | 0.9039414 |
| 33 | 2 | 0.4543001 | 0.1585152 | 0.1175853 | 0.7583432 | 0.5730774 | 0.2458493 | 0.2202015 | 0.8977573 |
| 35 | 2 | 0.4543001 | 0.1539195 | 0.1201275 | 0.7516283 | 0.5730774 | 0.2387218 | 0.2253380 | 0.8919892 |
| 37 | 2 | 0.4543001 | 0.1497018 | 0.1224931 | 0.7454046 | 0.5730774 | 0.2321804 | 0.2301103 | 0.8865928 |
| 39 | 2 | 0.4543001 | 0.1458125 | 0.1247019 | 0.7396150 | 0.5730774 | 0.2261488 | 0.2345588 | 0.8815296 |
| 41 | 2 | 0.4543001 | 0.1422117 | 0.1267702 | 0.7342126 | 0.5730774 | 0.2205639 | 0.2387183 | 0.8767665 |
| 43 | 2 | 0.4543001 | 0.1388653 | 0.1287123 | 0.7291561 | 0.5730774 | 0.2153732 | 0.2426181 | 0.8722752 |
| 27 | 3 | 0.4543001 | 0.1370851 | 0.1297532 | 0.7264525 | 0.5730774 | 0.1730882 | 0.2755081 | 0.8333827 |
| 29 | 3 | 0.4543001 | 0.1322733 | 0.1325937 | 0.7190979 | 0.5730774 | 0.1670129 | 0.2803813 | 0.8274612 |
| 31 | 3 | 0.4543001 | 0.1279354 | 0.1351877 | 0.7124108 | 0.5730774 | 0.1615359 | 0.2848026 | 0.8220523 |
| 33 | 3 | 0.4543001 | 0.1239980 | 0.1375694 | 0.7062956 | 0.5730774 | 0.1565642 | 0.2888378 | 0.8170850 |
| 35 | 3 | 0.4543001 | 0.1204029 | 0.1397662 | 0.7006757 | 0.5730774 | 0.1520250 | 0.2925394 | 0.8125023 |
| 37 | 3 | 0.4543001 | 0.1171039 | 0.1418006 | 0.6954888 | 0.5730774 | 0.1478594 | 0.2959504 | 0.8082573 |
| 39 | 3 | 0.4543001 | 0.1140617 | 0.1436921 | 0.6906814 | 0.5730774 | 0.1440182 | 0.2991071 | 0.8043096 |
| 41 | 3 | 0.4543001 | 0.1112449 | 0.1454566 | 0.6862098 | 0.5730774 | 0.1404614 | 0.3020393 | 0.8006260 |
| 43 | 3 | 0.4543001 | 0.1086269 | 0.1471079 | 0.6820368 | 0.5730774 | 0.1371561 | 0.3047720 | 0.7971787 |
| 27 | 4 | 0.4543001 | 0.1280700 | 0.1351068 | 0.7126190 | 0.5730774 | 0.1348734 | 0.3066634 | 0.7947845 |
| 29 | 4 | 0.4543001 | 0.1235748 | 0.1378269 | 0.7056358 | 0.5730774 | 0.1301395 | 0.3105962 | 0.7897844 |
| 31 | 4 | 0.4543001 | 0.1195219 | 0.1403078 | 0.6992933 | 0.5730774 | 0.1258714 | 0.3141535 | 0.7852365 |
| 33 | 4 | 0.4543001 | 0.1158434 | 0.1425826 | 0.6934997 | 0.5730774 | 0.1219976 | 0.3173909 | 0.7810763 |
| 35 | 4 | 0.4543001 | 0.1124849 | 0.1446783 | 0.6881807 | 0.5730774 | 0.1184607 | 0.3203538 | 0.7772512 |
| 37 | 4 | 0.4543001 | 0.1094029 | 0.1466173 | 0.6832753 | 0.5730774 | 0.1152143 | 0.3230787 | 0.7737182 |
| 39 | 4 | 0.4543001 | 0.1065605 | 0.1484187 | 0.6787316 | 0.5730774 | 0.1122218 | 0.3255949 | 0.7704429 |
| 41 | 4 | 0.4543001 | 0.1039288 | 0.1500977 | 0.6745085 | 0.5730774 | 0.1094503 | 0.3279288 | 0.7673938 |
| 43 | 4 | 0.4543001 | 0.1014832 | 0.1516674 | 0.6705702 | 0.5730774 | 0.1068745 | 0.3301008 | 0.7645466 |

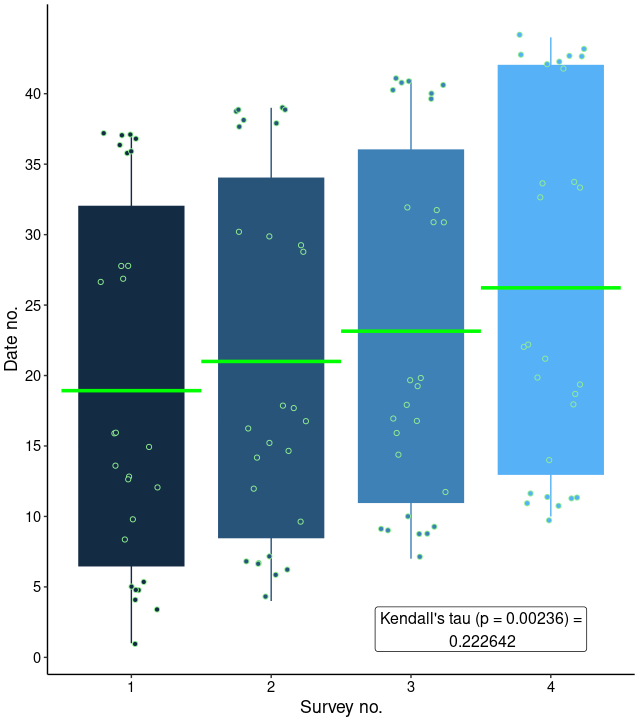

Figure S12: Correlation between survey numbers and date numbers.
